# Engineered microRNA feedback circuits enable tunable and autonomous control of synthetic receptor activity

**DOI:** 10.64898/2026.05.22.727153

**Authors:** Bryan L. Nathalia, Anna-Maria Makri-Pistikou, Theodorus J. Meuleman, Wilco Nijenhuis, Derin R. Gumustop, Eduardo González-García, Kasey S. Love, Werner Doensen, Anton F. van Galen, Kate E. Galloway, Lukas C. Kapitein, Tom F.A. de Greef

## Abstract

Synthetic receptors are powerful tools for cellular engineering, yet their utility is often constrained by constitutive activity and the absence of natural feedback mechanisms that maintain cellular homeostasis. To address this, we engineered an autonomous regulatory circuit for synthetic Notch (synNotch) receptor activation using an orthogonal microRNA (miRNA)-mediated negative feedback mechanism in mammalian cells. Specifically, synNotch activation induces the expression of a synthetic miRNA that targets complementary sites engineered into the synNotch transcript, resulting in a self-limiting feedback loop. We demonstrate that synNotch repression positively correlates with the number of miRNA target sites integrated into the receptor transcript, enabling tunable control of receptor expression. Furthermore, flow cytometry and live-cell imaging analyses revealed effective suppression of synNotch expression following activation. Downstream target gene expression peaked at approximately 48 hours post-activation before gradually declining as the feedback circuit engaged. This approach achieves autonomous synthetic receptor regulation—eliminating the need for external intervention—and enables tunable response dynamics through rational circuit design. Our miRNA feedback strategy provides a generalizable platform for engineering self-regulating synthetic receptor systems with improved control and predictability for cellular engineering applications.

## Introduction

Synthetic receptors, designed to emulate and augment natural signalling processes, are engineered molecular systems that transduce extracellular cues into programmable intracellular responses^1^. Their development represents a pivotal advance in mammalian cell engineering and has opened new avenues for the field of cell-based therapeutics^2–4^. Decades of synthetic receptor engineering dedicated to therapeutic applications have culminated in the regulatory approval and successful commercialization of several chimeric antigen receptor (CAR)-T cell therapies^5–7^, which demonstrate remarkable therapeutic benefit in the treatment of haematological malignancies^8–10^. Despite these advances, synthetic receptor therapies remain constrained by significant safety and efficacy challenges. Constitutive or excessive synthetic receptor activation can induce severe toxicities, including cytokine release syndrome and neurotoxicity, while also promoting premature clearance of therapeutic cells, thereby reducing persistence^11–14^. To mitigate these risks, several strategies for temporal control of receptor activation have been developed. These strategies utilize universal signalling mediators, including intermediary antibody switches^15–17^ and leucine zippers^18^, as well as small molecules that enable titratable pharmacologic regulation^19–22^. While these approaches allow external modulation and tunability of receptor activity, they depend upon exogenous inputs and consequently lack direct feedback mechanisms responsive to the presence of the targeted antigen. Importantly, this limitation is not unique to CAR-T cells, but represents a fundamental challenge for synthetic receptor platforms more broadly, as any receptor design lacking intrinsic regulatory mechanisms remains vulnerable to uncontrolled or sustained activation. A conceptual solution to this challenge would involve designing synthetic receptor circuits in which activation of a synthetic receptor not only triggers target gene expression but also induces regulatory elements that feed back to repress receptor activity (Fig. 1a). This design would ensure signal-dependent self-limitation, thereby mitigating the risk of sustained overactivation. Nature provides multiple precedents for this design principle, most prominently through microRNA (miRNA)-mediated feedback loops, wherein receptor activation induces expression of a miRNA that targets the receptor messenger RNA (mRNA)^23–27^. This mechanism results in a regulatory circuit that modulates receptor activity through post-transcriptional control and contributes to homeostatic maintenance. Canonical Notch signalling in neuronal cells exemplifies this principle, as receptor activity undergoes modulation through a negative feedback loop involving miR-9^28,29^. Beyond their natural function, miRNAs possess properties that make them suitable regulators for synthetic implementation: they are genetically compact, easily programmable, and can be designed to function in an orthogonal manner, independent of endogenous pathways^30–32^. Numerous studies have already established miRNA-based regulatory circuits in mammalian cells, where endogenous or synthetic miRNA inputs have been harnessed to implement Boolean logic operations as well as support more complex numerical operations and capture miRNA competition dynamics^33–37^. More recently, compact and orthogonal synthetic miRNA toolkits have expanded these applications, enabling precise, tunable and dosage-invariant control of gene expression across diverse mammalian cell types^38–40^, further underscoring the versatility of miRNAs for synthetic biology. However, to date, no synthetic receptor platform has harnessed the unique utility of miRNAs to establish intrinsic activation-induced miRNA-mediated feedback for autonomous regulation of receptor activity. We therefore sought to implement a comparable feedback strategy in a synthetic receptor system, focusing on synthetic Notch (synNotch)^41,42^, a modular platform for coupling ligand recognition to user-defined transcriptional outputs. Despite their modularity and widespread application^43–46^, synNotch receptors, like CARs, lack intrinsic regulatory mechanisms and remain prone to sustained activation once triggered.

**Figure 1:**
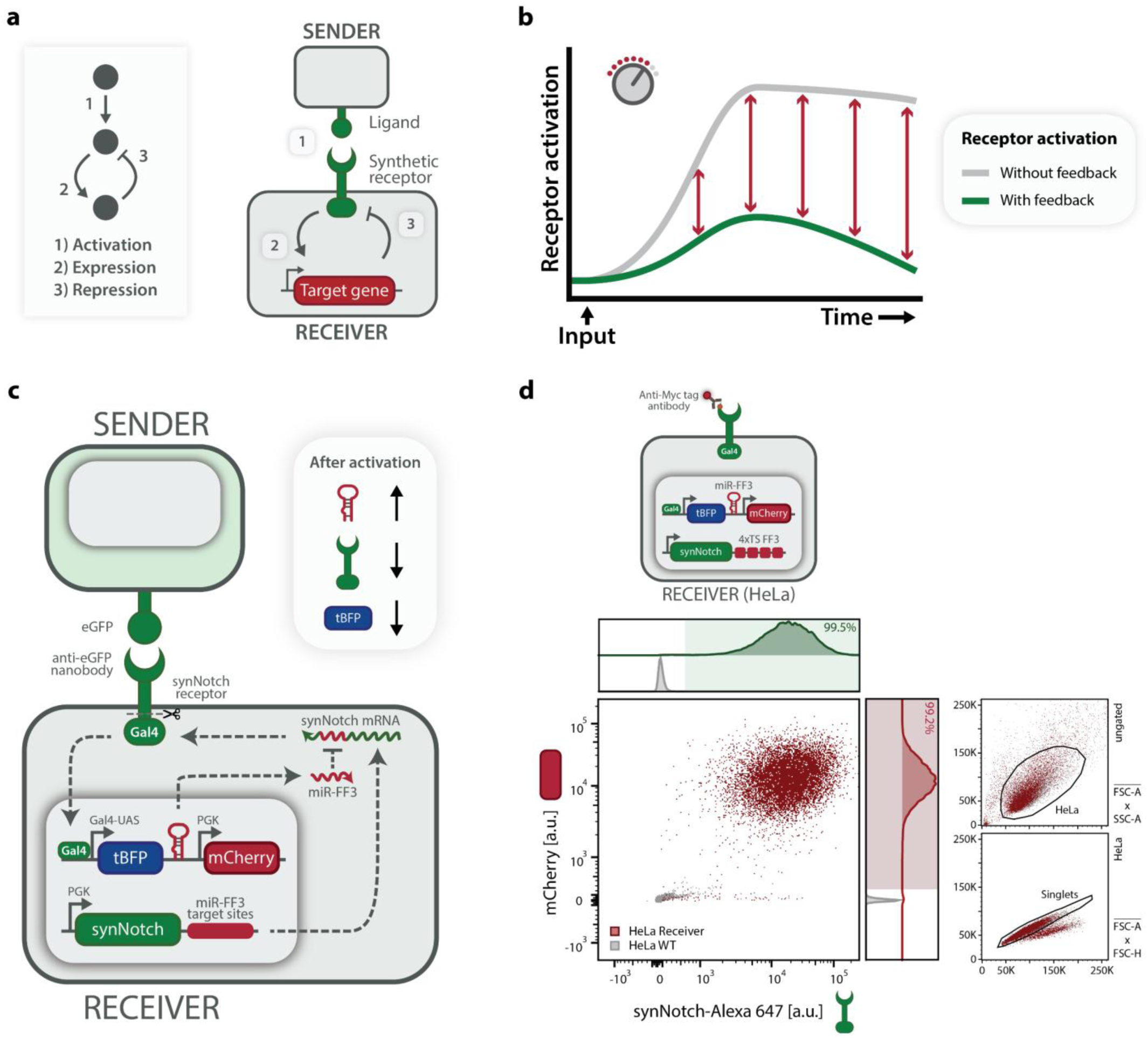
Design elements of a microRNA (miRNA)-based negative feedback circuit for autonomous control of synthetic receptor expression and activity. **a** Simple schematic of the proposed strategy to control synthetic receptor activation by implementing a negative feedback loop. Upon synthetic receptor activation by its cognate ligand (1), a target gene is expressed (2) that is designed to repress receptor expression and consequent activity (3). **b** Conceptual schematic of receptor activation dynamics with and without feedback. In the absence of a feedback mechanism (grey), receptors remain constitutively activated, whereas with negative feedback (green), receptor activity is attenuated over time. **c** Detailed schematic of the miRNA-based negative feedback circuit for autonomous control of synthetic receptor activation. HeLa cells are stably transduced (Methods) to constitutively express a synthetic Notch (synNotch) receptor, composed of an anti-eGFP nanobody, and a Gal4 transcriptional effector domain, under PGK promoter control. The synNotch receptor gene is engineered to contain complementary miR-FF3 target sites (Supplementary Fig. S3 and Table S4) in its transcript that are recognized by a synthetic miRNA, miR-FF3 (Supplementary Fig. S4 and Table S3). SynNotch receptor activation by a membrane-tethered eGFP ligand, expressed by engineered K562 *Sender* cells (Supplementary Fig. S1 and Methods), induces Gal4-UAS-driven expression of both tBFP reporter and miR-FF3, residing in a miR-30 backbone (Supplementary Fig. S2 and Table S3). Mature miR-FF3 can bind to its complementary miR-FF3 target sites (TS) in the synNotch mRNA, leading to inhibition of synNotch expression. Additionally, *Receiver* cells constitutively express the fluorescent protein mCherry under PGK promoter control. d Flow cytometry validation of engineered HeLa *Receiver* cells. Scatter plot and histograms of synNotch (stained with anti-Myc AF647 antibody) versus mCherry fluorescence confirm successful integration of both gene constructs in HeLa *Receiver* cells (red) compared to untransduced HeLa wild-type (WT) control (grey). Right panels show the gating strategy (Supplementary Fig. S6a) (HeLa: FSC-A vs. SSC-A; Singlets: FSC-A vs. FSC-H). PGK: Phosphoglycerate Kinase; UAS: Upstream Activating Sequence; eGFP: enhanced Green Fluorescent Protein; tBFP: tag Blue Fluorescent Protein; FSC: Forward Scatter; SSC: Side Scatter.

In this work, we establish a synthetic receptor platform equipped with ligand-triggered, genetically encoded miRNA-mediated feedback to autonomously regulate synthetic receptor expression and thereby modulate downstream activation. Specifically, we design a synthetic gene circuit in which synNotch activation induces expression of an orthogonal synthetic miRNA that targets synNotch mRNA, thereby creating a self-limiting circuit in which receptor activation directly drives its own repression. Conceptually, such a negative feedback loop is expected to attenuate receptor activation over time, in contrast to constitutive signalling observed in systems without a feedback mechanism (Fig. 1b). We first demonstrate that an orthogonal synthetic miRNA can robustly repress synNotch expression and subsequent activation, and that the degree of repression is tunable, scaling with the number of miRNA target sites incorporated in the receptor transcript. We then show that synNotch activation can induce expression of a functional synthetic miRNA, capable of downregulating the expression of an independent reporter construct. Having validated these components independently, we integrate them into a complete feedback loop architecture and analyse the system’s behaviour over time. We reveal that the circuit autonomously attenuates synNotch expression following activation, resulting in transient, self-limiting downstream reporter expression that peaks before gradually declining as feedback engages. In addition, we develop a simple ordinary differential equation (ODE) model of the circuit that reproduces the key dynamic features of our system, reinforcing our experimental observations with a quantitative framework. Altogether, these results demonstrate that miRNA-based negative feedback enables autonomous regulation of synthetic receptor activity and establishes a generalizable framework with tunable design features for receptor systems with built-in homeostatic control, guiding the development of robust, self-regulating synthetic circuits, and laying a foundation for future therapeutic applications.

## Results

### Design principles of a miRNA-mediated feedback mechanism for synNotch receptor control

To implement our proposed feedback strategy, we designed a synNotch-based gene circuit in which synNotch receptor activation triggers expression of a synthetic, orthogonal miRNA that specifically targets synNotch mRNA, enabling autonomous feedback regulation (Fig. 1c). Upon ligand engagement of the receptor, this architecture simultaneously initiates downstream reporter gene activation and expression of the regulatory miRNA, which is designed to ultimately limit receptor abundance.

In detail, the circuit was built around a synNotch receptor composed of an extracellular anti-eGFP nanobody (LaG17), a native Notch transmembrane domain, and an intracellular Gal4-VP64 transcriptional effector, constitutively expressed under PGK promoter control, as originally described by Morsut et al. (2016)^41^. Receptor activation by a membrane-tethered eGFP ligand expressed on the surface of engineered K562 *Sender* cells (Supplementary Fig. S1) leads to proteolytic cleavage of the synNotch receptor and release of the Gal4-VP64 effector domain. This transcription factor activates the Gal4-UAS promoter to drive transcription of a bicistronic cassette encoding both the blue fluorescent reporter tagBFP (tBFP) and a synthetic orthogonal miRNA, miR-FF3^33,47^, which is embedded in a miR-30 backbone^30,34^ and positioned in the 3’ UTR of the tBFP gene^30,48,49^ (Supplementary Fig. S2 and Table S3). Subsequently, mature miR-FF3 targets complementary miRNA target sites (TS) introduced in the 3’ UTR of the synNotch transcript^33,34,50^ (Supplementary Figs. S3-S4, and Table S4), linking receptor activation to its own repression through post-transcriptional feedback, effectively concluding the envisioned negative feedback loop.

To embed this circuit in mammalian cells, we engineered HeLa *Receiver* cells by stably integrating two lentiviral constructs encoding: (i) constitutive expression of the synNotch receptor with four miR-FF3 target site repeats (Supplementary Fig. S3 and Table S5), and (ii) synNotch-driven inducible expression of the tBFP reporter and miR-FF3, together with constitutive expression of mCherry under PGK promoter control (Supplementary Fig. S2, Table S3, and Methods). Following lentiviral transduction, *Receiver* cells were sorted by fluorescence-activated cell sorting (FACS) to enrich for cells co-expressing synNotch receptor and mCherry (Methods). Flow cytometry analysis post-sorting confirmed successful integration of both receptor and reporter constructs (Fig. 1d and Supplementary Fig. S5).

### Functional validation of miRNA-mediated repression of synNotch expression and signalling

Prior to implementing and testing the entire feedback loop architecture within a single cell line, we first sought to evaluate the functionality of its individual components. Specifically, we aimed to confirm whether the miRNA target sites introduced into the synNotch receptor transcript could mediate effective repression of receptor expression in the presence of a complementary miRNA. We designed a set of experiments to investigate whether synthetic precursor miRNA can downregulate synNotch expression and consequently attenuate downstream signalling activity (Fig. 2a). To this end, we transfected *Receiver* cells containing miR-FF3 target sites with synthetic precursor miRNAs and analysed synNotch receptor as well as tBFP reporter expression by flow cytometry and live-cell imaging (Methods).

**Figure 2:**
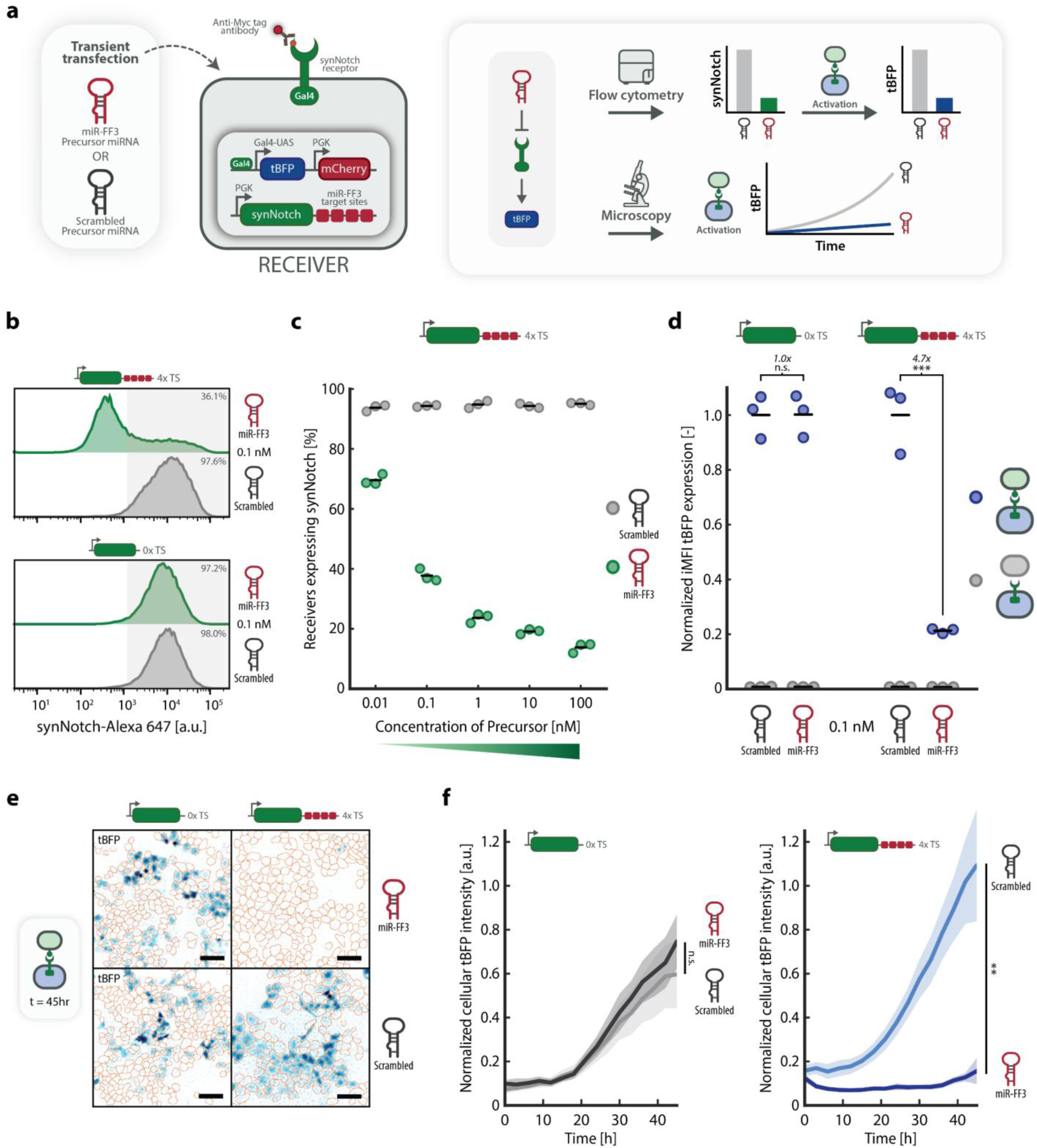
SynNotch expression and subsequent activation are repressed by miR-FF3 in a target site-dependent manner. **a** Precursor miRNA transfection setup. *Receiver* cells containing four miR-FF3 target sites (4xTS; Supplementary Fig. S7) are transiently transfected with complementary (miR-FF3) or scrambled (negative control) mirVana™ precursor miRNA. Following transfection, cells are either analysed for synNotch expression by flow cytometry, or activated and subsequently analysed for tBFP expression by flow cytometry and live-cell imaging (Methods). Schematics (right) illustrate the expected repression of synNotch and downstream signalling. **b** SynNotch expression 72 h post-transfection. Flow cytometry histograms show synNotch expression (anti-Myc AF647 staining) in cells hosting 4xTS or 0xTS, transiently transfected with 0.1 nM miR-FF3 (green) or scrambled (grey) precursor miRNA. Shaded area indicates synNotch-positive cells (threshold defined by HeLa WT control; Supplementary Fig. S9a). **c** Dose-response of synNotch repression. Percentage of synNotch-positive cells, quantified by flow cytometry analysis in *Receiver* cells with 4xTS, 72 h post-transfection with 0.01, 0.1, 1, 10 or 100 nM miR-FF3 (green) or scrambled (grey) precursor miRNA (Supplementary Fig. S12). Horizontal lines indicate mean expression; individual data points represent independent replicates seeded, transfected, and analysed in parallel (n=3 independent wells per condition). **d** Downstream effect of synNotch repression. *Receiver* cells with 0xTS (left) or 4xTS (right) were transiently transfected with 0.1 nM miR-FF3 or scrambled precursor miRNA. After 24 h incubation, *Receiver* cells were co-cultured for 48 h with K562 eGFP+ *Sender* cells (blue) or K562 WT cells (grey) (Methods). Plot shows integrated Median Fluorescence Intensity (iMFI) of tBFP expression (normalized to activated *Receiver*, with 0xTS or 4xTS, transfected with scrambled precursor), derived by flow cytometry analysis (Supplementary Fig. S6b and S14). Horizontal lines indicate mean expression; individual data points represent independent replicates seeded, transfected, and analysed in parallel (n=3 independent wells per condition). Fold change and significance (two-tailed, unpaired t-test) are noted above data points (Supplementary Table S8). **e** Representative epifluorescence images showing tBFP expression in *Receiver* cells with 0xTS (left) or 4xTS (right), transfected with 0.1 nM miR-FF3 (top) or scrambled (bottom) precursor miRNA and co-cultured with K562 *Senders* (not shown), at 45 h post-activation (Supplementary Fig. S15). tBFP fluorescence is displayed in blue; orange outlines indicate cellular boundaries derived from mCherry-guided image segmentation (Methods). Scale bar: 100 µm. **f** Time-lapse quantification of tBFP expression (measured every 3 hours) of *Receiver* cells with 0xTS (left) or 4xTS (right), transfected with 0.1 nM precursor miRNA and co-cultured with K562 *Senders* as in Fig. 2e (Supplementary Fig. S16a and Video S1). Lines and shaded areas indicate the mean and 95% CI of normalized cellular tBFP intensity, quantified using widefield microscopy, of n=12 replicates (fields of view; FOV) from 3 independently seeded and transfected wells per condition with 4 FOVs per well. Significance (two-tailed linear mixed effects model with ‘well’ treated as random intercept) is shown for endpoint measurements (45 h post-activation; Supplementary Fig. S16b and Table S10). n.s.: p > 0.05, *p ≤ 0.05, **p ≤ 0.01, ***p ≤ 0.001.

We initially assessed miRNA-mediated repression of synNotch expression by transiently transfecting *Receiver* cells containing four miR-FF3 target site repeats (4xTS; Supplementary Fig. S7) or no target sites (0xTS; Supplementary Fig. S8) with either a complementary miR-FF3 precursor or a non-complementary control (scrambled) precursor (Methods). At 72 hours post-transfection, the synNotch receptor was stained with an anti-Myc fluorescent antibody, and receptor levels were quantified by flow cytometry. Flow cytometry analysis revealed a pronounced reduction in synNotch expression in 4xTS *Receivers* transfected with miR-FF3 precursor compared to those transfected with scrambled precursor (Fig. 2b and Supplementary Fig. S9). In contrast, synNotch levels in *Receiver* cells lacking target sites (0xTS) remained unchanged regardless of the transfected precursor. To confirm sequence specificity of this repression, we similarly transfected *Receiver* cells harbouring alternative, non-complementary target sites – FF4, FF5^34^, or a heterologous sequence described by Endo et al. (2019)^36^ (Supplementary Table S4-S5, and Fig. S10-S11) – with the miR-FF3 precursor. In these cell lines, miR-FF3 transfection had no significant effect on synNotch expression (Extended Data Fig. 1), demonstrating that the miRNA-mediated repression is sequence-specific and that the miR-FF3 miRNA-target site pair functions as an orthogonal regulatory module for receptor control with respect to the tested non-cognate synthetic target site controls.

To determine whether miRNA-mediated repression of synNotch is dose-dependent, we next transfected 4xTS *Receivers* with increasing concentrations of miR-FF3 or scrambled precursor and quantified receptor expression after 72 hours (Fig. 2c and Supplementary Fig. S12). This titration experiment revealed a graded decrease in the percentage of synNotch-positive cells with increasing miR-FF3 concentrations, indicating that receptor repression scales with miRNA dose. In contrast, cells transfected with scrambled precursor maintained stable synNotch expression across all concentrations.

To examine whether miRNA-mediated repression of synNotch expression translates to functional attenuation of downstream signalling, we next evaluated tBFP reporter expression in *Receiver* cells following precursor transfection. First, we tested receptor activation and tBFP expression in our control 0xTS *Receiver* cells without precursor transfection. Upon co-culture with varying amounts of eGFP-expressing K562 *Sender* cells, these control *Receivers* showed robust reporter expression, confirming receptor activation and circuit function (Supplementary Fig. S13). We subsequently compared synNotch activation in the 0xTS *Receivers* to *Receivers* containing 4xTS, transfecting both cell lines with either 0.1 nM of miR-FF3 or scrambled precursor, before co-culturing with eGFP-expressing *Senders* or wild-type K562 cells for 48 hours. Flow cytometry analysis of tBFP reporter expression revealed that, upon activation with *Sender* cells, 4xTS *Receivers* transfected with miR-FF3 precursor exhibited a significant 4.7-fold reduction in tBFP levels, indicating that reduced receptor abundance leads to dampened downstream signalling (Fig. 2d and Supplementary Fig. S14). In contrast, 0xTS *Receivers* maintained comparable tBFP expression regardless of precursor type.

To further substantiate the flow cytometry results, we used live-cell imaging to visualize reporter expression dynamics (Fig. 2e). Fluorescence microscopy showed reduced tBFP fluorescence intensity in 4xTS *Receivers* transfected with miR-FF3 compared to scrambled controls, whereas 0xTS *Receivers* exhibited similar reporter expression across conditions (Fig. 2e and Supplementary Fig. S15). Quantification of tBFP dynamics over the activation period confirmed these trends, with 0xTS *Receivers* showing overlapping expression trajectories across conditions, and 4xTS *Receivers* transfected with miR-FF3 displaying attenuated reporter induction, culminating in significantly lower tBFP expression at the 45-hour post-activation endpoint (Fig. 2f, Supplementary Fig. S16, and Video S1). These results demonstrate that live-cell imaging effectively captures the temporal dynamics of synNotch activation and confirm that miRNA-mediated repression produces a sustained reduction in downstream reporter activity.

### Target site multiplicity enables tunable repression of synNotch and downstream signalling

To investigate whether the extent of miRNA-mediated repression can be systematically modulated, we next examined how the number of miRNA target sites within the synNotch receptor transcript influences repression strength. Previous studies have shown a correlation between miRNA target site multiplicity and repression strength, with transcripts containing more target sites exhibiting progressively stronger silencing^37,40,51,52^. We therefore reasoned that varying the number of miR-FF3 target sites within the synNotch 3’ UTR would allow us to tune the degree of miRNA-mediated repression and, consequently, modulate both receptor abundance and downstream signalling activity.

To systematically assess how the number of target sites affects miRNA-mediated repression, we constructed a library of *Receiver* cell lines in which the synNotch receptor transcript either contained 0, 1, 2, 3, or 4 tandem repeats (0-4xTS) of the miR-FF3 target site (Supplementary Fig. S7-S8 and S17-S18). Each *Receiver* cell line was then transiently transfected with miRNA precursor and analysed for synNotch receptor abundance and downstream tBFP reporter activation by flow cytometry (Fig. 3a). We hypothesised that target site multiplicity would proportionally scale miRNA-mediated repression of synNotch and thereby modulate downstream signalling output.

**Figure 3:**
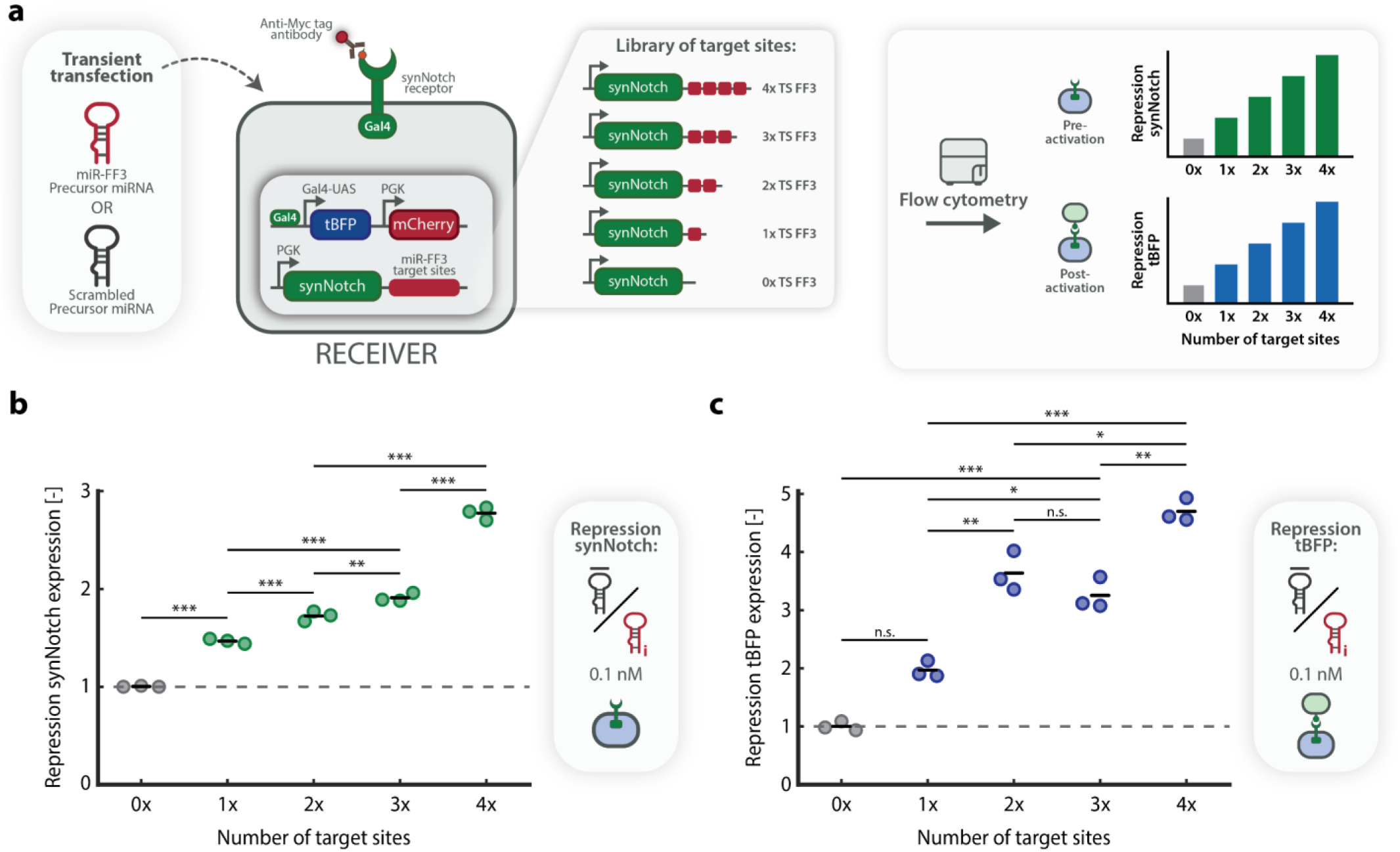
SynNotch repression and activity are tunable and correlate with the number of miRNA target sites. **a** Experimental scheme for testing target site-dependent repression. A library of engineered *Receiver* cell lines, containing 0, 1, 2, 3 or 4 miR-FF3 target sites (TS; Supplementary Fig. S7-S8 and S17-S18), is transiently transfected with complementary (miR-FF3) or scrambled (negative control) mirVana™ precursor miRNA (Methods). Following transfection, cells are either analysed for synNotch expression (anti-Myc AF647 staining) or activated and subsequently analysed for tBFP expression by flow cytometry. Schematics (right) illustrate the expected effect of the number of TS on the repression of synNotch and tBFP. **b** Target site-dependent repression of synNotch expression, quantified by flow cytometry analysis (Extended Data Fig. 2b-2c), in *Receiver* cells with 0, 1, 2, 3, or 4xTS, 72 h post-transfection with 0.1 nM precursor miRNA. Repression of synNotch is calculated as the mean percentage of synNotch-positive cells transfected with scrambled precursor, divided by the percentage of synNotch-positive cells transfected with miR-FF3 precursor. Horizontal lines indicate mean repression; individual data points represent independent replicates seeded, transfected, and analysed in parallel (n=3 independent wells per condition). **c** Target site-dependent repression of tBFP expression, quantified by flow cytometry analysis (Extended Data Fig. 2d-2e). *Receiver* cells with 0, 1, 2, 3, or 4xTS were transiently transfected with 0.1 nM precursor miRNA. Following 24 h incubation, *Receiver* cells were co-cultured for 48 h with K562 eGFP+ *Sender* cells (Methods). Plot shows repression of tBFP expression, calculated as the mean iMFI of tBFP expression of *Receiver* cells transfected with scrambled precursor, divided by the iMFI of tBFP expression of *Receiver* cells transfected with miR-FF3 precursor. Horizontal lines indicate mean repression; individual data points represent independent replicates seeded, transfected, and analysed in parallel (n=3 independent wells per condition). The standard deviation of each group of data points was corrected for error propagation (S.D. of individual data points not shown). Significance (one-way ANOVA with Tukey’s multiple comparisons test) of selected comparisons is noted above data points (full overview of statistics in Supplementary Table S9). n.s.: p > 0.05, *p ≤ 0.05, **p ≤ 0.01, ***p ≤ 0.001.

To determine how the number of miRNA target sites affects synNotch repression, we first quantified receptor abundance across the 0-4xTS *Receiver* panel following transfection with miR-FF3 or scrambled precursor. Flow cytometry analysis revealed a progressive and significant increase in synNotch repression for increasing numbers of miR-FF3 target sites, with 1xTS *Receivers* showing a modest 1.5-fold increase in repression and 4xTS *Receivers* exhibiting a more pronounced 2.8-fold increase in repression compared to 0xTS *Receivers* (Fig. 3b and Extended Data Fig. 2b-2c).

Next, we examined whether the graded repression of synNotch receptor levels translates into corresponding changes in downstream signalling output. To this end, each *Receiver* cell line was transfected with miRNA precursor, activated by co-culture with eGFP-expressing K562 *Sender* cells and then analysed for tBFP reporter expression by flow cytometry. Since the different *Receiver* cell lines exhibit modest differences in their baseline activation levels, signalling repression was calculated as the ratio of tBFP expression in *Receiver* cells transfected with scrambled precursor to that in cells transfected with miR-FF3 precursor. Using this metric, we observed minimal attenuation of tBFP expression in 1xTS *Receivers* with a 2.0-fold increase in tBFP repression compared to 0xTS *Receivers.* Moreover, 2xTS and 3xTS *Receivers* showed intermediate repression of tBFP (3.2- to 3.6-fold), and strongest repression was observed in the 4xTS *Receiver* cell line (4.7-fold; Fig. 3c and Extended Data Fig. 2d-2e). Although tBFP repression between 2xTS and 3xTS lines did not differ significantly, the overall increase in signalling repression with higher target site multiplicity indicates that miRNA-mediated tuning of synNotch abundance propagates to downstream signalling output. Together, these findings demonstrate that miRNA target site multiplicity provides a rational design parameter that enables graded tuning of receptor abundance and modulation of the resulting signalling response.

### SynNotch signalling drives inducible expression of functional miRNA

Having established that exogenous precursor miRNA transfection can repress synNotch expression in a target site-dependent and tunable manner, we next sought to evaluate whether synNotch activation can induce expression of functional miR-FF3, the second key component of our miRNA-based feedback architecture. To this end, we examined both the expression level of miR-FF3 upon synNotch activation and whether this miRNA could repress an independent cognate target transcript. To evaluate synNotch-driven miRNA induction, we first engineered a dedicated miR-FF3 *Receiver* cell line in which the miRNA is encoded within a miR-30 backbone, placed in the 3’ UTR of the GAL4-UAS-inducible tBFP reporter^30,48,49^ (Supplementary Fig. S19). We reasoned that high expression of the reporter cassette would be necessary to achieve robust induction of both tBFP and miR-FF3. Since the constitutive mCherry marker is encoded on the same construct, we enriched both the original (0xTS) and miR-FF3 *Receiver* populations for high mCherry expression, yielding cells with higher transgene abundance. Although this enrichment increased basal tBFP levels, it also conferred stronger activation capacity, with substantially higher activation-induced tBFP expression than in the non-enriched populations (Supplementary Fig. S19-S21a). Moreover, we observed that activating *Receiver* cells using plate-bound eGFP ligand^41^ (Methods) resulted in higher tBFP expression than co-culture with eGFP-expressing *Sender* cells, prompting us to use plate-bound ligand for all subsequent miRNA-induction experiments (Supplementary Fig. S21b).

To assess whether synNotch activation induces expression of the synthetic miRNA, we activated the enriched miR-FF3 *Receiver* cells on plates coated with eGFP ligand, and quantified miRNA levels 72 hours later using quantitative real-time PCR (qPCR; Fig. 4a). Flow cytometry of tBFP fluorescence confirmed synNotch activation, with *Receiver* cells cultured on eGFP-coated plates showing a clear increase in reporter expression relative to non-activated cells cultured on uncoated plates (Fig. 4b, left). We subsequently harvested these cells for quantification of miR-FF3 expression by qPCR, using miR-103a-3p as the internal reference^53,54^ (Supplementary Fig. S22 and Methods). Activated miR-FF3 *Receivers* exhibited a pronounced 68.9-fold increase in mature miR-FF3 abundance compared to non-activated control miR-FF3 *Receivers* (Fig. 4b, right), demonstrating that synNotch activation can robustly induce expression of mature synthetic miRNA.

**Figure 4:**
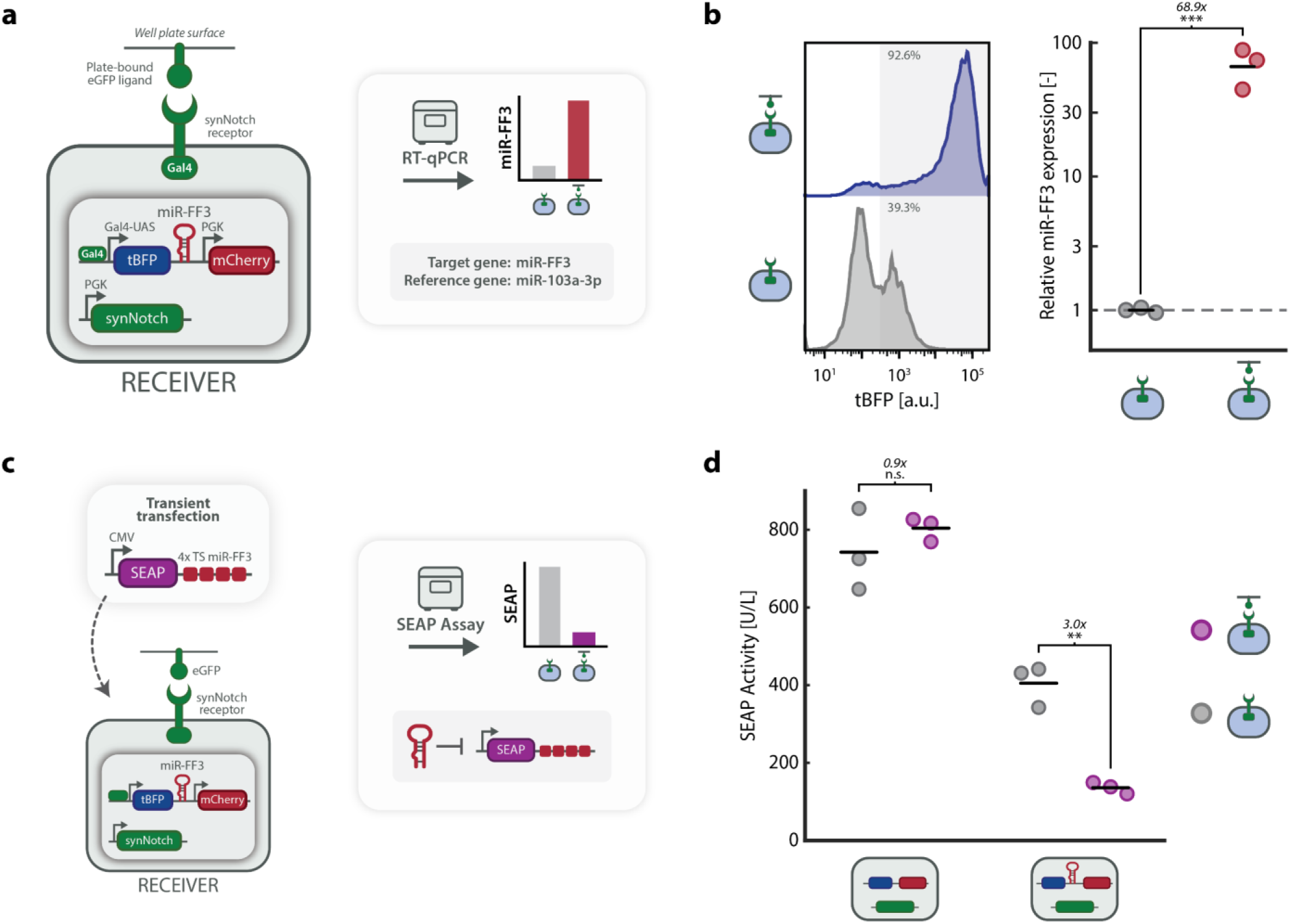
SynNotch activation induces expression of functional miR-FF3. **a** Experimental setup for miR-FF3 induction and quantification. *Receiver* cells engineered to have a reporter gene construct containing both tBFP and miR-FF3 genes (miR-FF3 *Receiver*; Supplementary Fig. S19) are activated using plate-bound eGFP ligand (Methods). Following a 72 h incubation period, cells are harvested and lysed for miRNA isolation. Isolated miRNA is reverse transcribed (RT) and miR-FF3 expression is quantified using quantitative real-time PCR (qPCR; Methods). Target gene: miR-FF3; Reference gene: miR-103a-3p (Supplementary Fig. S22 and Table S6) **b** Flow cytometry histogram (left) shows tBFP expression levels of activated and non-activated miR-FF3 *Receiver* cells. Relative miR-FF3 expression, normalized to non-activated miR-FF3 *Receiver* cells, was quantified using qPCR (right). Horizontal lines indicate mean expression; individual data points represent independent replicates seeded, lysed, and analysed in parallel (n=3 independent wells per condition). qPCR was performed in technical triplicates, which were averaged prior to analysis. Fold change and significance (two-tailed, unpaired lognormal t-test) are noted above data points (Supplementary Table S8). **c** Experimental design for functional validation of induced miR-FF3. MiR-FF3 *Receiver* cells are seeded on eGFP-coated plates and subsequently transfected with a plasmid containing a secreted embryonic alkaline phosphatase (SEAP) reporter gene, under the control of a CMV promoter, harbouring four miR-FF3 target sites (SEAP-4xTS; Supplementary Fig. S23). MiR-FF3 expression is expected to repress SEAP expression, quantified by a SEAP assay (Supplementary Fig. S24 and Methods). **d** SEAP activity [U/L], measured 48 h post-transfection, in *Receiver* cells lacking (left) or expressing (right) the miR-FF3 gene, seeded on uncoated (grey) or eGFP-coated plates (purple), and transiently transfected with the SEAP-4xTS reporter construct. Horizontal lines indicate mean activity; individual data points represent independent replicates seeded, transfected, and analysed in parallel (n=3 independent wells per condition). Fold change and significance (one-way ANOVA with Tukey’s multiple comparisons test) are noted above data points (Supplementary Table S9). n.s.: p > 0.05, *p ≤ 0.05, **p ≤ 0.01, ***p ≤ 0.001.

To evaluate whether the miR-FF3 produced upon synNotch activation is functionally active, we next tested its ability to repress an independent target transcript containing complementary target sites. For this purpose, we designed a reporter plasmid containing a constitutively expressed secreted embryonic alkaline phosphatase (SEAP) gene and four FF3 target site repeats, mirroring the target site architecture used in earlier assays (Supplementary Fig. S23). We then transiently transfected activated and non-activated miR-FF3 *Receiver* cells with this reporter plasmid and quantified SEAP expression using a colorimetric SEAP assay (Fig. 4c, Supplementary Fig. S24, and Methods). Activated miR-FF3 *Receivers* showed a significant 3.0-fold reduction in SEAP activity compared to non-activated miR-FF3 *Receivers*, demonstrating that synNotch activation induces sufficient levels of functional miR-FF3 to repress a target transcript containing complementary target sites (Fig. 4d). In contrast, SEAP activity remained unchanged between activated and non-activated control *Receiver* cells lacking the miR-FF3 gene, and an untargeted SEAP reporter lacking FF3 target sites was not repressed in activated miR-FF3 *Receivers* (Supplementary Fig. S25). Furthermore, we observed a modest reduction in SEAP levels in non-activated miR-FF3 *Receivers* compared to control *Receivers*, suggesting low-level background expression of miR-FF3 in the absence of ligand stimulation, consistent with previously observed tBFP background expression in these cells (Fig. 4b). Together, these results establish that synNotch activation produces functional miR-FF3 capable of repressing transcripts bearing cognate miR-FF3 target sites, providing the essential effector module required to implement the full miRNA-based feedback loop.

### Implementation of a miRNA-mediated feedback loop enables autonomous attenuation of synNotch signalling

Having established that miRNA-mediated repression of synNotch can durably attenuate downstream reporter signalling and that synNotch activation can drive inducible expression of functional miR-FF3, we next integrated these two components into a single *Receiver* cell line (enriched for high mCherry expression; Supplementary Fig. S5) to implement autonomous feedback regulation of synNotch abundance and downstream signalling. To assess the behaviour of the complete feedback circuit, we compared *Receiver* cells containing the full synNotch-miRNA feedback architecture to control *Receiver* cell lines lacking the miRNA gene (Supplementary Fig. S26-S27). Following activation with plate-bound eGFP ligand, synNotch receptor repression and downstream tBFP activation were quantified over time by flow cytometry, with live-cell imaging used to resolve higher-resolution dynamics of tBFP expression (Fig. 5a).

**Figure 5:**
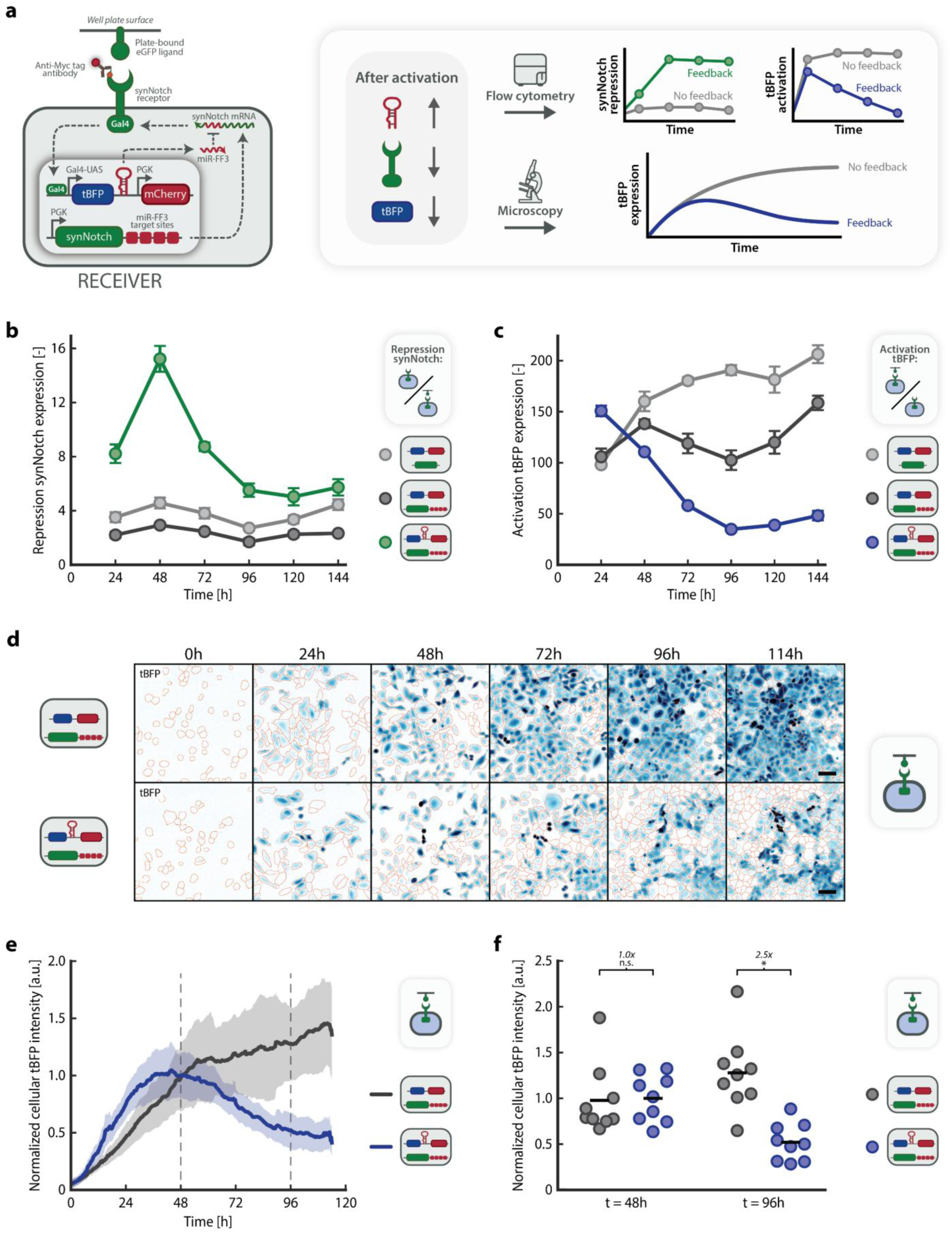
The synNotch-miRNA feedback loop autonomously represses synNotch receptor abundance and downstream reporter expression. **a** Experimental design and expected feedback dynamics. *Receiver* cells engineered to contain both the miR-FF3 gene and complementary miR-FF3 target sites are activated using plate-bound eGFP ligand and subsequently stained with anti-Myc AF647 antibody. Repression of synNotch and activation of tBFP in *Receiver* cells with and without the complete synNotch-miRNA feedback loop are tracked over time by flow cytometry, and live-cell imaging (Methods). Schematics (right) illustrate the expected effect of the synNotch-miRNA feedback loop on the repression of synNotch and activation of downstream reporter expression. **b-c** Time-course analysis of feedback circuit performance quantified by flow cytometry. Three types of *Receiver* cell lines (light grey: without feedback loop components, dark grey: without miR-FF3 gene and with miR-FF3 target sites, green/blue: with miR-FF3 gene and target sites; Supplementary Fig. S27) were seeded for activation at t = 0 h on uncoated or eGFP-coated plates (Methods). **b** Repression of synNotch expression over time was quantified by flow cytometry analysis, with measurements every 24 hours following activation. Repression of synNotch expression was calculated as the ratio of synNotch Median Fluorescence Intensity (MFI) in non-activated versus activated *Receiver* cells (Extended Data Fig. 3a-3b). Mean synNotch repression and standard deviation are shown for n = 3 independent replicates per timepoint (independent wells seeded and analysed in parallel). The standard deviation of each group of data points was corrected for error propagation (individual data points and their S.D. not shown). **c** Activation of tBFP expression over time was quantified by flow cytometry analysis, with measurements every 24 hours following activation. Activation of tBFP expression was calculated as the ratio of tBFP iMFI in activated versus tBFP MFI in non-activated *Receiver* cells (Extended Data Fig. 3c-3d). Mean tBFP activation and standard deviation are shown for n = 3 independent replicates per timepoint (independent wells seeded and analysed in parallel). The standard deviation of each group of data points was corrected for error propagation (individual data points and their S.D. not shown). **d-f** Time-course analysis of feedback circuit performance quantified using widefield microscopy. Two types of *Receiver* cell lines (dark grey: without miR-FF3 gene and with miR-FF3 target sites, blue: with miR-FF3 gene and target sites; Supplementary Fig. S27) were seeded for activation at t = 0 h on eGFP-coated plates (Methods). **d** Representative epifluorescence microscopy images showing tBFP expression of *Receiver* cells every 24 hours post-activation. tBFP fluorescence is displayed in blue; orange outlines indicate cellular boundaries derived from mCherry-guided image segmentation (Methods). Scale bar: 100 µm. **e** Time-lapse quantification of tBFP expression in activated *Receiver* cells. Lines and shaded areas indicate the mean and 95% CI of normalized cellular tBFP intensity, quantified every 15 minutes using widefield microscopy, of n=9 replicates from 3 biologically independent experiments (runs) performed on separate days with 3 wells per condition (Video S2). **f** Scatter plot showing normalized cellular tBFP intensity at 48 hours and 96 hours post-activation. Horizontal lines indicate the mean normalized cellular tBFP intensity; individual data points represent n=9 replicates from 3 biologically independent experiments (runs) performed on separate days with 3 wells per condition. Fold changes and significance (two-tailed linear mixed effects model with ‘run’ treated as random intercept) are noted above data points (Supplementary Table S11). n.s.: p > 0.05, *p ≤ 0.05, **p ≤ 0.01, ***p ≤ 0.001.

We first examined how implementation of the complete feedback circuit impacted synNotch receptor abundance by flow cytometry following ligand activation. In *Receiver* cells containing the full synNotch-miRNA feedback loop, synNotch expression was strongly repressed after activation, with repression emerging between 24 and 72 hours and peaking at approximately 48 hours post-activation (Fig. 5b). *Receiver* cells lacking the miRNA gene maintained high synNotch expression over the same time course, regardless of the presence of miRNA target sites in the synNotch transcript (Extended Data Fig. 3b and Supplementary Fig. S28). As synNotch activation is expected to reduce surface receptor levels through ligand-induced cleavage, activated cells show decreased synNotch abundance independent of miRNA expression (Extended Data Fig. 3a-3b). The additional reduction observed specifically in *Receiver* cells containing the miRNA gene compared to *Receiver* cells lacking the miRNA gene therefore isolates the contribution of miRNA-mediated repression, demonstrating that synNotch repression is driven by synNotch-inducible miRNA expression and unfolds in a time-dependent manner following receptor activation.

We next assessed how feedback-mediated repression of synNotch receptor abundance impacts downstream signalling output by quantifying tBFP activation over time. In *Receiver* cells containing the complete synNotch-miRNA feedback loop, tBFP activation markedly decreased following repression of synNotch receptor expression (Fig. 5c). *Receiver* cells lacking the miRNA gene maintained sustained tBFP expression over the same time course (Extended Data Fig. 3c-3d and Supplementary Fig. S29). These results indicate that repression of synNotch receptor expression by the feedback loop propagates to downstream transcriptional output, attenuating tBFP reporter expression over time.

To further resolve the temporal dynamics of miRNA-mediated feedback of synNotch expression and signalling, we next monitored tBFP expression of *Receiver* cell lines by live-cell imaging following ligand activation (Methods). Representative fluorescence microscopy images illustrate a pronounced reduction in tBFP fluorescence intensity in *Receiver* cells with the complete feedback loop compared to *Receivers* lacking the miRNA gene across the imaging time course (Fig. 5d and Supplementary Fig. S30). Quantification of cellular tBFP intensity in feedback loop *Receiver* cells revealed a transient response characterised by an initial increase in tBFP expression followed by pronounced attenuation over time. Cellular tBFP intensity reached its maximum around 48 hours post-activation and subsequently declined, consistent with autonomous attenuation of signalling output following activation (Fig. 5e, Supplementary Fig. S31, and Video S2). In contrast, *Receiver* cells lacking the miRNA gene maintained elevated tBFP expression over the same time window. Notably, *Receiver* cells containing the feedback loop exhibited a slightly steeper initial increase in tBFP expression compared to cells lacking the miRNA gene, likely reflecting inherent differences in gene expression between independently generated cell lines (Supplementary Fig. S27 and Extended Data Fig. 3c-3d). Despite this faster initial rise, feedback-containing *Receiver* cells ultimately displayed markedly lower tBFP levels at later time points once miRNA-mediated repression of synNotch took effect. In line with these observations, direct comparison of tBFP expression between *Receivers* with and without the complete feedback loop revealed no significant difference at 48 hours, whereas tBFP expression was significantly reduced in feedback loop *Receivers* at 96 hours post-activation (Fig. 5f). Together, these results demonstrate that integration of miRNA-mediated feedback into the synNotch receptor expression pathway ties ligand-induced activation to autonomous attenuation of receptor abundance, thereby controlling downstream signalling.

### Mathematical modelling qualitatively recapitulates robust transient reporter dynamics from miRNA-mediated regulation of synNotch signalling

To complement our experimental results and evaluate the robustness of miRNA-mediated regulation of synNotch signalling, we developed a mathematical ordinary differential equation (ODE) model that captures the key molecular processes of the feedback circuit. The model describes the dynamics of synNotch receptor expression, activation, and downstream transcriptional output, as well as miRNA-mediated repression of synNotch mRNA (Supplementary Fig. S32 and Supplementary Notes). Using this qualitative ODE model, we simulated the temporal dynamics of tBFP reporter expression following ligand-induced synNotch activation, comparing systems with and without miRNA-mediated regulation (Fig. 6a and Methods). In the absence of feedback, the model predicts a sustained increase in tBFP expression to a high steady state, reflecting continuous receptor activation. In contrast, inclusion of miRNA-mediated repression of synNotch produces a transient response, with tBFP expression peaking before declining to a lower steady state, qualitatively recapitulating the transient dynamics observed experimentally (Fig. 5c and 5e). Although absolute timescales differ, as model parameters were derived from literature values rather than fitted to experimental data, the qualitative behaviour is preserved between simulation and experiment. Incorporating ± 5% parameter variation into the model yields closely overlapping tBFP trajectories for repeated simulations of each condition, indicating limited sensitivity of the simulated behaviour to parameter uncertainty (Fig. 6a).

**Figure 6:**
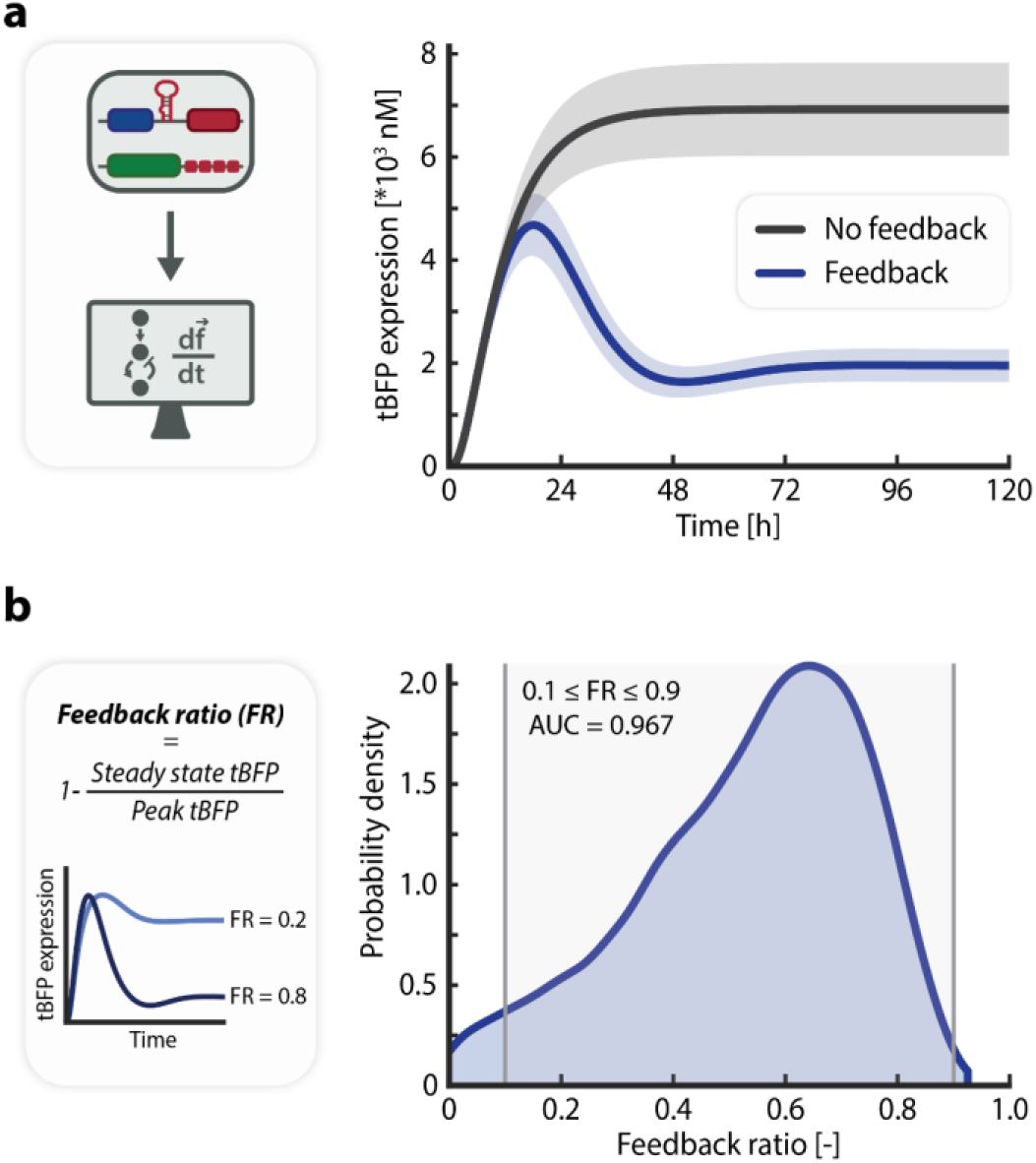
Mathematical modelling qualitatively recapitulates transient reporter dynamics. **a** A mathematical model capturing the key biological processes of the synNotch-miRNA feedback loop (Supplementary Fig. S32 and Supplementary Notes). The graph shows the simulated tBFP reporter expression over 120 hours in *Receiver* cells without (grey) or with (blue) miRNA-mediated regulation of synNotch following eGFP ligand exposure. Solid lines represent mean trajectories; shaded regions represent ± 2x S.D. derived from 10,000 Monte Carlo simulations with ± 5% parameter variation (Supplementary Notes and Methods). **b** Probability density distribution of the feedback ratio (FR) from 10,000 Monte Carlo simulations with parameter values uniformly sampled between 0.5x and 2x nominal values (Supplementary Notes and Methods). The FR quantifies regulatory strength of the feedback circuit for a set of parameter values and is defined as 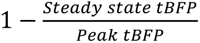. Low FR values indicate weak repression with a steady state tBFP expression close to peak tBFP expression, whereas high FR values indicate strong repression of tBFP expression. The graph (right) shows the probability density for each FR; grey shaded area indicates the probability of a parameter set yielding a feedback ratio within the range 0.1-0.9.

To further examine feedback behaviour across parameter space, we quantified system performance using a feedback ratio (FR) metric, defined as the fractional reduction in tBFP expression from peak to steady state resulting from miRNA-mediated repression (Fig. 6b). Model parameters were uniformly sampled over a twofold range around their nominal values, and FR distributions were obtained from 10,000 Monte Carlo simulations (Methods). We considered FR values between 0.1 and 0.9 to represent effective feedback, excluding regimes with minimal repression (FR < 0.1) or near-complete suppression of reporter expression (FR > 0.9). Across the twofold parameter variation explored, 96.7% of sampled parameter sets yielded FR values within the defined effective feedback range, indicating that effective miRNA-mediated feedback is preserved across substantial parameter variation. Together, these modelling results show that the ODE model recapitulates the emergence of the desired transient reporter dynamics and suggest that the regulatory properties designed into the synNotch-miRNA feedback loop are qualitatively robust in simulation across a range of plausible parameter values.

## Discussion

Precise control over synthetic receptor activity remains a central challenge in mammalian cell engineering, particularly in applications where prolonged or excessive signalling can compromise functional performance^11,12^. A key limitation of synthetic receptor systems is that they often lack intrinsic mechanisms to dynamically modulate receptor abundance or signalling strength following activation, resulting in excessive or sustained signalling outputs. In this study, we established a miRNA-based negative feedback strategy in which synNotch activation induces expression of an orthogonal synthetic miRNA that post-transcriptionally represses receptor expression, thereby coupling ligand recognition to autonomous attenuation of receptor abundance and downstream signalling output. We first demonstrated that synthetic miRNAs can mediate sequence-specific repression of synNotch expression and that this repression propagates to downstream signalling output. We further showed that repression strength can be tuned by varying the number of miRNA target sites, and that synNotch activation can robustly induce expression of a functional synthetic miRNA. By combining inducible miRNA expression with miRNA-mediated receptor repression in a single genetic circuit, we established an autonomous feedback loop in which ligand activation triggered delayed repression of synNotch receptor abundance and progressive attenuation of downstream signalling output over time. Together, these findings identify miRNA-mediated feedback as a compact, genetically encoded strategy for coupling ligand-triggered receptor activation to self-limiting attenuation of synthetic receptor activity, with target-site design providing a tunable parameter for controlling receptor repression strength.

A key feature of this approach is that feedback is encoded directly within the genetic circuit, enabling activation-dependent, self-limiting regulation without the need for external inputs. Because miRNA-mediated repression operates through sequence-specific interactions, regulatory control can be specified through target-site design, allowing precise and programmable modulation of gene expression. In addition, the modular nature of miRNA target-site incorporation and miRNA expression should enable straightforward integration of this strategy into diverse synthetic receptor systems without requiring substantial redesign of the underlying circuit architecture. Unlike alternative feedback implementations based on transcriptional repressors^55,56^ or protein-level degradation^57^, miRNA-mediated feedback acts directly at the post-transcriptional level to reduce receptor transcript abundance. In addition, sequence-specific target recognition makes miRNA-mediated feedback orthogonal, modular, and tunable through target-site design, without requiring re-engineering of protein domains or dependence on exogenous small-molecule inputs^19–22^. More broadly, our design is conceptually consistent with natural miRNA-based feedback regulation of endogenous receptor signalling^23–29^.

The feedback architecture described here also offers multiple points of tunability that can be leveraged to shape circuit behaviour. In the current work, we modulated receptor repression strength by varying the number of miRNA target sites present within the transcript of the synNotch receptor, providing a direct means to adjust the extent of receptor attenuation and downstream signalling. More broadly, future implementations could tune feedback dynamics across multiple layers within the circuit, including miRNA target site number and sequence complementarity, expression levels of the miRNA and the target transcript, and receptor-level properties such as ligand-binding affinity. This design space is consistent with recent work showing that compact miRNA modules can support precise, tunable, and dosage-invariant control of transgene expression across broad gene dosages and diverse cellular contexts, suggesting that the design principles underlying our feedback circuit are compatible with emerging miRNA engineering frameworks^38–40^. While the mathematical model presented here was intended to capture the qualitative behaviour of the feedback circuit rather than provide a fitted quantitative description of the experimental system, computational modelling could, in principle, be used to guide the selection and combination of these parameters to achieve desired signalling profiles. Moreover, the demonstrated orthogonality between miR-FF3 and alternative target-site sequences suggests that this framework could support multiplexed regulation, in which distinct receptor variants are independently controlled by different synthetic miRNA-target site pairs within the same cellular context^33,41^. Such modularity and tunability provide a rational design basis for fine-tuning feedback dynamics in future implementations of a miRNA-mediated feedback circuit.

While the present study establishes this feedback strategy in a cellular engineering context, it provides a proof-of-principle demonstration of an approach that could be adapted in future work for therapeutically relevant synthetic receptor systems. In previously described synNotch-CAR circuits, where synNotch recognition of a priming antigen is used to induce expression of a CAR directed against a secondary antigen^58^, our feedback design offers a route to tune inducible CAR expression and limit excessive or prolonged CAR activity following target antigen recognition. By coupling CAR expression to an activation-dependent negative feedback module, such systems have the potential to achieve more controlled and self-limiting responses. More broadly, analogous ligand-dependent feedback strategies can be implemented directly at the level of CAR expression and signalling, incorporated into combinatorial receptor designs such as AND-gate logic^46,59,60^, or extended to other classes of synthetic receptors. Although these possibilities remain to be explored experimentally, they highlight the potential of miRNA-mediated feedback to enhance the safety, controllability, and functional precision of engineered cell therapies.

The present work provides a foundation for extending miRNA-mediated feedback control towards more therapeutically relevant and translationally oriented synthetic receptor applications. A key next step will be to determine how well the synNotch-miRNA feedback architecture generalizes to primary and other therapeutically relevant cell types, and ultimately *in vivo*, particularly in settings requiring sustained or dynamic receptor control. Because synthetic gene circuit performance can depend strongly on cellular context and shared gene-expression resources^61,62^, circuit behaviour may vary across different implementation settings. Encouragingly, recent work has shown that model-guided miRNA circuit design can support precise transgene dosage across diverse primary cell types, providing a useful framework for adapting related feedback architectures to such cellular contexts^39^. Taken together, the findings presented in this study position autonomous miRNA-mediated feedback control of synthetic receptors as a general strategy for making synthetic receptor systems in mammalian cells more controllable, tunable, and adaptive.

## Methods

### Chemicals and reagents

All reagents and solvents used in this study were obtained from commercial sources and were used without further purification.

### Gene construct design and preparation

Plasmid backbones, engineered constructs, and insert sequences used in this study are listed in Supplementary Tables S1-S5, and plasmids generated in this work have been deposited to Addgene (Supplementary Table S2). The majority of engineered constructs were designed for this study and synthesized and cloned by GenScript Biotech Corporation into the indicated vectors, with sequence verification performed by GenScript Biotech Corporation prior to use. To generate the SEAP reporter construct containing miR-FF3 target sites, a 4xTS-FF3 cassette was cloned into the 3’ UTR of a CMV-driven SEAP plasmid (pBluescript_CMV_SEAP; Addgene #24595) by Gibson assembly following PCR amplification of insert and backbone using Q5 High-Fidelity DNA Polymerase (NEB, M0491), gel purification, and transformation into TOP10 competent *E. coli* cells (Invitrogen, C4040). Correct clones were identified by colony PCR, confirmed by Sanger sequencing (GENEWIZ), and propagated in TOP10 competent *E. coli* cells (Invitrogen, C4040). All other engineered plasmids were propagated in NEB Stable competent *E. coli* cells (NEB, C3040) and purified using the QIAprep Spin Miniprep Kit (QIAGEN, 27104) according to the manufacturer’s instructions prior to downstream applications.

### Cell culture

HeLa cells (ATCC, CCL-2), Lenti-X 293T cells (Takara Bio, 632180), and K562 cells (ATCC, CCL-243) were cultured in DMEM (Gibco, 41966) supplemented with 10% (v/v) FBS (Greiner Bio-One, 758093) and 1% (v/v) penicillin-streptomycin (Gibco, 15140), hereafter referred to as growth medium. Cells were maintained at 37 °C in a humidified incubator with 5% CO_2_ and passaged every 2-3 days to maintain exponential growth. Cell line authentication and mycoplasma testing were performed by the suppliers before distribution. Fresh cell cultures were regularly established from frozen aliquots. Adherent cells were detached using trypsin-EDTA (Gibco, 25200) for routine passaging, or enzyme-free Cell Dissociation Buffer (Gibco, 13150) two passages prior to the start of an experiment to prevent cleavage of synNotch receptor.

### Lentivirus production

Lentivirus was produced by co-transfecting Lenti-X 293T cells (Takara Bio, 632180), at ∼80% confluency in a T150 flask, with the second generation pHR plasmid carrying the desired transgene (Supplementary Table S2), the packaging plasmid (pCMVR8.74; Supplementary Table S1), and the VSV-G envelope plasmid (pMD2.G; Supplementary Table S1) at a mass ratio of 1:2:1.2 (50.4 µg of plasmid DNA in total) using FuGENE HD Transfection Reagent (Promega, E2311) in Opti-MEM I Reduced Serum Medium (Gibco, 31985062). The following day, the medium was refreshed to DMEM (Gibco, 41966) supplemented with 2% (v/v) heat-inactivated FBS (Greiner Bio-One, 758093), and cells were incubated under standard culture conditions (37 °C, 5% CO_2_). After 48 h, the medium was harvested and filtered with a 0.45 µm syringe filter (Sartorius, 16533). The produced lentivirus was then pelleted by centrifugation at 50,000x *g*, at 4 °C for 2 h, resuspended in 150 µL DMEM (Gibco, 41966) supplemented with 2% (v/v) heat-inactivated FBS (Greiner Bio-One, 758093), and either used directly for cell line transduction or snap-frozen and stored at -80 °C.

### Generation of stable K562 *Sender* and HeLa *Receiver* cell lines

Stable K562 *Sender* and HeLa *Receiver* cell lines were generated by lentiviral (co-)transduction using the lentiviral vectors as described in Supplementary Table S7. For transduction, WT cells were seeded in 24-well plates at ∼4x10^4^ cells per well. The following day, cells were incubated with lentiviral particles diluted 1:300 in 500 µL transduction medium consisting of DMEM (Gibco, 41966) supplemented with 10% (v/v) heat-inactivated FBS (Greiner Bio-One, 758093) and polybrene (10 µg/mL; Merck, TR-1003) for 48-72 h under standard culture conditions (37 °C, 5% CO_2_). Cells were subsequently expanded in growth medium prior to flow cytometric characterisation and fluorescence-activated cell sorting as described below. Viral titer of lentivirus was not formally determined; dilution ratio for transduction was empirically optimised and downstream fluorescence-activated cell sorting was used to select for successful transduction.

### Flow cytometry analysis

Adherent cells were harvested using trypsin-EDTA (Gibco, 25200) and resuspended in PBS supplemented with 0.5% (w/v) BSA (Bovine Serum Albumin; Merck, A7906). For detection of synNotch receptor expression, cells were harvested using enzyme-free Cell Dissociation Buffer (Gibco, 13150) and incubated with anti-Myc-tag antibody (9B11) conjugated to Alexa Fluor 488 (Cell Signaling, 2279S) or Alexa Fluor 647 (Cell Signaling, 2233S), diluted 1:50 in PBS containing 0.5% (w/v) BSA for 60 min at room temperature. Cells were subsequently washed with PBS containing 0.5% (w/v) BSA prior to flow cytometry analysis.

Flow cytometry data were acquired on a BD FACSymphony A3 Cell Analyzer (BD Biosciences) using the following channels: BB515 for eGFP/Alexa Fluor 488, APC for Alexa Fluor 647, PE-CF594 for mCherry, and BV421 for tBFP measurements. Events were first gated on FSC-A vs. SSC-A to identify the main cell population, followed by gating on FSC-A vs. FSC-H to retain single cells. In case of *Sender-Receiver* co-cultures, singlets were subsequently gated on positive mCherry expression to discriminate *Receiver* cells from *Sender* cells. A minimum of 10,000 events within the singlet gate (or mCherry+ gate for co-culture experiments) were recorded on the machine using the BD FACSDiva Software (BD Biosciences) and downstream analysis was performed in FlowJo (v10.8; BD Life Sciences; Supplementary Fig. S6). Positivity gates for each fluorescence channel were defined based on the 99.9^th^ percentile of the corresponding negative control populations and were applied consistently across samples within each experiment. SynNotch receptor expression was quantified either as the percentage of synNotch-positive cells or as the median fluorescence intensity (MFI) of the synNotch staining channel, depending on the experiment. Spillover of mCherry fluorescence into the APC/Alexa Fluor 647 channel was compensated using unstained mCherry-expressing *Receiver* cells as a reference control prior to quantification of synNotch staining. For activated samples, the integrated median fluorescence intensity (iMFI), calculated as the product of the percentage of tBFP-positive cells (defined relative to negative control populations) and the MFI of the tBFP-positive population, was used to capture the integrated reporter signal arising from the responding cell population. For non-activated samples, baseline tBFP expression was uniformly low and therefore quantified using the MFI. Using MFI for non-activated samples avoids instability caused by near-zero fractions of tBFP-positive cells, for which iMFI values approach zero and normalization becomes unreliable.

### Fluorescence-activated cell sorting

To enrich stable *Sender* or *Receiver* cell lines for positive expression of eGFP (*Senders)* or synNotch and mCherry expression (*Receivers)*, transduced cells were sorted by fluorescence-activated cell sorting on a BD FACSAria III Cell Sorter (BD Biosciences) using a 70 µm nozzle at low-medium pressure. Fluorescence intensity of fluorescent reporters was measured using the FITC channel for eGFP/Alexa 488, the mCherry channel for mCherry, and the Pacific Blue channel for tBFP. Cell samples were prepared for sorting as described for flow cytometry analysis. Cells expressing the synNotch receptor and/or fluorescent reporter markers (mCherry or eGFP) were isolated using gating strategies analogous to those used for analytical flow cytometry and collected as polyclonal populations. To enrich a population of *Receiver* cells for high expression of mCherry reporter, the top ∼20% highest mCherry-expressing cells were sorted. Sorted cells were collected in growth medium and subsequently expanded under standard culture conditions (37 °C, 5% CO_2_) for downstream experiments. Post-sort flow cytometric analysis confirmed >95% purity for eGFP-, synNotch- and mCherry-expressing populations. See Supplementary Table S7 for a list of sorted engineered cell lines and their respective characterizations.

### Expression and purification of eGFP ligand

A pET28a(+) vector encoding His-tagged eGFP was transformed into *E. coli* BL21(DE3) (Novagen, 69450). A single colony of freshly transformed bacteria was cultured in a shaking incubator at 140 rpm and 37 °C in 1 L of LB medium, supplemented with 50 µg/mL kanamycin (Carl Roth, T832). When the OD600 (optical density measured at 600 nm) of the culture reached ∼0.6-0.8, protein expression was induced by addition of IPTG (AppliChem, A1008) to a final concentration of 100 µM, and the culture was incubated overnight at 140 rpm and 16 °C. Cells were harvested by centrifugation at 8,000x *g* for 10 min. Pelleted cells were resuspended in 13.33 mL BugBuster (Merck, 70584) supplemented with 5 µL benzonase (Merck, E1014) and one protease inhibitor tablet (Thermo Scientific, A32965) and incubated for 1 h on a shaking plate. The lysate was centrifuged at 40,000x *g* for 20 min at 4 °C and the supernatant containing soluble His-tagged eGFP was collected for purification.

Protein purification was performed by Ni^2+^ affinity chromatography using Ni-NTA resin (QIAGEN, 30230) on a gravity column with a column volume of 2 mL. The column was equilibrated with binding buffer (20 mM Tris-Cl (pH 7.9), 500 mM NaCl, 5 mM imidazole), after which the lysate was loaded onto the column. The column was subsequently washed with binding buffer followed by washing buffer (20 mM Tris-Cl (pH 7.9), 500 mM NaCl, 20 mM imidazole). The His-tagged eGFP was eluted using elution buffer (20 mM Tris-Cl (pH 7.9), 500 mM NaCl, 100 mM imidazole), and eluted protein fractions were dialysed using SnakeSkin Dialysis Tubing (3.5 kDa MWCO, Thermo Scientific, 68035) overnight at room temperature against PBS (2L) to remove imidazole. Following dialysis, protein samples were concentrated using an Amicon Ultra Centrifugal Filter (3 kDa MWCO; Millipore, UFC9003) and stored at -80 °C until further use. Protein concentration was determined by spectrophotometry (NanoDrop One, Thermo Scientific) and purity was assessed by Coomassie-stained SDS-PAGE.

### *Receiver* cell activation

Sender-Receiver co-culture experiments were performed to activate synNotch signalling through cell-cell contact. HeLa Receiver cells were seeded in 24-well plates at a density of 5x10^4^ cells per well in growth medium approximately 24 h prior to co-culture. K562 *Sender* cells or K562 WT control cells were subsequently added to the Receiver monolayer in fresh growth medium. Different *Receiver*:*Sender* ratios were initially tested, and a ratio of 1:9 was used for all subsequent experiments. Co-cultures were maintained for 48 h under standard culture conditions (37 °C, 5% CO_2_). Following co-culture, wells were washed three times with PBS to remove suspension K562 cells prior to detachment. *Receiver* cells were subsequently analysed by flow cytometry as described above. During flow cytometry analysis, *Receiver* cells were identified based on constitutive mCherry expression.

To activate synNotch signalling using immobilized ligand, wells of 24-well plates were coated overnight at 4 °C with purified eGFP diluted in PBS. Coating concentrations were initially titrated, after which 3.2 µg eGFP in 300 µL PBS per well was used for all subsequent experiments. The following day, the coating solution was removed and wells were gently washed once with growth medium. HeLa *Receiver* cells were seeded onto the coated wells at a density of 5x10^4^ cells per well (unless explicitly stated otherwise) in growth medium and incubated under standard culture conditions (37 °C, 5% CO_2_) for 48 h prior to downstream analysis. For feedback loop experiments analysed using flow cytometry, Receiver cells were seeded at densities ranging from 8x10^4^ cells per well (interrogated at t = 24 h) to 0.5x10^4^ cells per well (interrogated at t = 144 h) for optimal cell density at time of analysis.

### Precursor miRNA transfection

HeLa Receivers were seeded at a density of 5x10^4^ cells per well in 24-well plates one day prior to transfection to achieve ∼70-80% confluency at the time of transfection. For transfection, miR-FF3 precursor miRNA (mirVana custom miRNA mimic; Invitrogen, 4464068; Assay ID: AKFARAD) or scrambled precursor miRNA (mirVana miRNA mimic, negative control #1; Invitrogen, 4464058) was diluted in Opti-MEM I Reduced Serum Medium (Gibco, 31985062) to final concentrations ranging from 0.01-100 nM. Transfections were performed using 1.25 µL Lipofectamine 2000 (Invitrogen, 11668019) per well in a total volume of 300 µL. After 6 h of incubation under standard culture conditions (37 °C, 5% CO_2_), the transfection mixture was replaced with fresh growth medium. *Receiver* cells were subsequently cultured for either an additional 72 h prior to analysis, or for 24 h before co-culture with K562 *Sender* cells (1:9) was initiated. Co-cultures were maintained for 48 h before analysis by flow cytometry as described above.

### Live-cell imaging

For miRNA transfection experiments, HeLa *Receiver* cells were seeded in glass-bottom 24-well imaging plates (Ibidi, 82426) at 2x10^4^ cells per well in regular growth medium and cultured overnight under standard culture conditions (37 °C, 5% CO_2_). The following day, cells were transfected with 0.1 nM precursor miRNA using Lipofectamine 2000 as described above, with a 5 h incubation period before replacement with fresh regular growth medium. Approximately 24 h after transfection, Sender-Receiver co-cultures were initiated by removing growth medium from the *Receiver* cells, washing once with PBS, and adding K562 *Sender* or WT cells to the *Receiver* monolayer at a *Receiver*:*Sender* ratio of 1:9 as described above; K562 cells were resuspended in phenol red-free imaging medium (Capricorn, DMEM-HXRXA), supplemented with 2 mM L-glutamine (Gibco, 25030), 10% (v/v) FBS (Greiner Bio-One, 758093) and 1% (v/v) penicillin-streptomycin (Gibco, 15140), that had been equilibrated at 37 °C with 5% CO_2_ for at least 2 h prior to use. Plates were then transferred to the microscope stage incubator for time-lapse imaging.

For feedback loop experiments, glass-bottom 24-well imaging plates (Ibidi, 82426) were coated overnight at 4 °C with purified eGFP ligand (3.2 µg per well in 300 µL PBS) as described above. The following day, *Receiver* cells were seeded at 1x10^4^ cells per well in pre-equilibrated phenol red-free imaging medium (as described above) on uncoated wells (control) or eGFP-coated wells, and transferred to the microscope stage incubator for time-lapse imaging.

Epifluorescence images were acquired using a 20x Plan Apochromat dry objective (NA 0.75; Nikon) on a Nikon Ti inverted microscope equipped with a stage incubator (Tokai-Hit), an LED illumination system (pE-4000, CoolLED), ET-DAPI (49000), ET-GFP (49002), and ET-mCherry (49008) filter cubes (Chroma), and a pco.edge 4.2 sCMOS camera (Excelitas). Image acquisition was controlled with µManager 1.4 software^63^. Cells were imaged at 3 h intervals for 45 h (for miRNA transfection experiments) or at 15 min intervals for 114 h (for feedback loop experiments) in the stage incubator set at 37 °C with 5% CO_2_.

### Image processing, cell segmentation, quantification and analysis of live-cell imaging experiments

All image processing and quantification were performed using FIJI/ImageJ (ImageJ 1.54)^64^. To correct for static background signal, a background reference image was generated for each individual imaging session from wells containing cells lacking tBFP and mCherry expression. The reference background image was generated by computing a minimum projection from these reference wells, followed by Gaussian smoothing (rolling ball with 2 pixel radius). The smoothed background image was then subtracted from all experimental time-lapse stacks. To measure cellular tBFP and mCherry intensities, the background-subtracted stacks were segmented to generate outlines of the *Receiver* cells based on the constitutive mCherry expression in these cells. Individual frames of the mCherry channels were segmented using Cellpose v2.2.3^65^, with a custom pretrained model that had been fine-tuned in-house by retraining a built-in Cellpose model on 17 manually annotated images, containing a total of 442 cells. Segmentation masks were then matched to the corresponding fluorescence stacks, and converted to regions of interest (ROIs) on a per frame basis. For each frame, the total tBFP fluorescence intensity per cell was quantified as mean tBFP intensity multiplied by the cellular area. The average tBFP intensity per cell was then calculated to generate a time-resolved trace of mean tBFP intensity per cell for every field of view. No images, frames, or positions were excluded from analysis except in cases of strong camera noise or segmentation failures. Fields of view were treated as technical replicates and were kept separate during image processing and quantification.

For downstream analysis, tBFP intensity traces were normalized per field of view by dividing each trace by the median tBFP intensity from a defined reference time range (t = 45 h for miRNA transfection experiments, and t = 47.5-49.5 h for feedback loop experiments). Each position was normalized to a control condition (‘4xTS-FF3 + *Senders* with scrambled’ for transfection experiments, and ‘4xTS-FF3 + miR-FF3 + eGFP’ for feedback loop experiments) acquired during the same imaging session to correct for day-to-day variability. For long term imaging on eGFP-coated wells, normalized time series were smoothed using a rolling median filter with a window size of seven time points (current frame ± 3 frames) to reduce technical noise, such as camera noise, during imaging. Final traces per experimental condition were generated by averaging all smoothed, normalized traces per condition. Positions were pooled across multiple fields of view and imaging sessions.

For visualization of cellular outlines in representative images and movies, segmented cell masks were rendered using Trackmate v7.14.0^66^. Segmented objects were detected using the label image detector on the segmentation channel and visualized as spot overlays.

### Quantitative RT-PCR analysis of miRNA expression

HeLa miR-FF3 *Receiver* cells (1x10^5^ cells per well in a 6-well plate) were activated on eGFP-coated plates for 72 h as described above, after which cells were harvested and split for flow cytometry and RNA isolation. Cell pellets (≤ 5x10^5^ cells per sample) were snap-frozen and stored at -80 °C prior to RNA extraction. Total RNA, including miRNAs, was isolated using the miRNeasy Tissue/Cells Advanced Micro Kit (QIAGEN, 217684) according to the manufacturer’s instructions. Total RNA was quantified by spectrophotometry (NanoDrop One, Thermo Scientific) and RNA purity was assessed using A260/A280 ratios. Reverse transcription was performed using the miRCURY LNA RT Kit (QIAGEN, 339340). For each sample, 100 ng total RNA was used in a 10 µL reaction and reverse transcribed according to manufacturer’s protocol. Quantitative PCR was performed using the miRCURY LNA SYBR Green PCR Kit (QIAGEN, 339345) on a QuantStudio 6 Pro Real-Time PCR System (Applied Biosystems) using 384-well plates (Applied Biosystems, 4309849). cDNA samples were diluted 1:20 in 0.02% BSA (NEB, B9200) in TE buffer (IDT, 11-05-01-05) prior to amplification. miR-FF3 expression levels were quantified using a custom miRCURY LNA PCR Assay (QIAGEN, 339317, ID: YCP2152773). An hsa-miR-103a-3p miRCURY LNA PCR Assay (QIAGEN, 339306, ID: YP00204063) was used as an endogenous normalization control. Amplification of n=3 independent replicates (seeded, lysed and analysed in parallel) was performed in technical triplicates, of which Ct values were averaged prior to further analysis. PCR efficiency of both primer assays was determined using a serial dilution of cDNA generated from a positive control sample with elevated miR-FF3 expression (Supplementary Table S6 and Fig. S22). Ct values were determined using QuantStudio Design and Analysis v2 software (Applied Biosystems). Relative miRNA expression levels were calculated using the Pfaffl method^67^, incorporating the experimentally determined primer efficiencies, and normalized to a non-activated *Receiver* control.

### SEAP reporter assay

HeLa miR-FF3 *Receiver* cells were seeded on uncoated or eGFP-coated 24-well plates (3.2 µg eGFP per well) at a density of 2x10^4^ cells per well in growth medium and cultured overnight under standard culture conditions (37 °C, 5% CO_2_). The following day, cells were transiently transfected with a SEAP reporter plasmid using Lipofectamine 2000 (Invitrogen, 11668019). For each well, 50 ng plasmid DNA was transfected using 1.25 µL Lipofectamine 2000 diluted in 300 µL Opti-MEM I Reduced Serum Medium (Gibco, 31985062). After 5 h of incubation under standard culture conditions (37 °C, 5% CO_2_), the transfection mixture was replaced with fresh growth medium. *Receiver* cells were subsequently cultured for an additional 48 h prior to analysis of SEAP activity in the culture medium.

For the SEAP reporter assay, cell culture medium was harvested from the wells, cell debris was removed by means of centrifugation, and medium was heat-inactivated at 65 °C for 30 min to inactivate endogenous phosphatases. SEAP concentration in cell culture medium was quantified in terms of absorbance increase due to the hydrolysis of p-Nitrophenylphosphate (pNPP) to p-Nitrophenol (pNP). SEAP Sample Dilution Buffer (10x; Novus Biologicals, NBP2-25285, IMK-515-6) was used to dilute medium samples 1:2. SEAP pNPP substrate was prepared by dissolving a 5 mg pNPP Substrate Tablet (Thermo Scientific, 34047) in Diethanolamine Substrate Buffer (5x; Thermo Scientific, 34064). 10 µL diluted sample was then added to 10 µL MilliQ water and 100 µL pNPP substrate (1 mg/mL) in a transparent 96-well plate. Absorbance values were measured every 30 s at 405 nm for 60 min using a Tecan Spark 10M platereader (Tecan) at 25 °C. To determine sample SEAP concentration, a calibration curve was constructed by titrating known concentrations of the hydrolysed pNPP product pNP (Merck, 1048; Supplementary Fig. S24). Absorbance units are converted to amount of substrate conversion and SEAP activity is calculated from the slope of the time trace, expressed in units per litre (U/L). One unit is defined as the amount of enzyme that converts 1 µmole pNPP in 5 µL cell culture medium in 1 min at 25 °C.

### Mathematical modelling

The synNotch-miRNA feedback circuit was mathematically described using a system of ten coupled ordinary differential equations (Supplementary Notes). The model employs mass-action kinetics for most biochemical reactions and Hill kinetics for GAL4-VP64-mediated transcriptional activation. Model parameters were derived from literature values for similar biological systems. Parameters not directly available from literature were adjusted within biologically plausible ranges to qualitatively reproduce experimentally observed dynamics (Supplementary Notes). The base synNotch model without miRNA feedback was obtained by setting all miRNA-related rate constants to zero, reducing the system to eight equations. The ODE system was implemented in Python using the SciPy library. Numerical integration was performed using the ‘solve_ivp’ function with the implicit ‘Radau’ method suited for stiff biochemical reaction networks. Initial conditions were determined by simulating the system in the absence of GFP ligand until all species reached steady state concentrations (t = 120 h). To simulate ligand-induced activation, GFP ligand was introduced at t = 0 h with a concentration of 16.5 nM and maintained constant throughout the simulation (Supplementary Notes). Simulations were run over 120-hour time courses with 500 evaluation points. For parameter robustness analysis, Monte Carlo simulations (10,000 iterations per model variant) were performed with each parameter (except ‘H_GAL’) independently and uniformly sampled within ±5% of nominal values. For feedback ratio analysis across parameter space, the model parameters (except ‘H_GAL’) were independently and uniformly sampled over a twofold range (0.5x to 2x nominal values) in 10,000 Monte Carlo simulations.

### Data analysis and statistics

Data analysis and visualization were performed using Python (v3.9.13; with packages NumPy, SciPy, pandas, and matplotlib) and MATLAB (R2021b; The Mathworks Inc.). Statistical analyses were performed using GraphPad Prism (v11.0.0; GraphPad Software). Flow cytometry data were analysed using FlowJo (v10.8; BD Life Sciences), and microscopy data were processed and quantified using FIJI/ImageJ (ImageJ 1.54)^64^, Cellpose (v2.2.3)^65^, and TrackMate (v7.14.0)^66^ as described above. Data are presented as mean with individual data points representing independent replicates of wells seeded, transfected and analysed in parallel, unless explicitly stated otherwise. Technical replicates were used for qPCR analyses, and included for microscopy experiments as FOVs. For experiments with nested replicate structure, statistical comparisons were performed using appropriate mixed-effects models with well or experimental day treated as random effects, as indicated in the figure legends and Supplementary Tables S10 and S11. For comparisons between two groups, two-tailed unpaired t-tests were used; for comparisons among three or more groups, one-way ANOVA with Tukey’s multiple comparisons test was used. For qPCR, fold changes were calculated using the Pfaffl method^67^ and statistical comparisons were performed using Prism’s lognormal t-test. For ratio-based metrics derived from fluorescence measurements, uncertainty was corrected by propagation of standard deviations. A significance threshold of p < 0.05 was used. No statistical method was used to predetermine sample size. Experiments were not randomized and investigators were not blinded. No data were excluded from analyses unless necessitated by excessive cell death or technical failure, such as strong camera noise or segmentation failure.

## Data availability

Datasets generated and/or analysed in the present study have been deposited in the Zenodo database at https://zenodo.org/records/20343345 (https://doi.org/10.5281/zenodo.20343345).

## Materials availability

Unique biological materials, such as plasmids and cell lines are available from the corresponding author upon request. Plasmids engineered and used in the present study are also available from Addgene (see Supplementary Table S2). ODE model code and simulation scripts have been deposited in the Zenodo database at https://zenodo.org/records/20343345 (https://doi.org/10.5281/zenodo.20343345).

## Acknowledgements

The authors thank David Schrijver for his help with fluorescence-activated cell sorting of the *Sender* cell line, Jesse Lentjes for her help with expression and purification of the eGFP ligand, Angelina Yurchenko for her help with data analysis, and Indra van Zundert for her help with microscopy experiments. The authors would also like to thank Wendell Lim for providing the original synNotch receptor and reporter plasmids, Didier Trono for providing the plasmids necessary for lentivirus production, and Alan Cochrane for providing the SEAP reporter plasmid. This work was supported by the European Research Council (ERC; project no. 101000199 AMIGA, awarded to T.F.A.d.G.), the Netherlands Organization for Scientific Research (NWO) Gravitation programme IMAGINE! (project no. 24.005.009, awarded to L.C.K.), and the EWUU Alliance (TU/e, WUR, UU, UMC Utrecht) through the Centre for Living Technologies.

## Author contributions

B.L.N., A.M.P., and T.J.M. designed the study, performed experiments, and analysed the data. B.L.N. wrote the manuscript. B.L.N. and T.J.M. established cell lines and performed related experiments. W.N. and T.J.M. performed microscopy experiments. W.N. and L.C.K. analysed microscopy data and provided key insights with regard to live-cell imaging. D.R.G., K.S.L., and K.E.G. provided key insights with regard to cell experiments. E.G.G., A.F.v.G., and B.L.N. designed the mathematical model and ran simulations. A.M.P., T.J.M., and W.D. designed plasmids and assisted in cloning and cell experiments. K.E.G. and L.C.K. provided critical feedback on experiments and revised the manuscript. T.F.A.d.G. conceived, designed and supervised the study, analysed the data, and wrote the manuscript.

---

**Extended Data Figure 1:**
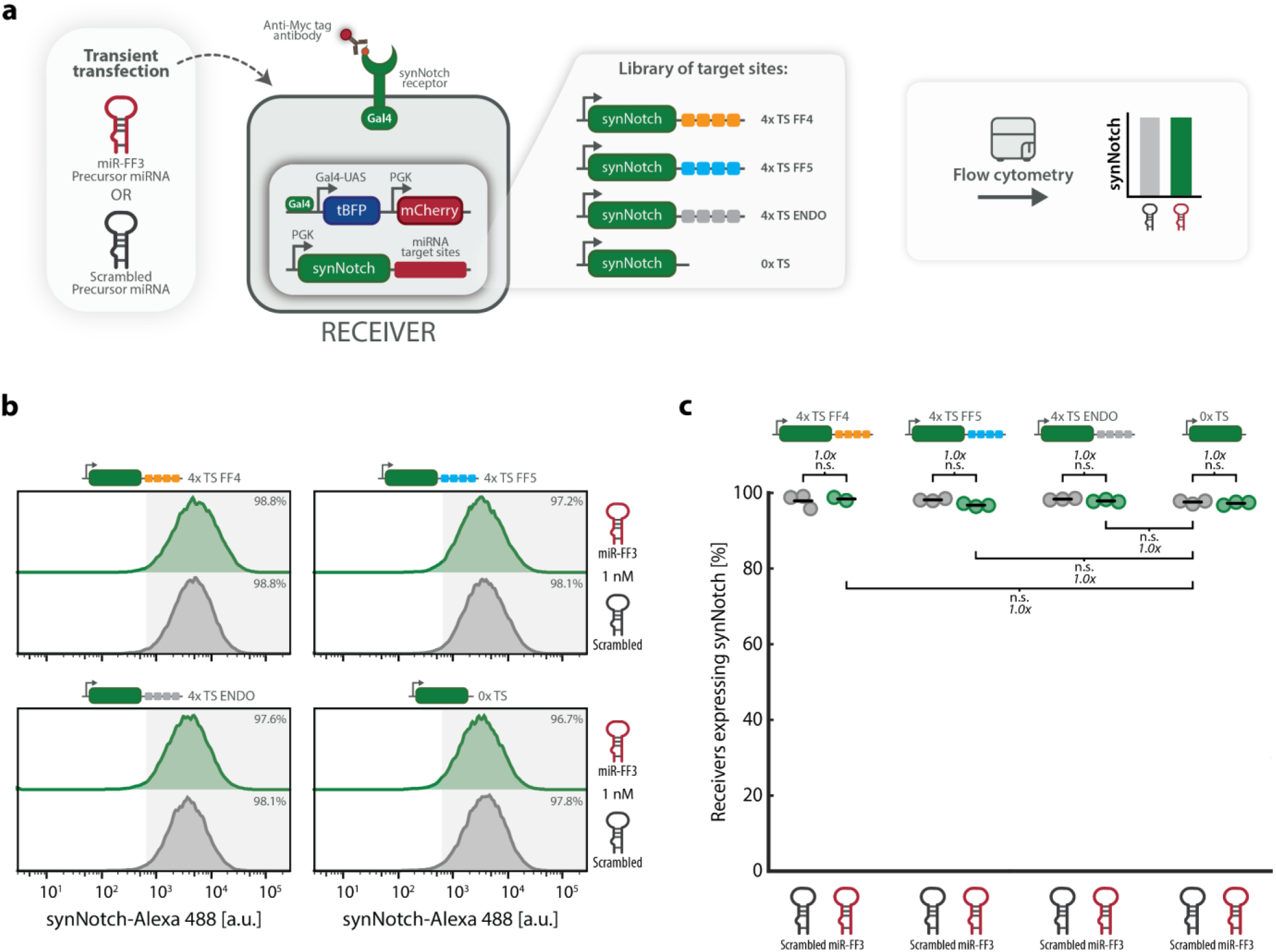
The miR-FF3 miRNA-target site pair functions as an orthogonal regulatory module for synNotch receptor control. **a** *Receiver* cells containing four alternative, non-complementary miRNA target sites (FF4, FF5^34^, or a heterologous sequence described by Endo et al. (2019)^36^; Supplementary Table S4-S5, and Supplementary Fig. S10-S11) are transiently transfected with miR-FF3 or scrambled miRNA precursor. Following transfection, cells are analysed for synNotch expression using flow cytometry. Schematics (right) illustrate the expected lack of repression of synNotch for non-complementary miRNA-target site pairs. **b** SynNotch expression 72 h post-transfection. Flow cytometry histograms show synNotch expression (anti-Myc AF488 staining) in cells hosting 4xTS-FF4, 4xTS-FF5, 4xTS-ENDO or 0xTS, transiently transfected with 0.1 nM miR-FF3 (green) or scrambled (grey) precursor miRNA. Shaded area indicates synNotch-positive cells (threshold defined by HeLa WT control). **c** Percentage of synNotch-positive cells (anti-Myc AF488 staining), quantified by flow cytometry analysis, in cells hosting 4xTS-FF4, 4xTS-FF5, 4xTS-ENDO or 0xTS, transiently transfected with 0.1 nM scrambled (grey) or miR-FF3 (green) precursor miRNA. Horizontal lines indicate mean expression; individual data points represent independent replicates seeded, transfected, and analysed in parallel (n=3 independent wells per condition). Fold change and significance (one-way ANOVA with Tukey’s multiple comparisons test) are noted above or below data points (Supplementary Table S9). n.s.: p > 0.05, *p ≤ 0.05, **p ≤ 0.01, ***p ≤ 0.001.

**Extended Data Figure 2:**
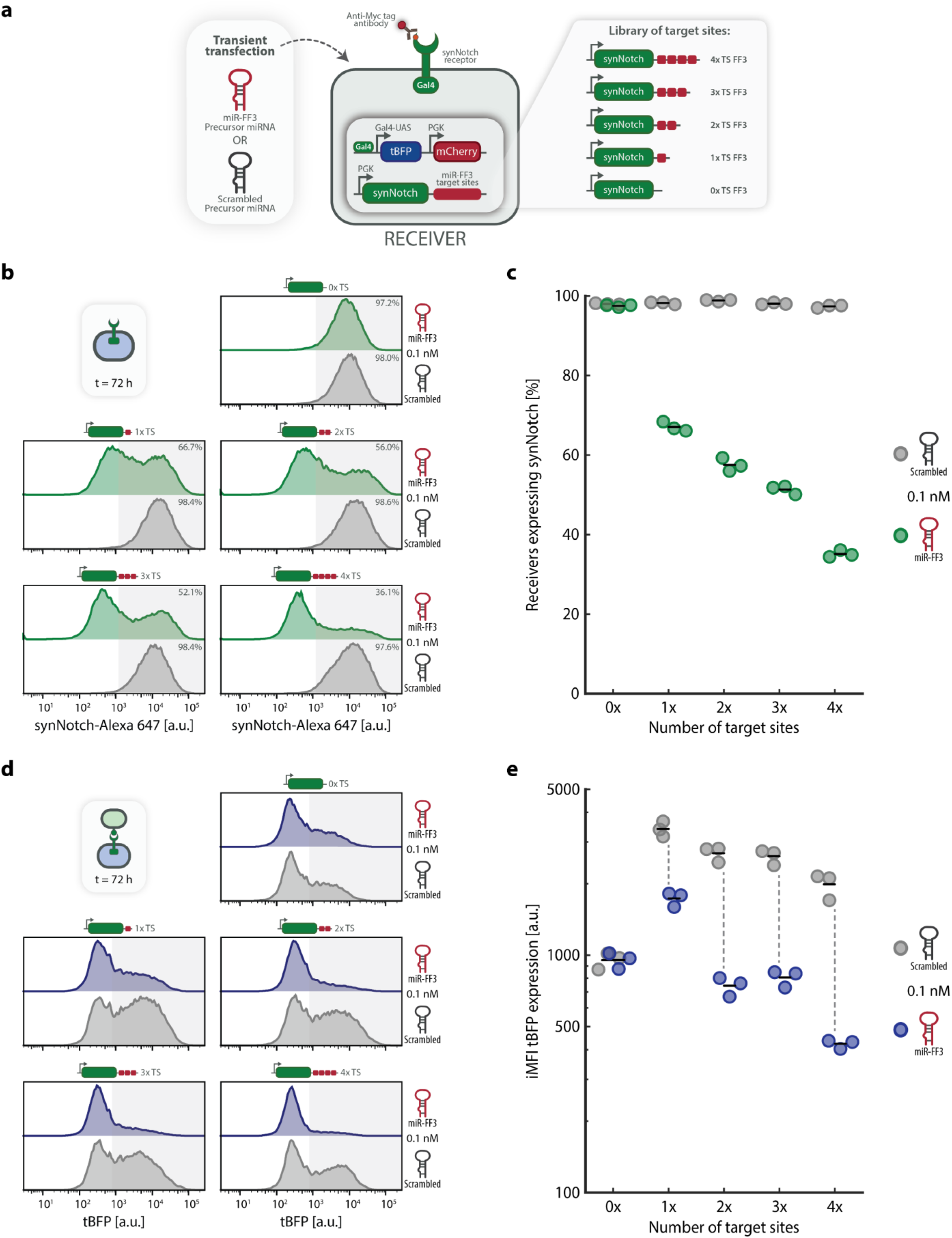
SynNotch repression and downstream activation are tunable and correlate with the number of miRNA target sites. **a** Experimental scheme for testing target site-dependent repression of synNotch and tBFP expression. A library of engineered *Receiver* cell lines, containing 0, 1, 2, 3 or 4 miR-FF3 target sites (TS; Supplementary Fig. S7-S8 and S17-S18), is transiently transfected with 0.1 nM complementary (miR-FF3) or scrambled (negative control) mirVana™ precursor miRNA (Methods). Following transfection, cells are either analysed for synNotch expression (anti-Myc AF647 staining) or activated and subsequently analysed for tBFP expression by flow cytometry. **b** Representative histograms showing the effect of precursor miRNA on the expression of synNotch, 72 h post-transfection. Shaded area indicates synNotch-positive cells (percentage also shown in top-right corner). **c** Percentages of *Receiver* cells expressing synNotch as used in Fig. 3b to calculate repression of synNotch expression. Horizontal lines indicate mean expression; individual data points represent independent replicates seeded, transfected, and analysed in parallel (n=3 independent wells per condition). **d** Representative histograms showing the effect of precursor miRNA transfection on the expression of tBFP. *Receiver* cells were transiently transfected with precursor miRNA, incubated for 24 h, and subsequently co-cultured for 48 h with K562 eGFP+ *Sender* cells. Shaded area indicates tBFP-positive cells (threshold defined by HeLa WT control). **e** Integrated Median Fluorescence Intensity (iMFI) of tBFP expression of *Receiver* cells as used in Fig. 3c to calculate repression of tBFP expression. Horizontal lines indicate mean expression; individual data points represent independent replicates seeded, transfected, and analysed in parallel (n=3 independent wells per condition).

**Extended Data Figure 3:**
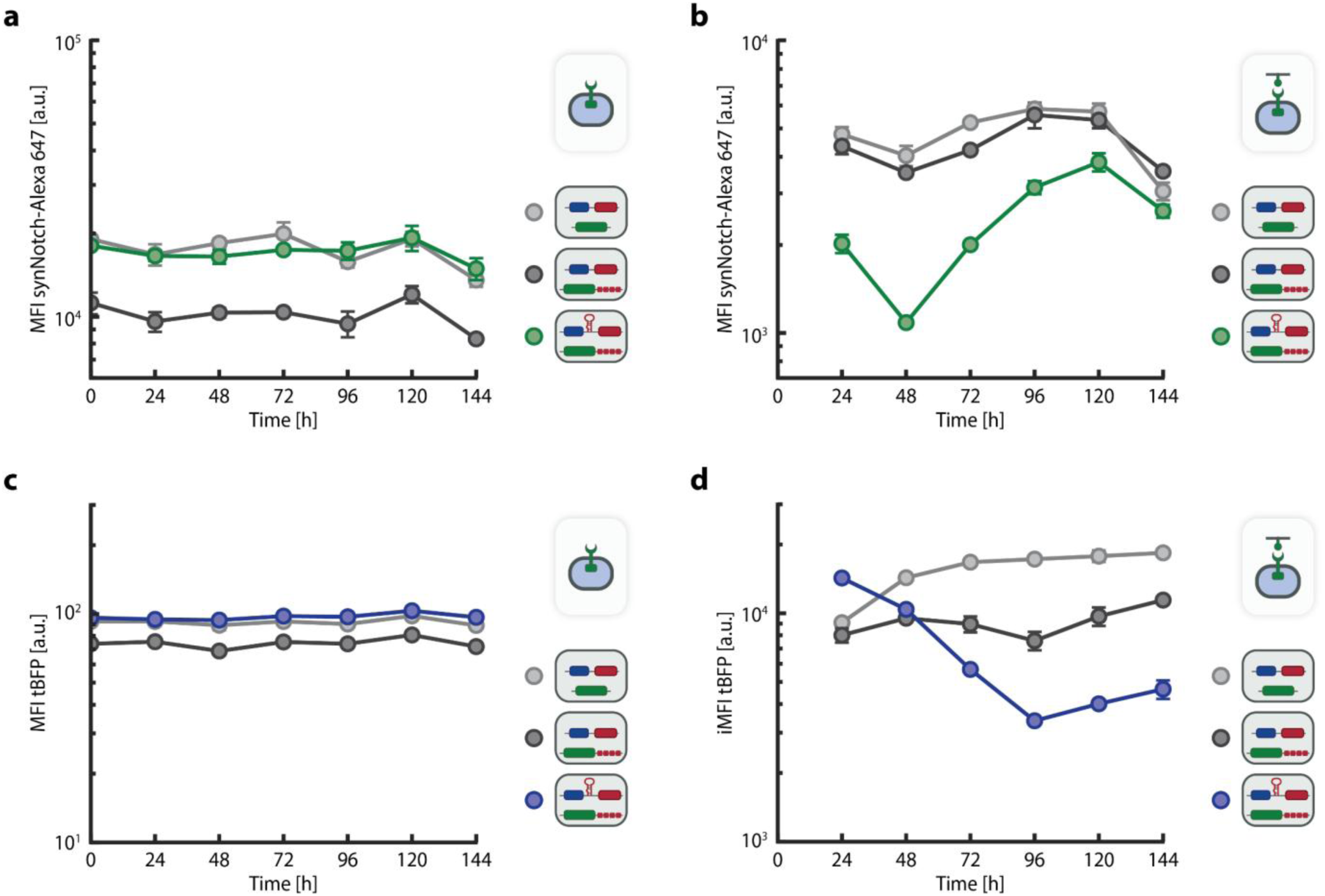
Expression of synNotch and tBFP in three different *Receiver* cell lines cultured on uncoated and eGFP-coated plates. **a-d** Time-course analysis of synNotch and tBFP expression used to calculate synNotch repression and tBFP activation as shown in Fig. 5b and 5c, respectively. Three types of *Receiver* cell lines (light grey: without feedback loop components, dark grey: without miR-FF3 gene and with miR-FF3 target sites, green/blue: with miR-FF3 gene and target sites) were seeded at t = 0 h on uncoated **(a/c)** or eGFP-coated plates **(b/d)**. Expression of synNotch and tBFP over time were quantified by flow cytometry analysis, with measurements every 24 hours. Mean synNotch MFI **(a-b)**, tBFP MFI **(c)** or tBFP iMFI **(d),** and standard deviations are shown for n = 3 replicates per timepoint (independent wells seeded and analysed in parallel). For non-activated samples, baseline tBFP expression was uniformly low and therefore quantified using the MFI. Using MFI for non-activated samples avoids instability caused by near-zero fractions of tBFP-positive cells, for which iMFI values approach zero and normalization becomes unreliable. For representative flow cytometry histograms of each condition and timepoint, see Supplementary Fig. S28-S29.

