## Supplementary Information for "Engineered microRNA feedback circuits enable tunable and autonomous control of synthetic receptor activity"

**This document includes:**

- Supplementary Tables S1-S11
- Supplementary Figures S1-S32
- Captions for Supplementary Videos S1-S2
- Supplementary Notes

### Supplementary Tables

**Supplementary Table S1: Plasmid backbones used in the present study.** Overview of plasmid backbones, their function and corresponding source.

| Plasmid ID | Description | Source |
|---|---|---|
| pHR_SFFV | Second generation lentiviral transfer plasmid for constitutive expression of genes in mammalian cells (SFFV promoter) | Morsut et al. (2016)[1] |
| pHR_PGK | Second generation lentiviral transfer plasmid for constitutive expression of genes in mammalian cells (PGK promoter) | Morsut et al. (2016)[1] |
| pHR_5xGal4UAS | Second generation lentiviral transfer plasmid for Gal4-VP64 inducible expression of genes in mammalian cells (Gal4-UAS promoter) | Morsut et al. (2016)[1] |
| pCMVR8.74 | Second generation lentiviral packaging plasmid | Unpublished work;<br>Gift from Didier Trono<br>(Addgene plasmid # 22036 ;<br>http://n2t.net/addgene:22036 ;<br>RRID:Addgene_22036) |
| pMD2.G | VSV-G envelope expressing plasmid | Unpublished work;<br>Gift from Didier Trono<br>(Addgene plasmid # 12259 ;<br>http://n2t.net/addgene:12259 ;<br>RRID:Addgene_12259) |
| pBluescript_CMV_SEAP | Mammalian expression vector for constitutive expression (CMV promoter) of SEAP reporter gene | Unpublished work;<br>Gift from Alan Cochrane<br>(Addgene plasmid # 24595 ;<br>http://n2t.net/addgene:24595 ;<br>RRID:Addgene_24595) |

**Supplementary Table S2: Engineered plasmids used in the present study.** Overview of engineered plasmids, including plasmid backbone and gene insert. Plasmids generated in this study have been deposited to Addgene (Plasmid IDs #249359 – #249367).

| Plasmid | Plasmid backbone | Gene insert | Notes |
|---|---|---|---|
| pHR_SFFV_eGFP-ligand | pHR_SFFV | eGFP-ligand | Gift from Wendell Lim<br>(Addgene plasmid # 79129 ;<br>http://n2t.net/addgene:79129 ;<br>RRID:Addgene_79129)[1] |
| pHR_PGK_LaG17_synNotch_<br>Gal4-VP64 | pHR_PGK | LaG17_synNotch_<br>Gal4-VP64 | Gift from Wendell Lim<br>(Addgene plasmid # 79127 ;<br>http://n2t.net/addgene:79127 ;<br>RRID:Addgene_79127)[1] |
| pHR_Gal4-UAS_tBFP_<br>PGK_mCherry | pHR_<br>5xGal4UAS | tBFP_PGK_mCherry | Gift from Wendell Lim<br>(Addgene plasmid # 79130 ;<br>http://n2t.net/addgene:79130 ;<br>RRID:Addgene_79130)[1] |
| pHR_PGK_LaG17_synNotch_<br>Gal4-VP64_1xTS-FF3 | pHR_PGK | LaG17_synNotch_<br>Gal4-VP64_1xTS-FF3 | Deposited to Addgene;<br>Plasmid ID #249359 |
| pHR_PGK_LaG17_synNotch_<br>Gal4-VP64_2xTS-FF3 | pHR_PGK | LaG17_synNotch_<br>Gal4-VP64_2xTS-FF3 | Deposited to Addgene;<br>Plasmid ID #249360 |
| pHR_PGK_LaG17_synNotch_ | pHR_PGK | LaG17_synNotch_ | Deposited to Addgene; |

| | | | |
|---|---|---|---|
| Gal4-VP64_3xTS-FF3 | | Gal4-VP64_3xTS-FF3 | Plasmid ID #249361 |
| pHR_PGK_LaG17_synNotch_ Gal4-VP64_4xTS-FF3 | pHR_PGK | LaG17_synNotch_ Gal4-VP64_4xTS-FF3 | Deposited to Addgene; Plasmid ID #249362 |
| pHR_PGK_LaG17_synNotch_ Gal4-VP64_4xTS-FF4 | pHR_PGK | LaG17_synNotch_ Gal4-VP64_4xTS-FF4 | Deposited to Addgene; Plasmid ID #249363 |
| pHR_PGK_LaG17_synNotch_ Gal4-VP64_4xTS-FF5 | pHR_PGK | LaG17_synNotch_ Gal4-VP64_4xTS-FF5 | Deposited to Addgene; Plasmid ID #249364 |
| pHR_PGK_LaG17_synNotch_ Gal4-VP64_4xTS-ENDO | pHR_PGK | LaG17_synNotch_ Gal4-VP64_4xTS-ENDO | Deposited to Addgene; Plasmid ID #249365 |
| pHR_Gal4-UAS_tBFP_ miR30-FF3_PGK_mCherry | pHR_ 5xGal4UAS | tBFP_miR30-FF3_ PGK_mCherry | Deposited to Addgene; Plasmid ID #249366 |
| pBluescript_CMV_SEAP | pBluescript_ CMV_SEAP | - | Gift from Alan Cochrane (Addgene plasmid # 24595 ; http://n2t.net/addgene:24595 ; RRID:Addgene_24595) |
| pBluescript_CMV_SEAP_4xTS-FF3 | pBluescript_ CMV_SEAP | 4xTS-FF3 | Deposited to Addgene; Plasmid ID #249367 |

**Supplementary Table S3: DNA sequence of microRNA (miRNA) gene.** Overview of miRNA gene sequence incorporated in reporter plasmid for inducible miRNA expression. For context and position in reporter plasmid, also see Supplementary Fig. S2. MiR-FF3 guide strand is shown underlined.

| miRNA construct | DNA Sequence (5' to 3') | Source |
|---|---|---|
| **miR30-FF3** | TAGGCGCGCCATAACTTCCGGCCGCAAGCCTTGTTAAGTGCTCGC TTCGGCAGCACATATACTATGTTGAATGAGGCTTCAGTACTTTAC AGAATCGTTGCCTGCACATCTTGGAAACACTTGCTGGGATTACTT CTTCAGGTTAACCCAACAGAAGGCTCGAGAAGGTATATTGCTGTT GACAGTGAGCGCACGATATGGGCTGAATACAAATAGTGAAGCCAC AGATGTATTTGTATTCAGCCCATATCGTTTGCCTACTGCCTCGGA ATTCAAGGGGCTACTTTAGGAGCAATTATCTTGTTTACTAAAACT GAATACCTTGCTATCTCTTTGATACATTTTTACAAAGCTGAATTA AAATGGTATAAATTAAATCACTTTTT | Leisner et al. (2010)[2], Stegmeier et al. (2005)[3], Venniro et al. (2020)[4] |

**Supplementary Table S4: DNA sequences of miRNA target sites.** Overview of miRNA target site sequences used in this study.

| miRNA target site | DNA Sequence (5' to 3') | Source |
|---|---|---|
| **FF3** | AACGATATGGGCTGAATACAAA | Leisner et al. (2010)[2] |
| **FF4** | CCGCTTGAAGTCTTTAATTAAA | Leisner et al. (2010)[2] |
| **FF5** | AAGCACTCTGATTTGACAATTA | Leisner et al. (2010)[2] |
| **ENDO** | CGAACGGGCACGCTGACAATTC | Endo et al. (2019)[5] |

**Supplementary Table S5: DNA sequences of miRNA target site regions.** Overview of sequences of full miRNA target site regions incorporated in 3' UTR of a synNotch transcript. For context and position in receptor plasmid, also see Supplementary Fig. S3, S10 and S17.

| Target site construct | DNA Sequence (5' to 3') |
|---|---|
| **1xTS FF3** | GATCCTTGACTTGCGGGATCCAACGATATGGGCTGAATACAAACCCGG |
| **2xTS FF3** | GATCCTTGACTTGCGGGATCCAACGATATGGGCTGAATACAAACGACAACGATATGGG<br>CTGAATACAAACCCGG |
| **3xTS FF3** | GATCCTTGACTTGCGGGATCCAACGATATGGGCTGAATACAAACGACAACGATATGGG<br>CTGAATACAAACCCTAACGATATGGGCTGAATACAAACCCGG |
| **4xTS FF3** | GATCCTTGACTTGCGGGATCCAACGATATGGGCTGAATACAAACGACAACGATATGGG<br>CTGAATACAAACCCTAACGATATGGGCTGAATACAAAACCAAACGATATGGGCTGAAT<br>ACAAACCCGG |
| **4xTS FF4** | GATCCTTGACTTGCGGGATCCCCGCTTGAAGTCTTTAATTAAACGACCCGCTTGAAGT<br>CTTTAATTAAACCCTCCGCTTGAAGTCTTTAATTAAAACCACCGCTTGAAGTCTTTAA<br>TTAAACCCGG |
| **4xTS FF5** | GATCCTTGACTTGCGGGATCCAAGCACTCTGATTTGACAATTACGACAAGCACTCTGA<br>TTTGACAATTACCCTAAGCACTCTGATTTGACAATTAACCAAAGCACTCTGATTTGAC<br>AATTACCCGG |
| **4xTS ENDO** | GATCCTTGACTTGCGGGATCCCGAACGGGCACGCTGACAATTCCGACCGAACGGGCAC<br>GCTGACAATTCCCCTCGAACGGGCACGCTGACAATTCACCACGAACGGGCACGCTGAC<br>AATTCCCCGG |

**Supplementary Table S6: qPCR primers used in this study.** Overview of qPCR primer pairs (miRCURY LNA miRNA PCR Assay; Methods) used to quantify miR-FF3 expression in *Receiver* cell lines. For PCR efficiency of primer pairs, also see Supplementary Fig. S22.

| Target | Function | PCR Efficiency | Manufacturer | ID |
|---|---|---|---|---|
| **hsa-miR-103a-3p** | Reference gene | 101.5% | QIAGEN (Cat #339306) | YP00204063 |
| **miR-FF3-3p** | Target gene | 88.2% | QIAGEN (Cat #339317) | YCP2152773 |

**Supplementary Table S7: List of cell lines engineered in this study.** Overview of engineered cell lines that have been stably transduced using a lentiviral transduction system (Methods). For details on transfer plasmids used to make the lentivirus, also see Supplementary Table S2.

| Cell line | Transfer plasmid 1 | Transfer plasmid 2 | Characterization |
|---|---|---|---|
| **K562 *Sender* (+eGFP ligand)** | pHR_SFFV_eGFP-ligand | n.a. | Supplementary Fig. S1 |
| **HeLa *Receiver*** | pHR_PGK_LaG17_synNotch_ Gal4-VP64 | pHR_Gal4-UAS_tBFP_ PGK_mCherry | Supplementary Fig. S8 |
| **HeLa *Receiver*** *(High expression)* | pHR_PGK_LaG17_synNotch_ Gal4-VP64 | pHR_Gal4-UAS_tBFP_ PGK_mCherry | Supplementary Fig. S20a |
| **HeLa *Receiver* +1xTS-FF3** | pHR_PGK_LaG17_synNotch_ Gal4-VP64_1xTS-FF3 | pHR_Gal4-UAS_tBFP_ PGK_mCherry | Supplementary Fig. S18a |
| **HeLa *Receiver* +2xTS-FF3** | pHR_PGK_LaG17_synNotch_ Gal4-VP64_2xTS-FF3 | pHR_Gal4-UAS_tBFP_ PGK_mCherry | Supplementary Fig. S18b |
| **HeLa *Receiver* +3xTS-FF3** | pHR_PGK_LaG17_synNotch_ Gal4-VP64_3xTS-FF3 | pHR_Gal4-UAS_tBFP_ PGK_mCherry | Supplementary Fig. S18c |

| | | | |
|---|---|---|---|
| **HeLa *Receiver* +4xTS-FF3** | pHR_PGK_LaG17_synNotch_ Gal4-VP64_4xTS-FF3 | pHR_Gal4-UAS_tBFP_ PGK_mCherry | Supplementary Fig. S7 |
| **HeLa *Receiver* +4xTS-FF3** *(High expression)* | pHR_PGK_LaG17_synNotch_ Gal4-VP64_4xTS-FF3 | pHR_Gal4-UAS_tBFP_ PGK_mCherry | Supplementary Fig. S26 |
| **HeLa *Receiver* +4xTS-FF4** | pHR_PGK_LaG17_synNotch_ Gal4-VP64_4xTS-FF4 | pHR_Gal4-UAS_tBFP_ PGK_mCherry | Supplementary Fig. S11a |
| **HeLa *Receiver* +4xTS-FF5** | pHR_PGK_LaG17_synNotch_ Gal4-VP64_4xTS-FF5 | pHR_Gal4-UAS_tBFP_ PGK_mCherry | Supplementary Fig. S11b |
| **HeLa *Receiver* +4xTS-ENDO** | pHR_PGK_LaG17_synNotch_ Gal4-VP64_4xTS-ENDO | pHR_Gal4-UAS_tBFP_ PGK_mCherry | Supplementary Fig. S11c |
| **HeLa *Receiver* +miR30-FF3** *(High expression)* | pHR_PGK_LaG17_synNotch_ Gal4-VP64 | pHR_Gal4-UAS_tBFP_ miR30-FF3_PGK_mCherry | Supplementary Fig. S19 |
| **HeLa *Receiver* +4xTS-FF3 +miR30-FF3** *(High expression)* | pHR_PGK_LaG17_synNotch_ Gal4-VP64_4xTS-FF3 | pHR_Gal4-UAS_tBFP_ miR30-FF3_PGK_mCherry | Supplementary Fig. S5 |

**Supplementary Table S8: p-values, t-values and degrees of freedom for unpaired, two-tailed t-tests.** Overview of fold change, p-values, t-values and degrees of freedom (df) for unpaired, two-tailed t-tests, indicating the compared groups and corresponding Figure. n.s.: p > 0.05, *p ≤ 0.05, **p ≤ 0.01, ***p ≤ 0.001.

| **Figure** | **Compared groups** | **Fold change** | **p-value summary** | **p-value** | **t-value** | **df** |
|---|---|---|---|---|---|---|
| S9b | 0xTS with scrambled vs. 0xTS with miR-FF3 | 1.0 | n.s. | 0.0569 | 2.652 | 4 |
| S9b | 4xTS-FF3 with scrambled vs. 4xTS-FF3 with miR-FF3 | 2.8 | *** | <0.001 | 114.8 | 4 |
| 2d | 0xTS + *Senders* with scrambled vs. 0xTS + *Senders* with miR-FF3 | 1.0 | n.s. | 0.9794 | 0.02753 | 4 |
| 2d | 4xTS-FF3 + *Senders* with scrambled vs. 4xTS-FF3 + *Senders* with miR-FF3 | 4.7 | *** | <0.001 | 10.95 | 4 |
| 4b | miR-FF3 *Receivers* vs. miR-FF3 *Receivers* + eGFP | 68.9 | *** | <0.001 | 20.49 (lognormal t-test) | 4 |

**Supplementary Table S9: F-values, p-values and degrees of freedom for one-way ANOVAs.** Overview of F-values, p-values and degrees of freedom (residual df) for one-way ANOVAs, with Tukey's multiple comparison test, indicating the compared groups and corresponding Figure. n.s.: p > 0.05, *p ≤ 0.05, **p ≤ 0.01, ***p ≤ 0.001.

| Figure | Compared groups (ANOVA) | F-value | p-value summary (ANOVA) | Compared groups (multiple comparisons) | Adjusted p-value (multiple comparisons) | df |
|---|---|---|---|---|---|---|
| Ext. Data 1c | 4xTS-FF4 with scrambled (A) vs. 4xTS-FF4 with miR-FF3 (B) vs. 4xTS-FF5 with scrambled (C) vs. 4xTS-FF5 with miR-FF3 (D) vs. 4xTS-ENDO with scrambled (E) vs. 4xTS-ENDO with miR-FF3 (F) vs. 0xTS with scrambled (G) vs. 0xTS with miR-FF3 (H) | 1.574 | n.s. (p=0.2178) | A-B | n.s. (p=0.9947) | 15 |
| | | | | A-C | n.s. (p=0.9998) | |
| | | | | A-D | n.s. (p=0.6089) | |
| | | | | A-E | n.s. (p=0.9973) | |
| | | | | A-F | n.s. (p>0.9999) | |
| | | | | A-G | n.s. (p=0.9984) | |
| | | | | A-H | n.s. (p=0.9389) | |
| | | | | B-C | n.s. (p>0.9999) | |
| | | | | B-D | n.s. (p=0.3231) | |
| | | | | B-E | n.s. (p>0.9999) | |
| | | | | B-F | n.s. (p=0.9890) | |
| | | | | B-G | n.s. (p=0.9018) | |
| | | | | B-H | n.s. (p=0.6659) | |
| | | | | C-D | n.s. (p=0.3696) | |
| | | | | C-E | n.s. (p>0.9999) | |
| | | | | C-F | n.s. (p=0.9991) | |
| | | | | C-G | n.s. (p=0.9629) | |
| | | | | C-H | n.s. (p=0.7628) | |
| | | | | D-E | n.s. (p=0.2732) | |
| | | | | D-F | n.s. (p=0.6722) | |
| | | | | D-G | n.s. (p=0.9066) | |
| | | | | D-H | n.s. (p=0.9956) | |
| | | | | E-F | n.s. (p=0.9932) | |
| | | | | E-G | n.s. (p=0.9066) | |
| | | | | E-H | n.s. (p=0.6407) | |
| | | | | F-G | n.s. (p=0.9996) | |
| | | | | F-H | n.s. (p=0.9629) | |
| | | | | G-H | n.s. (p=0.9991) | |
| 3b | 4xTS-FF3 (A) vs. 3xTS-FF3 (B) vs. 2xTS-FF3 (C) vs. 1xTS-FF3 (D) vs. 0xTS (E) | 660.7 | *** (p<0.001) | A-B | *** (p<0.001) | 10 |
| | | | | A-C | *** (p<0.001) | |
| | | | | A-D | *** (p<0.001) | |
| | | | | A-E | *** (p<0.001) | |
| | | | | B-C | ** (p=0.0023) | |
| | | | | B-D | *** (p<0.001) | |
| | | | | B-E | *** (p<0.001) | |
| | | | | C-D | *** (p<0.001) | |
| | | | | C-E | *** (p<0.001) | |
| | | | | D-E | *** (p<0.001) | |
| 3c | 0xTS (A) vs. 1xTS-FF3 (B) vs. 2xTS-FF3 (C) vs. 3xTS-FF3 (D) vs. | 42.59 | *** (p<0.001) | A-B | n.s. (p=0.0690) | 10 |
| | | | | A-C | *** (p<0.001) | |
| | | | | A-D | *** (p<0.001) | |
| | | | | A-E | *** (p<0.001) | |
| | | | | B-C | ** (p=0.0025) | |
| | | | | B-D | * (p=0.0147) | |
| | | | | B-E | *** (p<0.001) | |
| | | | | C-D | n.s. (p=0.7519) | |

| | | | | | | |
|---|---|---|---|---|---|---|
| | 4xTS-FF3 (E) | | | C-E | * (p=0.0396) | |
| | | | | D-E | ** (p=0.0064) | |
| 4d | *Receivers* (no miRNA) (A)<br>vs.<br>*Receivers* (no miRNA) + eGFP (B)<br>vs.<br>miR-FF3 *Receivers* (C)<br>vs.<br>miR-FF3 *Receivers* + eGFP (D) | 77.00 | ***<br>(p<0.001) | A-B | n.s. (p=0.6259) | 8 |
| | | | | A-C | *** (p<0.001) | |
| | | | | A-D | *** (p<0.001) | |
| | | | | B-C | *** (p<0.001) | |
| | | | | B-D | *** (p<0.001) | |
| | | | | C-D | ** (p=0.0030) | |
| S25 | *Receivers* (no miRNA) (A)<br>vs.<br>*Receivers* (no miRNA) + eGFP (B)<br>vs.<br>miR-FF3 *Receivers* (C)<br>vs.<br>miR-FF3 *Receivers* + eGFP (D)<br>vs.<br>WT (E)<br>vs.<br>WT + eGFP (F) | 356.4 | ***<br>(p<0.001) | A-B | *** (p<0.001) | 12 |
| | | | | A-C | *** (p<0.001) | |
| | | | | A-D | n.s. (p=0.0724) | |
| | | | | A-E | *** (p<0.001) | |
| | | | | A-F | *** (p<0.001) | |
| | | | | B-C | *** (p<0.001) | |
| | | | | B-D | *** (p<0.001) | |
| | | | | B-E | *** (p<0.001) | |
| | | | | B-F | *** (p<0.001) | |
| | | | | C-D | * (p=0.0345) | |
| | | | | C-E | *** (p<0.001) | |
| | | | | C-F | *** (p<0.001) | |
| | | | | D-E | *** (p<0.001) | |
| | | | | D-F | *** (p<0.001) | |
| | | | | E-F | n.s. (p=0.2170) | |

**Supplementary Table S10: Mixed effects model analyses of tBFP expression of *Receiver* cells transfected with precursor miRNA and co-cultured with K562 *Senders*.** Overview of two-tailed linear mixed effects models used to compare tBFP expression of *Receiver* cells transfected with precursor miRNA and co-cultured with K562 *Senders*, as shown in Fig. 2f and Supplementary Fig. S16b. 'Condition' was modeled as a fixed effect, with 'well' treated as random intercept. Sample size is n=12 with 3 independently seeded and transfected wells per condition and 4 fields of view (FOV) per well. Reported values include group means, fold change, effect size (mean differences, B-A ± S.E.M.), 95% confidence interval (95% CI), t-values, two-tailed p-values, and degrees of freedom (df). n.s.: p > 0.05, *p ≤ 0.05, **p ≤ 0.01, ***p ≤ 0.001.

| Compared groups | Mean A | Mean B | Fold change | B-A ± S.E.M. | 95% CI | t-value | p-value | df |
|---|---|---|---|---|---|---|---|---|
| 0xTS + *Senders* with scrambled (A)<br>vs.<br>0xTS + *Senders* with miR-FF3 (B) | 0.5970 | 0.7464 | 0.8 | 0.1494 ± 0.1030 | -0.1365 to 0.4354 | 1.451 | n.s. (p=0.2205) | 4 |
| 4xTS-FF3 + *Senders* with scrambled (A)<br>vs.<br>4xTS-FF3 + *Senders* with miR-FF3 (B) | 1.094 | 0.1567 | 6.5 | -0.9370 ± 0.1877 | -1.458 to -0.4159 | 4.992 | **<br>(p=0.0075) | 4 |

**Supplementary Table S11: Mixed effects model analyses of feedback circuit performance in *Receiver* cells cultured on plate-bound eGFP.** Overview of two-tailed linear mixed effects models used to compare tBFP expression of *Receiver* cells cultured on plate-bound eGFP, as shown in Fig. 5f. 'Condition' was modeled as a fixed effect, with 'run' treated as random intercept. Sample size is n=9 with 3 independent runs per condition, performed on different days, and 3 wells per run. Reported values include group means, fold change, effect size (mean differences, B-A ± S.E.M.), 95% confidence interval (95% CI), t-values, two-tailed p-values, and degrees of freedom (df). n.s.: $p > 0.05$, $^{*}p \leq 0.05$, $^{**}p \leq 0.01$, $^{***}p \leq 0.001$.

| Compared groups | Mean A | Mean B | Fold change | B-A ± S.E.M. | 95% CI | t-value | p-value | df |
|---|---|---|---|---|---|---|---|---|
| 4xTS-FF3 + eGFP (t=48h) (A) vs. 4xTS-FF3 + miR-FF3 + eGFP (t=48h) (B) | 0.9786 | 1.000 | 1.0 | 0.02144 ± 0.2373 | -0.6373 to 0.6802 | 0.09035 | n.s. (p=0.9323) | 4 |
| 4xTS-FF3 + eGFP (t=96h) (A) vs. 4xTS-FF3 + miR-FF3 + eGFP (t=96h) (B) | 1.279 | 0.5215 | 2.5 | -0.7572 ± 0.1750 | -1.243 to -0.2713 | 4.327 | * (p=0.0124) | 4 |

#### Supplementary Figures

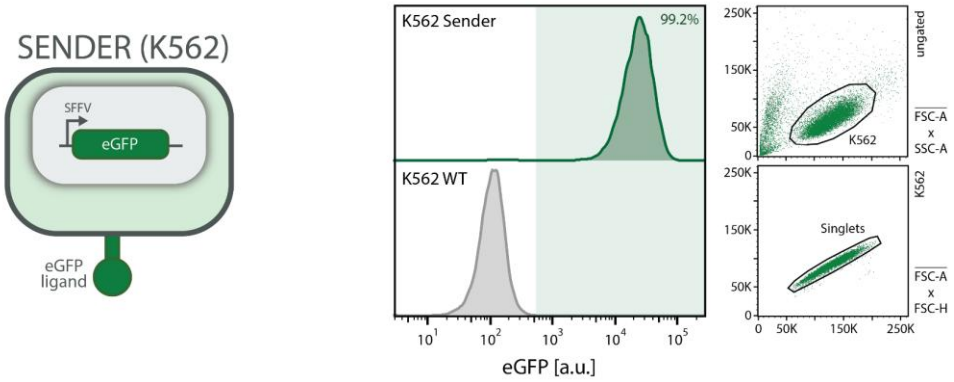


**Supplementary Figure S1: Characterization of K562 *Sender* cells expressing membrane-tethered eGFP ligand.** K562 cells were stably transduced to express membrane-tethered eGFP ligand under the control of a SFFV promoter (Methods). Lentiviral transfer plasmid, pHR_SFFV_eGFP-ligand, used to engineer K562 *Sender* cells was a gift from Wendell Lim (Supplementary Table S2; Addgene plasmid # 79129; http://n2t.net/addgene:79129; RRID:Addgene_79129)[1]. Following transduction, cells were bulk-sorted for positive eGFP expression using fluorescence-activated cell sorting (FACS; Methods). Flow cytometry validation confirms successful integration of the eGFP gene in sorted K562 *Sender* cells (green) compared to untransduced K562 wild-type (WT) control (grey). Right panels show the gating strategy (K562: FSC-A vs. SSC-A; Singlets: FSC-A vs. FSC-H).

**a**

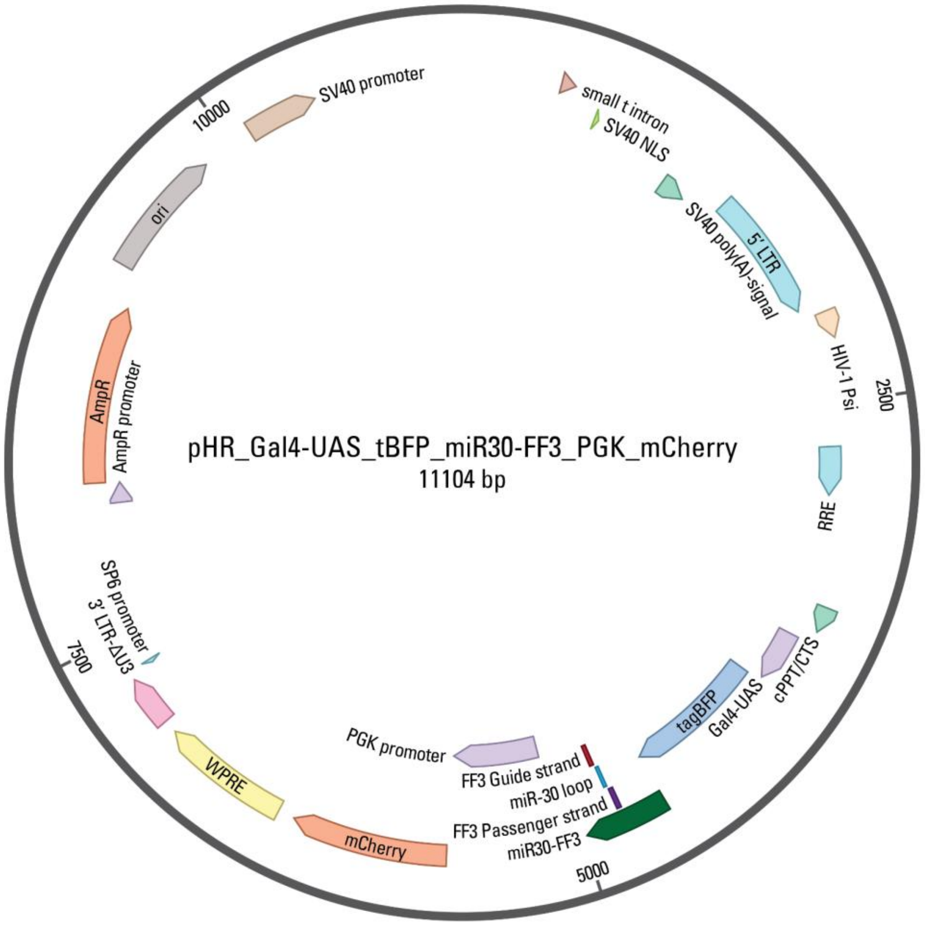


**b**

```
taggcgcgccataacttccggccgcaagccttgttaagtgctcgcttcg
gcagcacatatactatgttgaatgaggcttcagtactttacagaatcgt
tgcctgcacatcttggaaacacttgctgggattacttcttcaggttaac
ccaacagaaggctcgagaaggtatattgctgttgacagtgagcgcacga
tatgggctgaatacaaatagtgaagccacagatgtatttgtattcagcc
catatcgtttgcctactgcctcggaattcaaggggctactttaggagca
attatcttgtttactaaaactgaataccttgctatctctttgatacatt
tttacaaagctgaattaaaatggtataaattaaatcacttttt
```

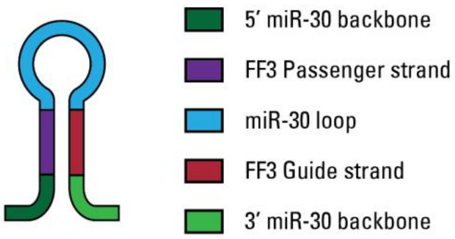


**Supplementary Figure S2: Plasmid map of the lentiviral transfer plasmid for Gal4-VP64 inducible expression of tBFP and miR30-FF3, and constitutive expression of mCherry. a** Plasmid map of pHR vector containing the tBFP, miR30-FF3 (miR-FF3) and mCherry genes (Supplementary Table S2 and S3). Both tBFP and miR-FF3 expression are under control of the Gal4-UAS promoter, which can be activated by the Gal4-VP64 transcription factor. MiR-FF3 gene is placed in the 3' UTR of the tBFP gene. Expression of mCherry is constitutive under PGK promoter control, and serves as marker for successful reporter construct integration in mammalian cells. **b** Sequence of miR30-FF3 miRNA gene used in the present study (Supplementary Table S3). The sequences and positions of the miR-30 backbone and loop, and FF3 passenger and guide strand are color-coded as shown in the legend on the right.

**a**

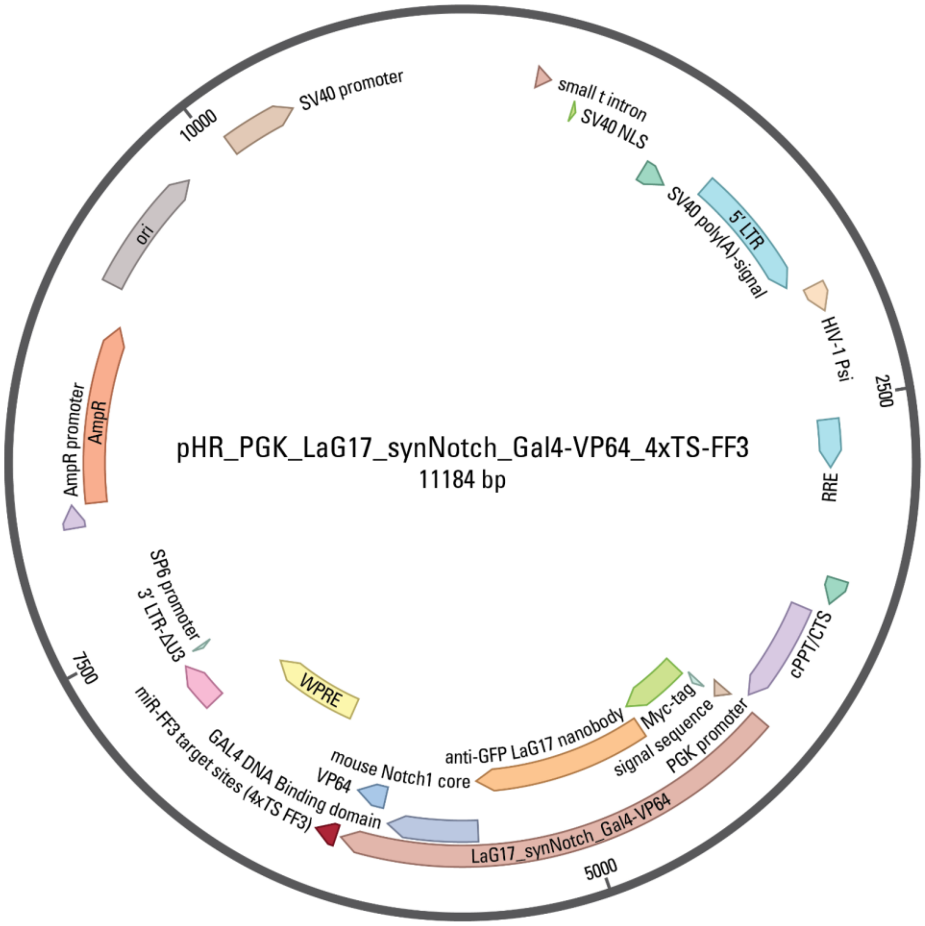


**b**

**4xTS FF3**

gatccttgacttgcgggatccaacgatatgggctgaatacaaacgacaacgatatgggctgaa
tacaaaccctaacgatatgggctgaatacaaaaccaaacgatatgggctgaatacaaacccgg

- 5' Linker
- FF3 Target site
- Spacer
- 3' Linker

**Supplementary Figure S3: Plasmid map of the lentiviral transfer plasmid for constitutive expression of the synNotch receptor with four miR-FF3 target sites. a** Plasmid map of pHR vector containing the synNotch receptor gene and miR-FF3 target sites (Supplementary Table S2 and S4). Expression of the synNotch receptor is constitutive and under control of a PGK promoter. The synNotch receptor consists of a Myc-tag, an extracellular anti-eGFP nanobody (LaG17), a native Notch transmembrane domain (mouse Notch1), and an intracellular Gal4-VP64 transcriptional effector. Four tandem repeats of the miR-FF3 target sites are placed in the 3' UTR of the synNotch gene. **b** Sequence of the four FF3 target site repeats, including linkers and spacers (Supplementary Table S5). The sequences and positions of the linkers, miR-FF3 target sites and spacers are color-coded as shown in the legend on the right.

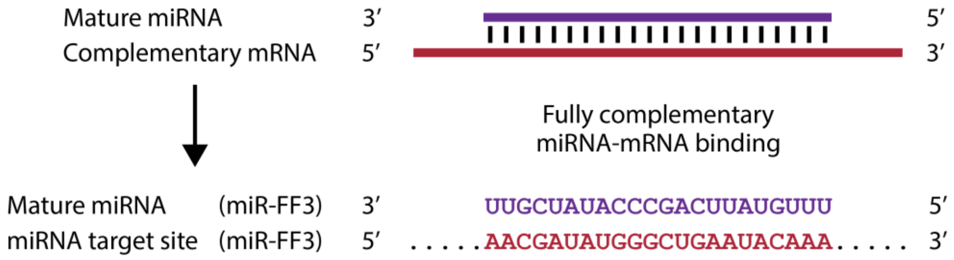


**Supplementary Figure S4: Interaction and sequence complementarity of miR-FF3 to its cognate target site.** Simple schematic showing the interaction between a miRNA and mRNA pair in case of full miRNA–target site complementarity (top). Sequence complementarity between miR-FF3 and its cognate miRNA target site (bottom; Supplementary Table S3 and S4).

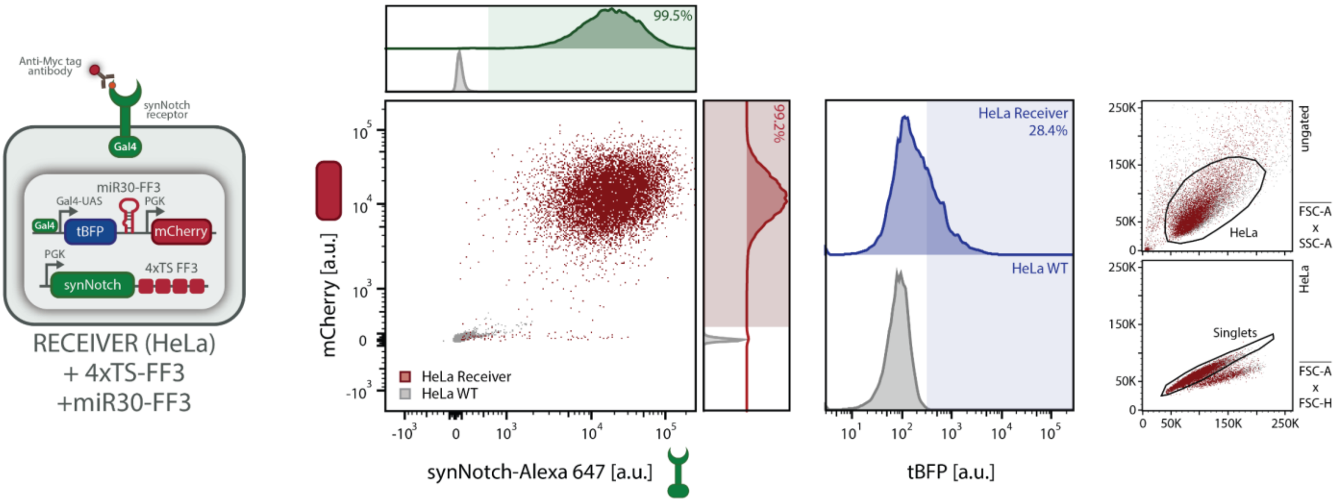


**Supplementary Figure S5: Flow cytometry characterization of HeLa *Receivers* containing a synNotch gene with four miR-FF3 target sites (4xTS) and a reporter construct harbouring the miR-FF3 gene.** Flow cytometry validation of engineered HeLa *Receiver* cells containing a synNotch gene with four miR-FF3 target sites (4xTS) and a reporter construct harbouring the miR-FF3 gene. Scatter plot and histograms of synNotch (stained with anti-Myc AF647 antibody; Methods) versus mCherry fluorescence confirm successful integration of both gene constructs (Supplementary Table S2) in HeLa *Receiver* cells (red) compared to untransduced HeLa WT control (grey), as also shown in Fig. 1d. Histogram on the right shows tBFP background expression of *Receiver* cells (blue) compared to HeLa WT control (grey). Right panels show the gating strategy (Supplementary Fig. S6a) (HeLa: FSC-A vs. SSC-A; Singlets: FSC-A vs. FSC-H).

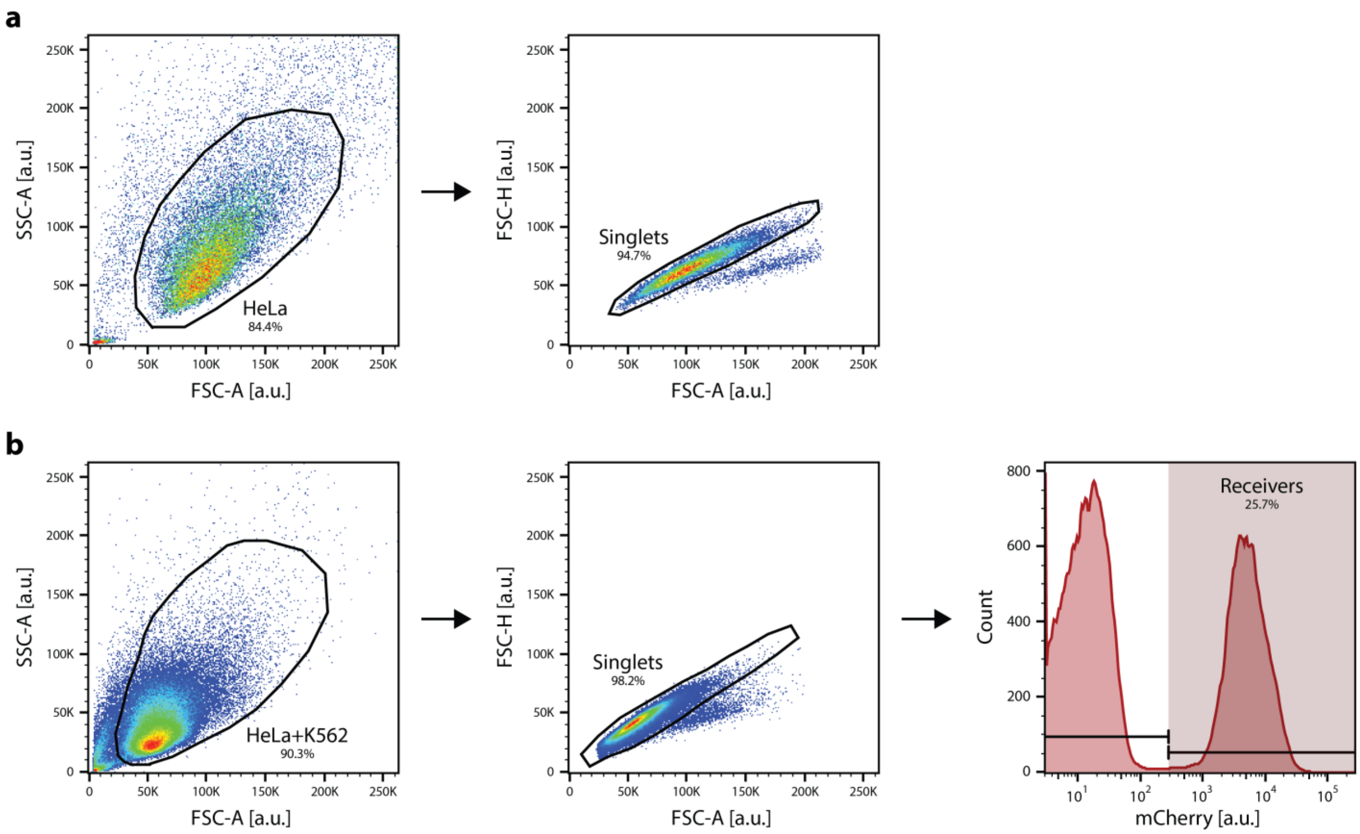


**Supplementary Figure S6: Representative gating strategy for flow cytometry analyses of *Receiver* cells. a** Representative gating strategy in pseudocolor plots used to gate for single *Receiver* cells in flow cytometry experiments (Methods). Ungated flow cytometry samples are first gated to identify HeLa cells in a FSC-A vs. SSC-A plot, whereafter HeLa cells are gated to discriminate single cells from doublets in a FSC-A vs. FSC-H plot. Subsequent analyses are performed on at least 10,000 events recorded in the 'Singlets' gate. **b** Representative gating strategy in pseudocolor plots used to gate for single *Receiver* cells in HeLa *Receiver* and K562 *Sender* co-culture experiments (Methods). Ungated flow cytometry samples are first gated to identify HeLa and K562 cells in a FSC-A vs. SSC-A plot, whereafter these events are gated to discriminate single cells from doublets in a FSC-A vs. FSC-H plot. To discriminate HeLa *Receiver* cells from K562 *Sender* cells, events are gated on positive mCherry expression. Subsequent analyses are performed on at least 10,000 events recorded in the '*Receivers*' gate.

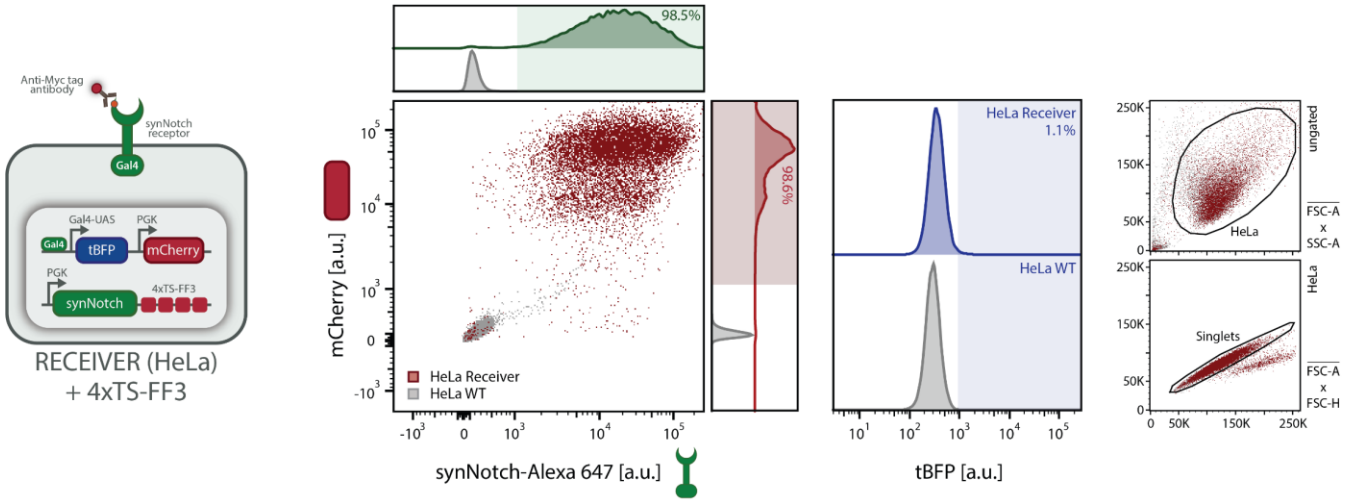


**Supplementary Figure S7: Flow cytometry characterization of HeLa *Receivers* containing a synNotch gene with four miR-FF3 target sites (4xTS).** Flow cytometry validation of engineered HeLa *Receiver* cells containing a synNotch gene with four miR-FF3 target sites (4xTS). Scatter plot and histograms of synNotch (stained with anti-Myc AF647 antibody; Methods) versus mCherry fluorescence confirm successful integration of both gene constructs (Supplementary Table S2) in HeLa *Receiver* cells (red) compared to HeLa WT control (grey). Histogram on the right shows tBFP background expression of *Receiver cells* (blue) compared to HeLa WT control (grey). Right panels show the gating strategy (Supplementary Fig. S6a) (HeLa: FSC-A vs. SSC-A; Singlets: FSC-A vs. FSC-H).

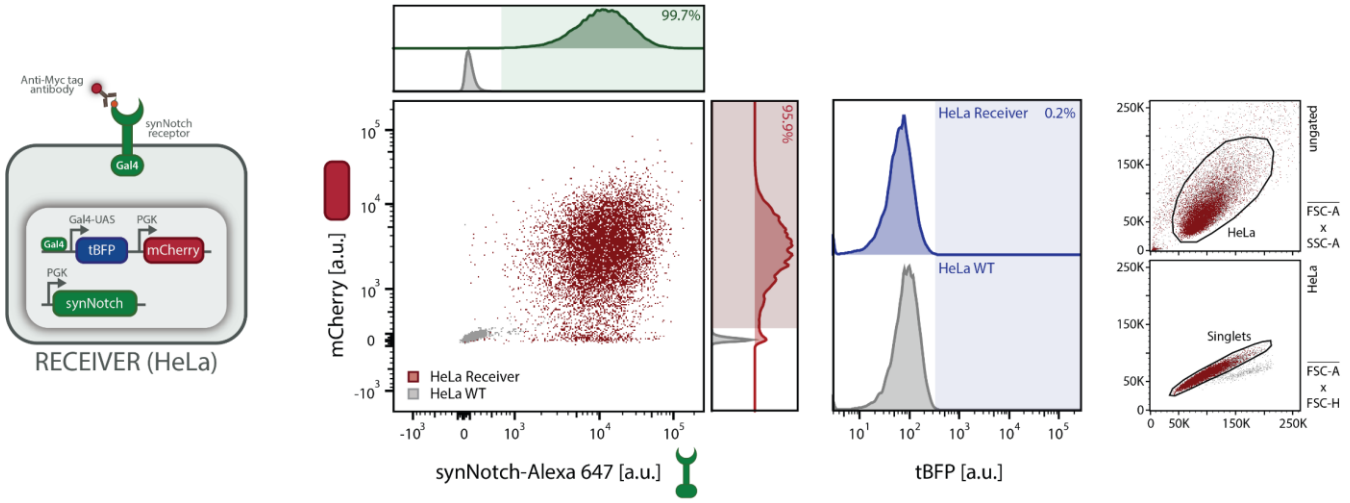


**Supplementary Figure S8: Flow cytometry characterization of the control HeLa *Receivers* containing a synNotch gene without miRNA target sites (0xTS).** Flow cytometry validation of engineered HeLa *Receiver* cells containing a synNotch gene without miRNA target sites (0xTS). Scatter plot and histograms of synNotch (stained with anti-Myc AF647 antibody; Methods) versus mCherry fluorescence confirm successful integration of both gene constructs (Supplementary Table S2) in HeLa *Receiver* cells (red) compared to HeLa WT control (grey). Histogram on the right shows tBFP background expression of *Receiver cells* (blue) compared to HeLa WT control (grey). Right panels show the gating strategy (Supplementary Fig. S6a) (HeLa: FSC-A vs. SSC-A; Singlets: FSC-A vs. FSC-H).

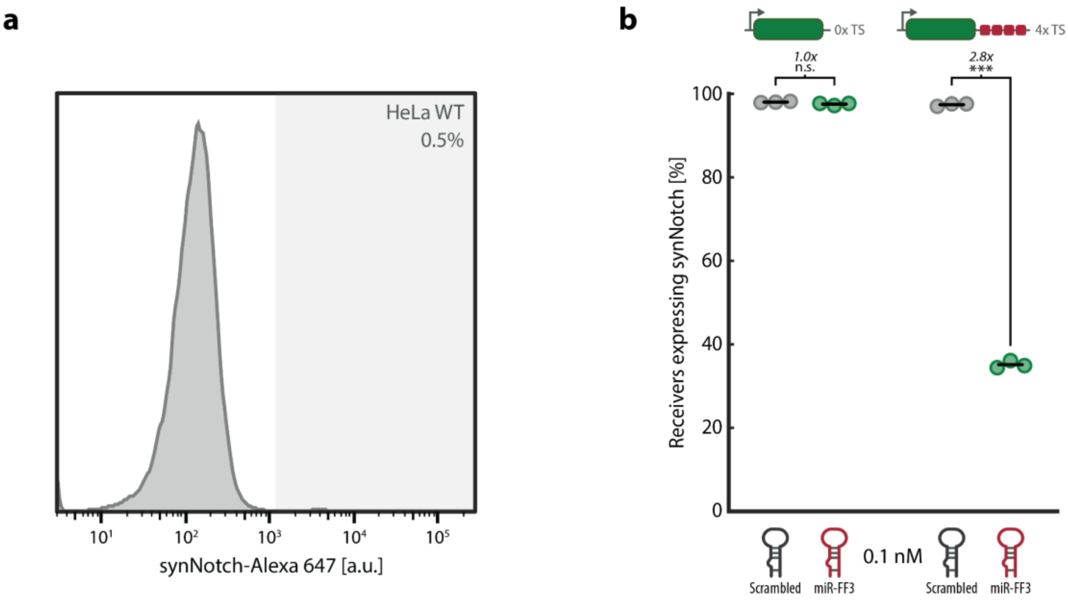


**Supplementary Figure S9: Precursor miRNA transfection of *Receiver* cells containing no miRNA target sites (0xTS) or four miR-FF3 target site repeats (4xTS) and subsequent antibody staining of synNotch receptor for flow cytometry analysis. a** Representative histogram of HeLa WT control cells used for gating of synNotch-positive cells. Shaded area indicates intensities judged as synNotch-positive cells. **b** Percentage of synNotch-positive cells (anti-Myc AF647 staining), quantified by flow cytometry analysis, in cells hosting no miRNA target sites (left) or four miR-FF3 target site repeats (right), transiently transfected with 0.1 nM scrambled (grey) or miR-FF3 (green) precursor miRNA. Horizontal lines indicate mean expression; individual data points represent independent replicates seeded, transfected, and analysed in parallel (n=3 independent wells per condition). Fold change and significance (two-tailed, unpaired t-test) are noted above data points (Supplementary Table S8). n.s.: $p > 0.05$, *$p \leq 0.05$, **$p \leq 0.01$, ***$p \leq 0.001$.

**a**

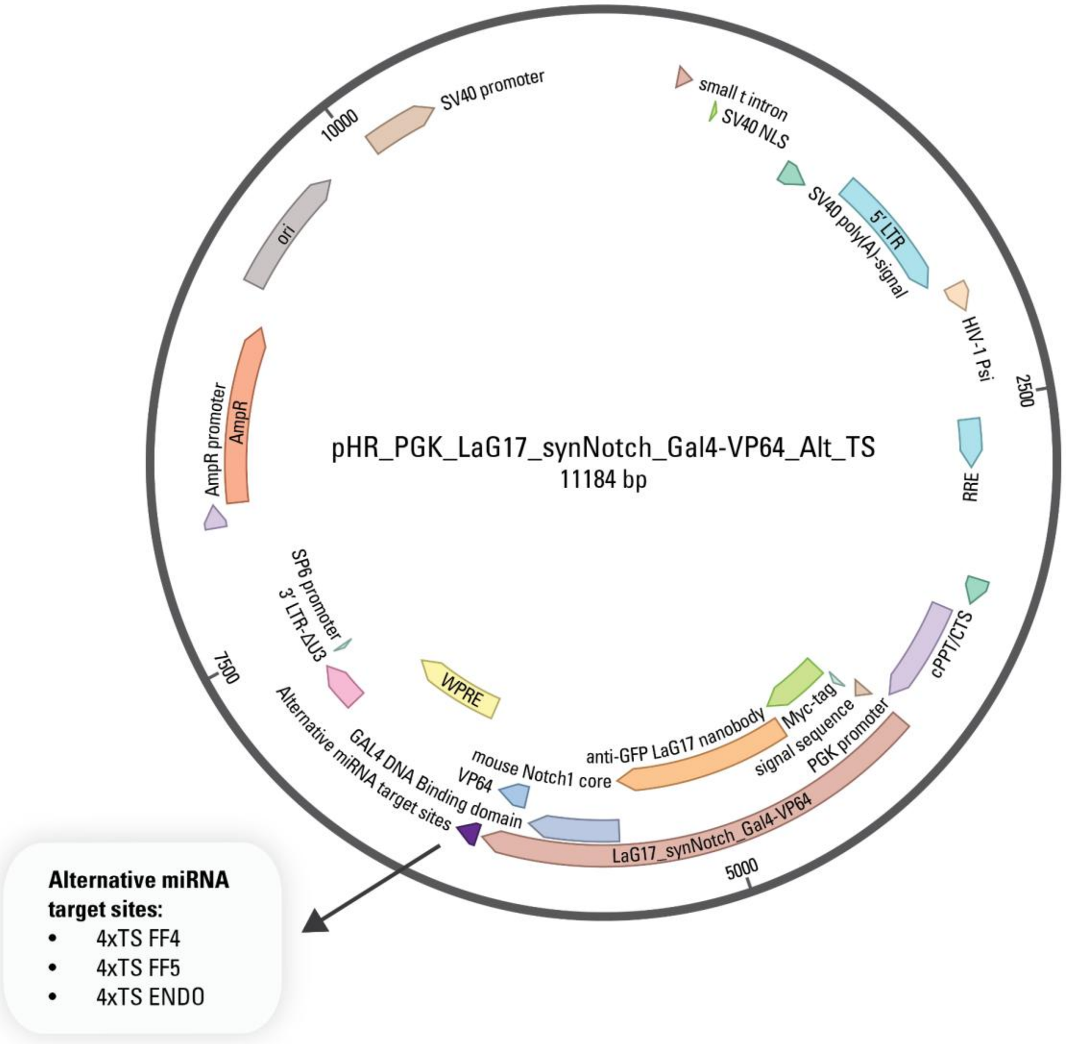


**b**

**4xTS FF4**

```
gatccttgacttgcgggatccccgcttgaagtctttaattaaacgacccgcttgaagtcttta
attaaaccctccgcttgaagtctttaattaaaaccaccgcttgaagtctttaattaaacccgg
```

**4xTS FF5**

```
gatccttgacttgcgggatccaagcactctgatttgacaattacgacaagcactctgatttga
caattaccctaagcactctgatttgacaattaaccaaagcactctgatttgacaattacccgg
```

**4xTS ENDO**

```
gatccttgacttgcgggatcccgaacgggcacgctgacaattccgaccgaacgggcacgctga
caattcccctcgaacgggcacgctgacaattcaccacgaacgggcacgctgacaattccccgg
```

- 5' Linker
- FF4 Target site
- FF5 Target site
- ENDO Target site
- Spacer
- 3' Linker

**Supplementary Figure S10: Plasmid map of the lentiviral transfer plasmid for constitutive expression of the synNotch receptor with alternative miRNA target sites. a** Plasmid map of pHR vector containing the synNotch receptor gene and alternative miRNA target sites (Supplementary Table S2). Expression of the synNotch receptor is constitutive and under control of a PGK promoter. The synNotch receptor consists of a Myc-tag, an extracellular anti-eGFP nanobody (LaG17), a native Notch transmembrane domain (mouse Notch1), and an intracellular Gal4-VP64 transcriptional effector. Four tandem repeats of either of the alternative miRNA target sites (FF4, FF5, or ENDO; Supplementary Table S4) are placed in the 3' UTR of the synNotch gene. **b** Sequences of the four miRNA target site repeats, including linkers and spacers (Supplementary Table S5). The sequences and positions of the linkers, miRNA target sites and spacers are color-coded as shown in the legend on the right.

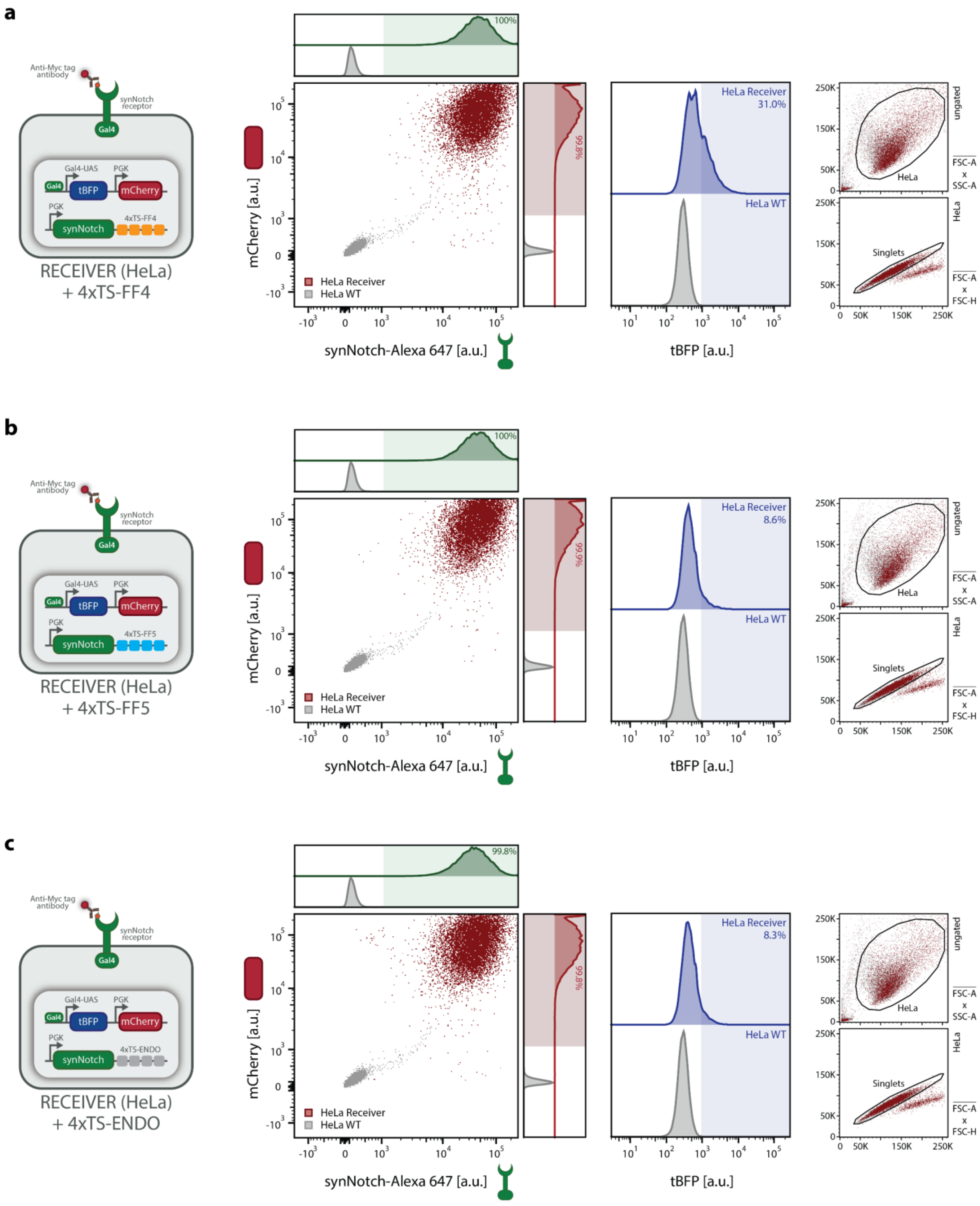


**Supplementary Figure S11: Flow cytometry characterization of HeLa *Receivers* containing a synNotch gene with four alternative miRNA target sites. a-c** Flow cytometry validation of engineered HeLa *Receiver* cells containing a synNotch gene with four alternative, non-complementary miRNA target sites

(FF4, FF5[2], or a heterologous sequence described by Endo et al. (2019)[5]; Supplementary Table S4 and S5). Scatter plot and histograms of synNotch (stained with anti-Myc AF647 antibody; Methods) versus mCherry fluorescence confirm successful integration of both gene constructs (Supplementary Table S2) in HeLa *Receiver* cells (red) compared to HeLa WT control (grey). Histogram on the right shows tBFP background expression of *Receiver cells* (blue) compared to HeLa WT control (grey). Right panels show the gating strategy (Supplementary Fig. S6a) (HeLa: FSC-A vs. SSC-A; Singlets: FSC-A vs. FSC-H).

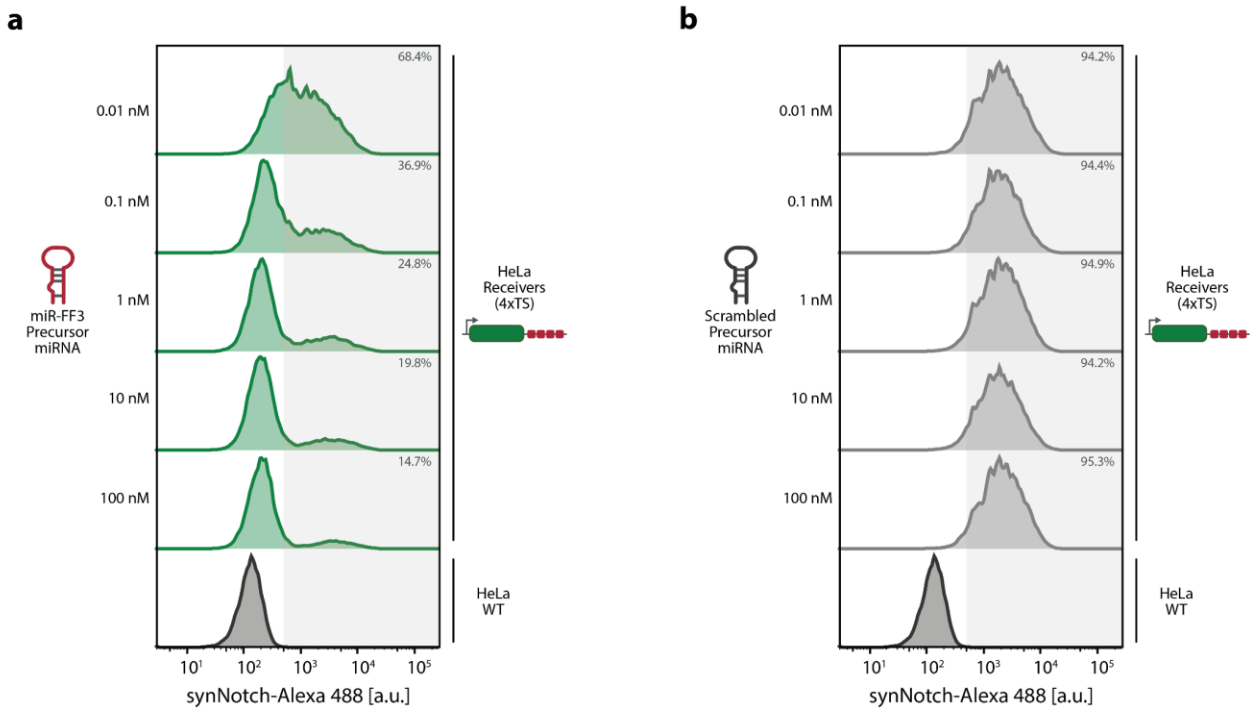


**Supplementary Figure S12: Representative flow cytometry histograms showing synNotch expression levels for different concentrations of transfected miR-FF3 and scrambled precursor in *Receiver* cells with 4xTS. a-b** Dose-response of synNotch repression. Representative flow cytometry histograms of *Receiver* cells with 4xTS, 72 h post-transfection with 0.01, 0.1, 1, 10 or 100 nM miR-FF3 (left) or scrambled (right) precursor miRNA, as also shown in Fig. 2c. Shaded area indicates synNotch-positive cells (threshold defined by HeLa WT control).

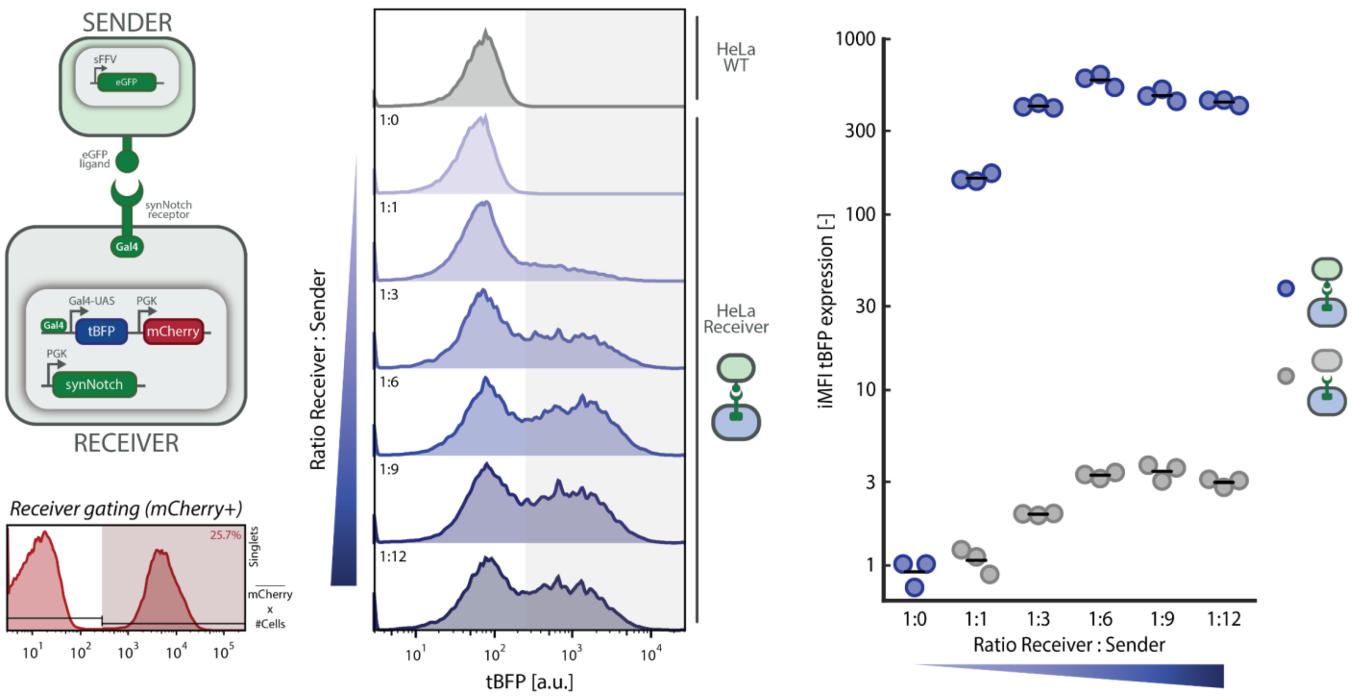


**Supplementary Figure S13: Activation of control *Receiver* cell line (0xTS) using different amounts of K562 *Sender* cells.** Activation experiment of control *Receiver* cell line (0xTS) using eGFP-expressing K562 *Sender* cells. 50,000 *Receiver* cells were seeded in a 24-well plate, before co-culturing with either eGFP-expressing K562 *Sender* cells or K562 WT control cells (Methods). *Receiver* cells are co-cultured in different ratios with K562 cells, ranging from 1:0 to 1:12 (*Receiver* : *Sender*). Samples are then analysed using flow-cytometry, where they are first gated to identify HeLa and K562 cells in a FSC-A vs. SSC-A plot, whereafter these events are gated to discriminate single cells from doublets in a FSC-A vs. FSC-H plot. To discriminate HeLa *Receiver* cells from K562 *Sender* cells, events are gated on positive mCherry expression (Supplementary Fig. S6b). Subsequent analyses are performed on at least 10,000 events recorded in the '*Receivers*' gate. Representative histograms (left) show tBFP expression of *Receivers* activated by increasing amounts of K562 *Sender* cells. Shaded area indicates tBFP-positive cells (threshold defined by HeLa WT control). Scatter plot (right) shows integrated Median Fluorescence Intensity (iMFI) of tBFP expression. Horizontal lines indicate mean expression; individual data points represent independent replicates seeded and analysed in parallel (n=3 independent wells per condition).

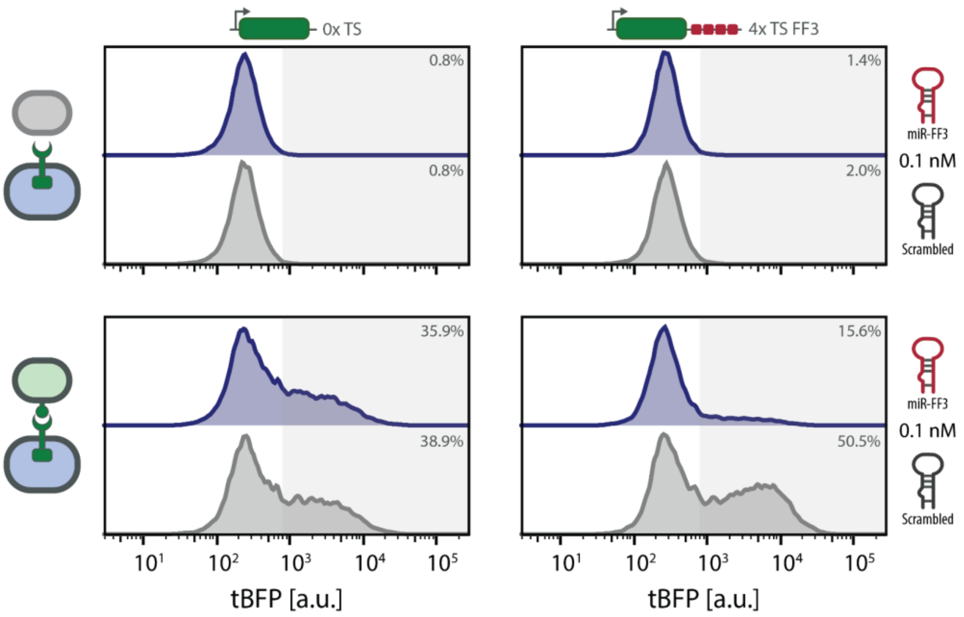


**Supplementary Figure S14: Representative flow cytometry histograms showing tBFP expression of *Receivers*, co-cultured with K562 cells, after precursor transfection.** Representative flow cytometry histograms of *Receiver* cells with 0xTS (left) or 4xTS (right) that were transiently transfected with 0.1 nM miR-FF3 or scrambled precursor miRNA, as also shown in Fig. 2d. After 24 h incubation, *Receiver* cells were co-cultured for 48 h with K562 WT cells (top) or eGFP+ *Sender* cells (bottom). Shaded area indicates tBFP-positive cells (threshold defined by HeLa WT control).

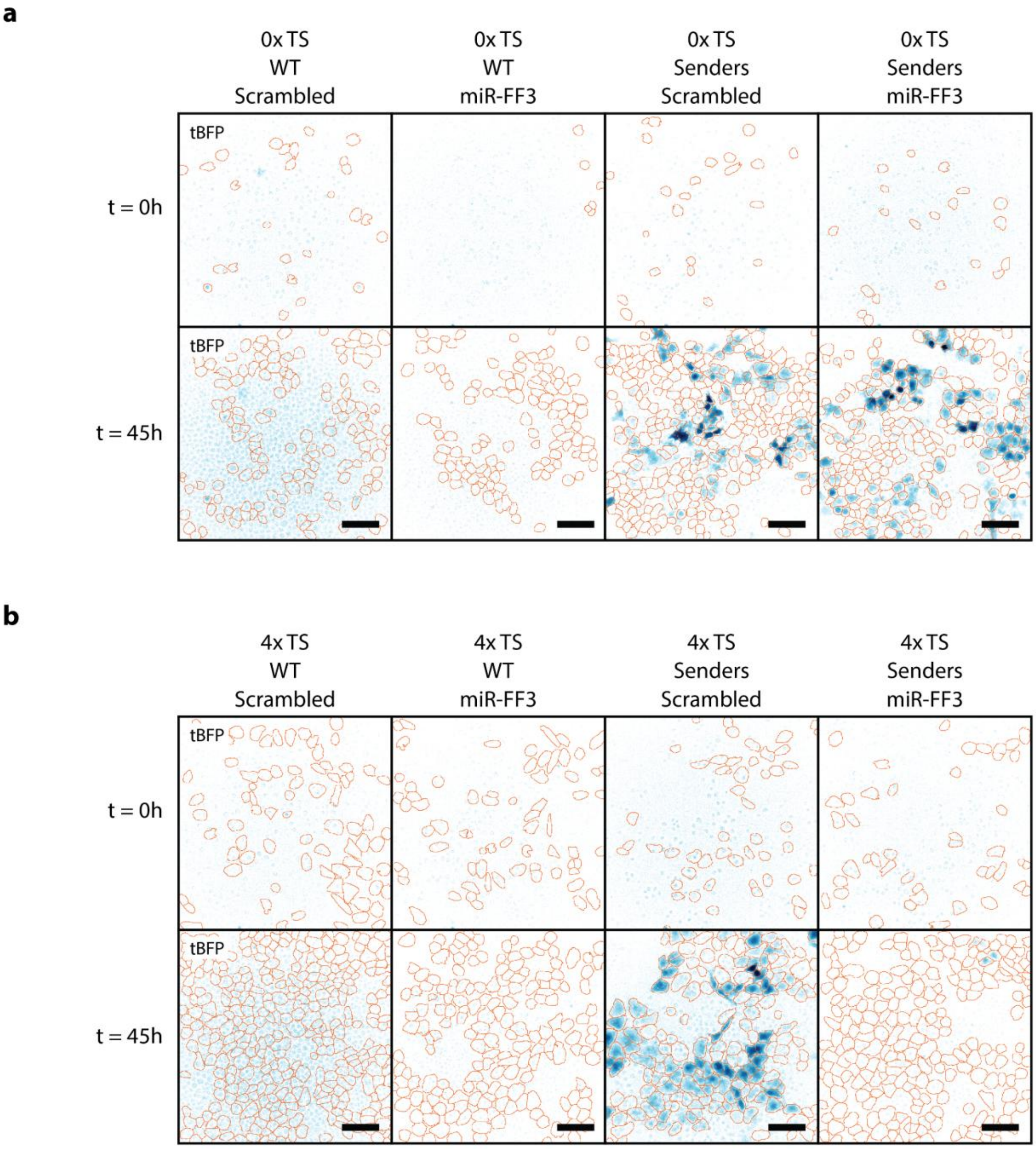


**Supplementary Figure S15: Fluorescence microscopy images showing tBFP expression of *Receivers*, co-cultured with K562 cells, after precursor transfection. a-b** Representative epifluorescence images showing tBFP expression in *Receiver* cells with 0xTS **(a)** or 4xTS **(b)**, transfected with 0.1 nM scrambled or miR-FF3 precursor miRNA and co-cultured with K562 WT or K562 *Senders*, at both 0 h and 45 h post-activation. tBFP fluorescence is displayed in blue; orange outlines indicate cellular boundaries derived from mCherry-guided image segmentation (Methods). Scale bar: 100 µm.

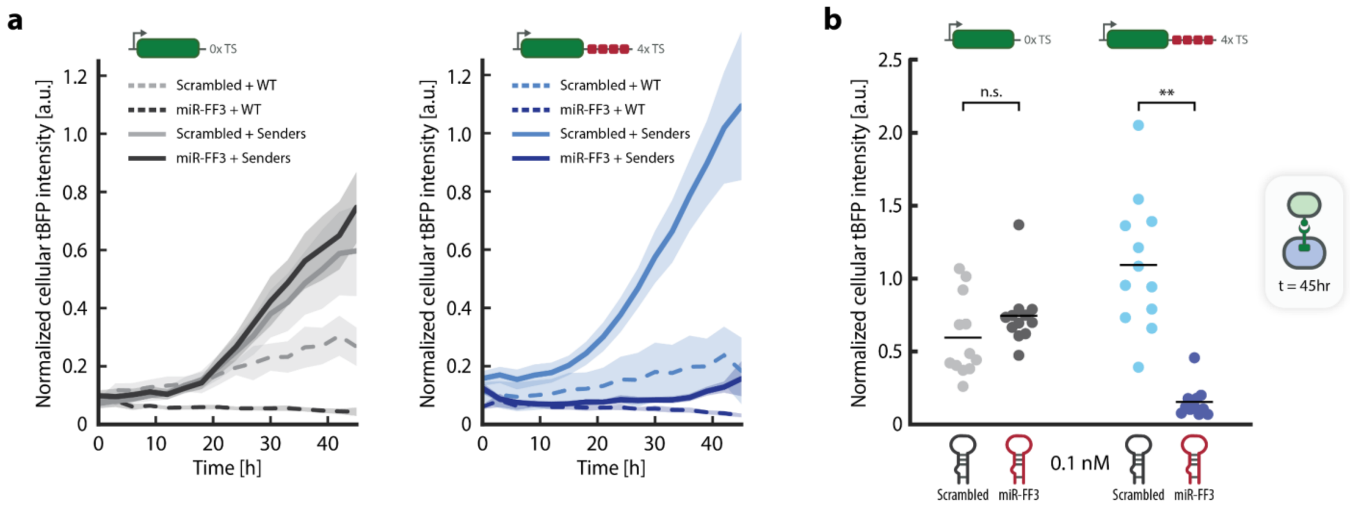


**Supplementary Figure S16: Time-lapse quantification of tBFP expression of *Receiver* cells transfected with precursor miRNA and co-cultured with K562 cells. a** Time-lapse quantification of tBFP expression (measured every 3 hours) of *Receiver* cells with 0xTS (left) or 4xTS (right), transfected with 0.1 nM precursor miRNA and co-cultured with K562 WT (dashed lines) or K562 *Senders* (solid lines) as also shown in Fig. 2f. Lines and shaded areas indicate the mean and 95% CI of normalized cellular tBFP intensity, quantified using widefield microscopy, of n=12 replicates (fields of view; FOV) from 3 independently seeded and transfected wells per condition with 4 FOVs per well. **b** Scatter plot showing normalized cellular tBFP intensity at t = 45hr post-activation of experiment performed in Fig. 2f. *Receiver* cells with 0xTS (left) or 4xTS (right) were transfected with 0.1 nM precursor miRNA and co-cultured with K562 *Senders*. Horizontal lines indicate mean normalized cellular tBFP intensity; individual data points represent n=12 replicates (FOVs) from 3 independently seeded and transfected wells per condition with 4 FOVs per well. Significance (two-tailed linear mixed effects model with 'well' treated as random intercept) are noted above data points (Supplementary Table S10) and also shown in Fig. 2f. n.s.: $p > 0.05$, *$p \leq 0.05$, **$p \leq 0.01$, ***$p \leq 0.001$.

**a**

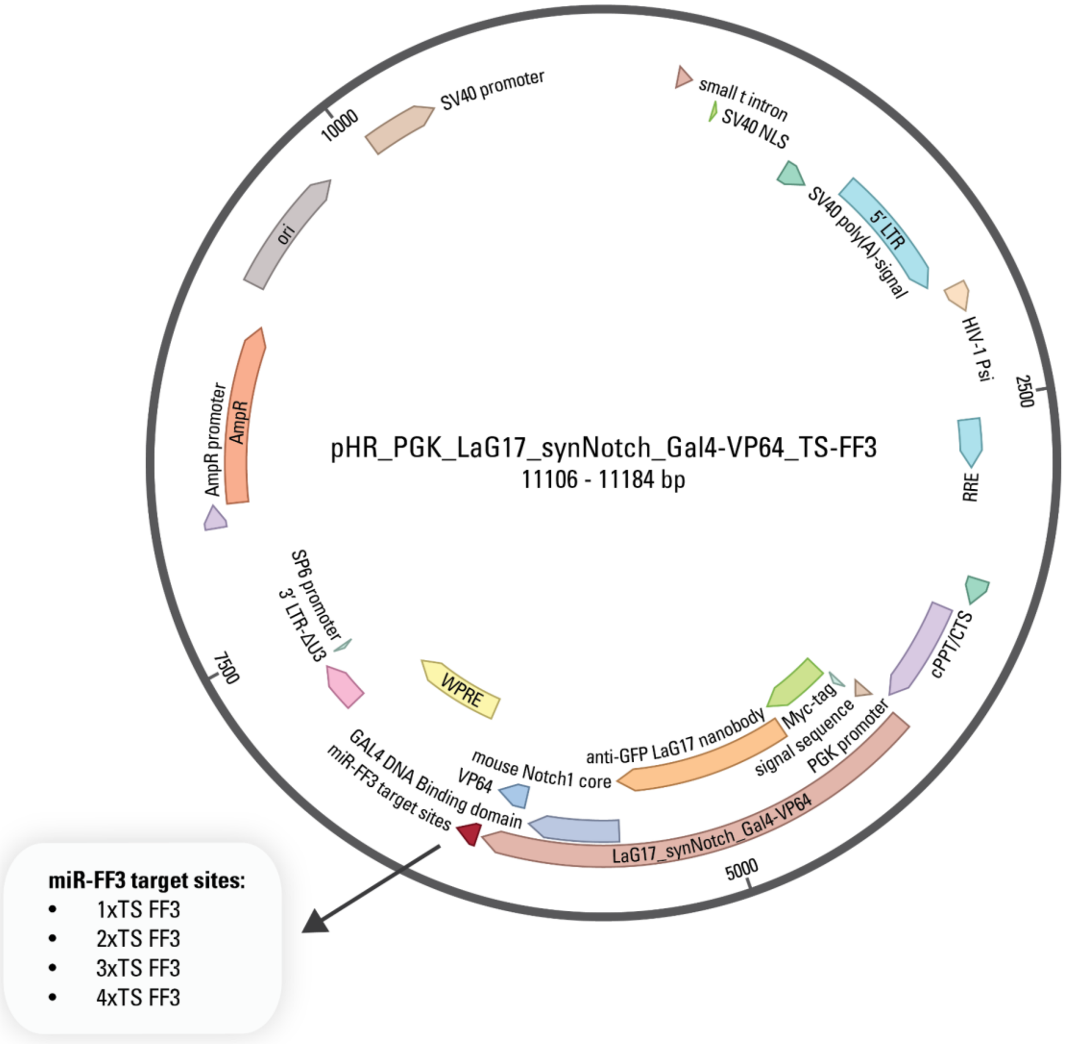


**b**

**1xTS FF3**

gatccttgacttgcgggatccaacgatatgggctgaatacaaacccgg

**2xTS FF3**

gatccttgacttgcgggatccaacgatatgggctgaatacaaacgacaacgatatgggctgaa
tacaaacccgg

**3xTS FF3**

gatccttgacttgcgggatccaacgatatgggctgaatacaaacgacaacgatatgggctgaa
tacaaaccctaacgatatgggctgaatacaaacccgg

**4xTS FF3**

gatccttgacttgcgggatccaacgatatgggctgaatacaaacgacaacgatatgggctgaa
tacaaaccctaacgatatgggctgaatacaaaaccaaacgatatgggctgaatacaaacccgg

5' Linker
FF3 Target site
Spacer
3' Linker

**Supplementary Figure S17: Plasmid map of the lentiviral transfer plasmid for constitutive expression of the synNotch receptor with one to four miR-FF3 target sites. a** Plasmid map of pHR vector containing the synNotch receptor gene and miR-FF3 target sites (Supplementary Table S2 and S4). Expression of the synNotch receptor is constitutive and under control of a PGK promoter. The synNotch receptor consists of a Myc-tag, an extracellular anti-eGFP nanobody (LaG17), a native Notch transmembrane domain (mouse Notch1), and an intracellular Gal4-VP64 transcriptional effector. One to four tandem repeats of miR-FF3 target sites are placed in the 3' UTR of the synNotch gene. **b** Sequences of the FF3 target site repeats, including linkers and spacers (Supplementary Table S5). The sequences and positions of the linkers, miR-FF3 target sites and spacers are color-coded as shown in the legend on the right.

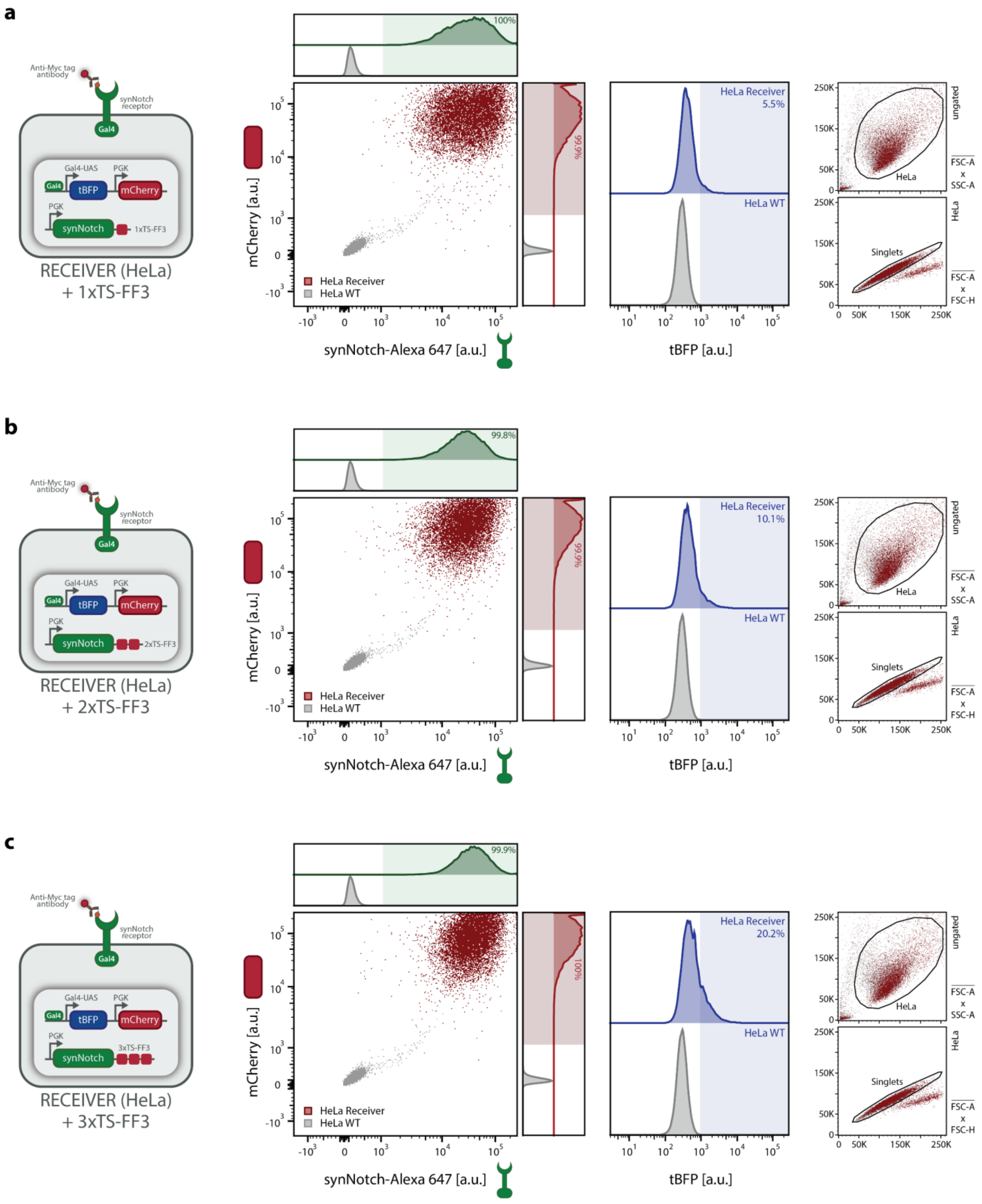


**Supplementary Figure S18: Flow cytometry characterization of HeLa *Receivers* containing a synNotch gene with one, two, or three miR-FF3 target sites (1xTS, 2xTS, 3xTS). a-c** Flow cytometry validation of engineered HeLa *Receiver* cells containing a synNotch gene with one, two, or three miR-FF3 target sites

(1xTS, 2xTS, 3xTS). Scatter plot and histograms of synNotch (stained with anti-Myc AF647 antibody; Methods) versus mCherry fluorescence confirm successful integration of both gene constructs (Supplementary Table S2) in HeLa *Receiver* cells (red) compared to HeLa WT control (grey). Histogram on the right shows tBFP background expression of *Receiver cells* (blue) compared to HeLa WT control (grey). Right panels show the gating strategy (Supplementary Fig. S6a) (HeLa: FSC-A vs. SSC-A; Singlets: FSC-A vs. FSC-H).

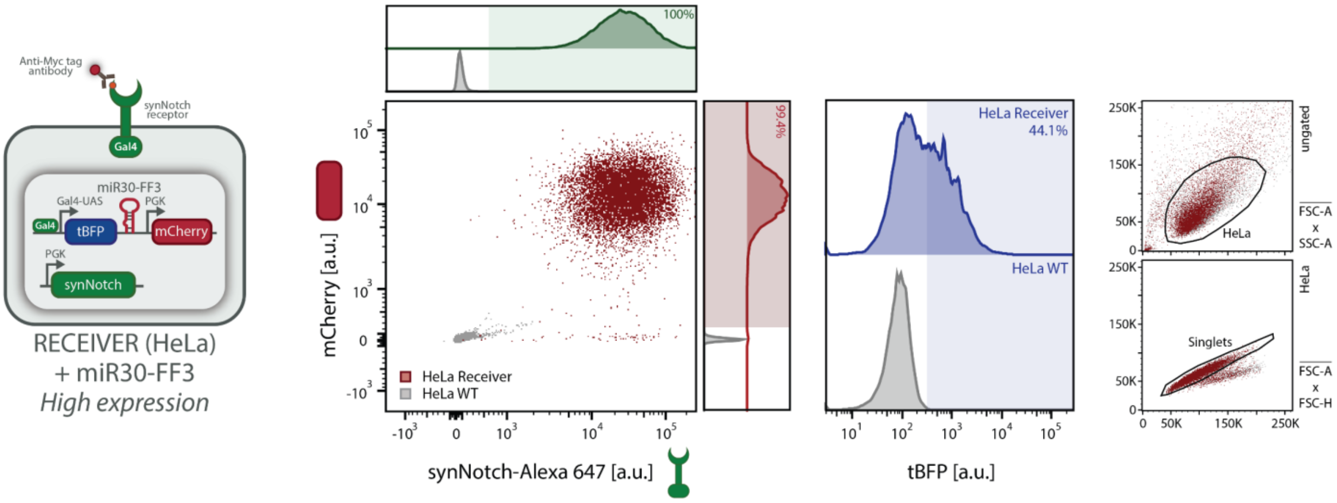


**Supplementary Figure S19: Flow cytometry characterization of HeLa *Receivers* containing a reporter construct harbouring the miR-FF3 gene.** Flow cytometry validation of engineered HeLa *Receiver* cells containing a synNotch gene without miRNA target sites and a reporter construct harbouring the miR30-FF3 gene (miR-FF3). Scatter plot and histograms of synNotch (stained with anti-Myc AF647 antibody; Methods) versus mCherry fluorescence confirm successful integration of both gene constructs (Supplementary Table S2) in HeLa *Receiver* cells (red) compared to HeLa WT control (grey). Histogram on the right shows tBFP background expression of *Receiver cells* (blue) compared to HeLa WT control (grey). Right panels show the gating strategy (Supplementary Fig. S6a) (HeLa: FSC-A vs. SSC-A; Singlets: FSC-A vs. FSC-H).

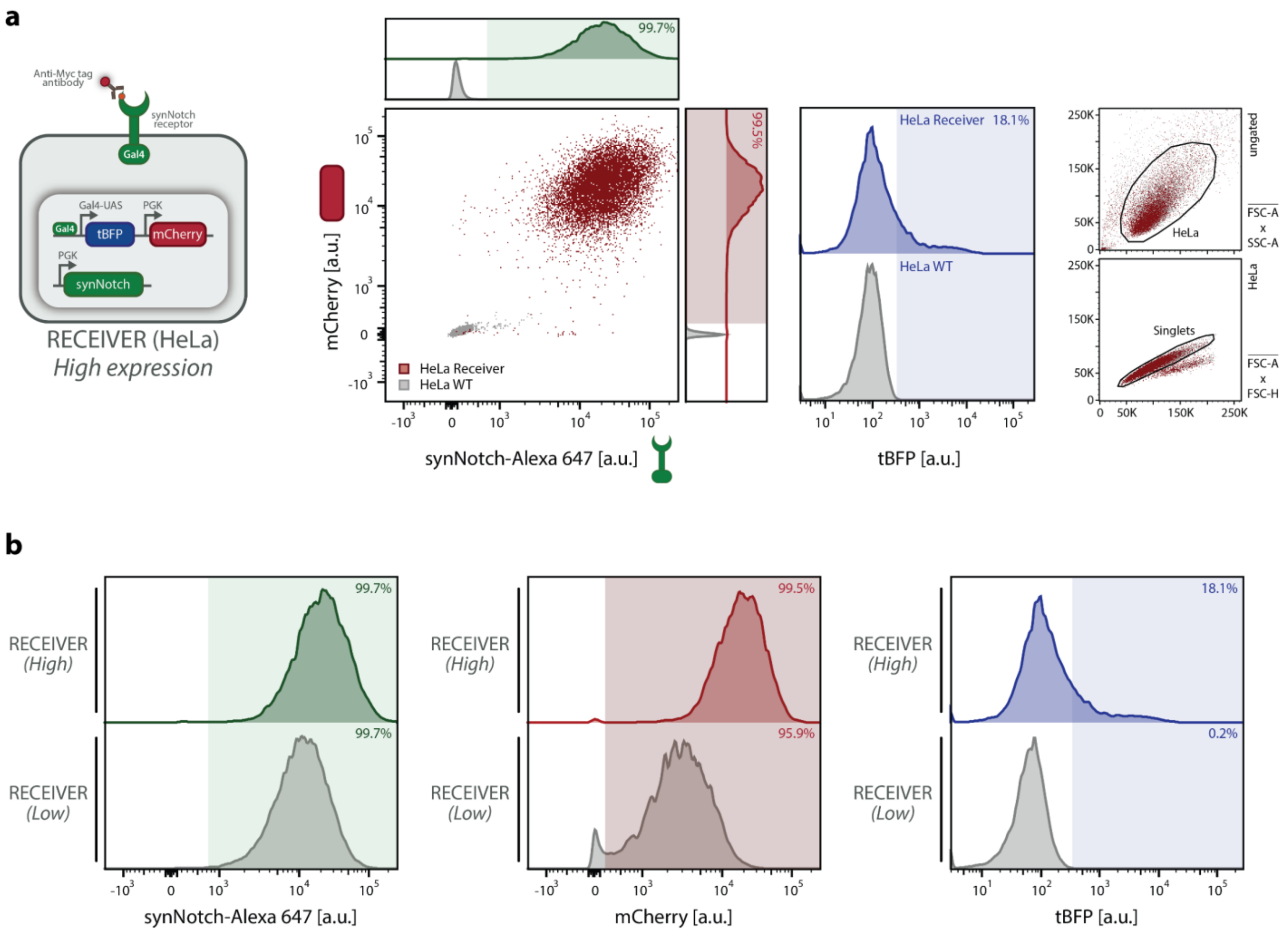


**Supplementary Figure S20: Flow cytometry characterization of HeLa *Receivers* containing a synNotch gene without miRNA target sites (0xTS) sorted for high reporter expression. a** Flow cytometry validation of engineered HeLa *Receiver* cells containing a synNotch gene without miRNA target sites (0xTS) that have been sorted for higher expression of the mCherry reporter (compared to HeLa *Receiver* shown in Supplementary Fig. S8). Scatter plot and histograms of synNotch (stained with anti-Myc AF647 antibody; Methods) versus mCherry fluorescence confirm successful integration of both gene constructs (Supplementary Table S2) in HeLa *Receiver* cells (red) compared to HeLa WT control (grey). Histogram on the right shows tBFP background expression of *Receiver cells* (blue) compared to HeLa WT control (grey). Right panels show the gating strategy (Supplementary Fig. S6a) (HeLa: FSC-A vs. SSC-A; Singlets: FSC-A vs. FSC-H). **b** Comparison of synNotch, mCherry and tBFP expression levels between 0xTS *Receivers* sorted for high (top) and low (bottom; Supplementary Fig. S8) expression of mCherry reporter. Cells that have been sorted for higher expression of mCherry, also show higher background expression of the tBFP reporter.

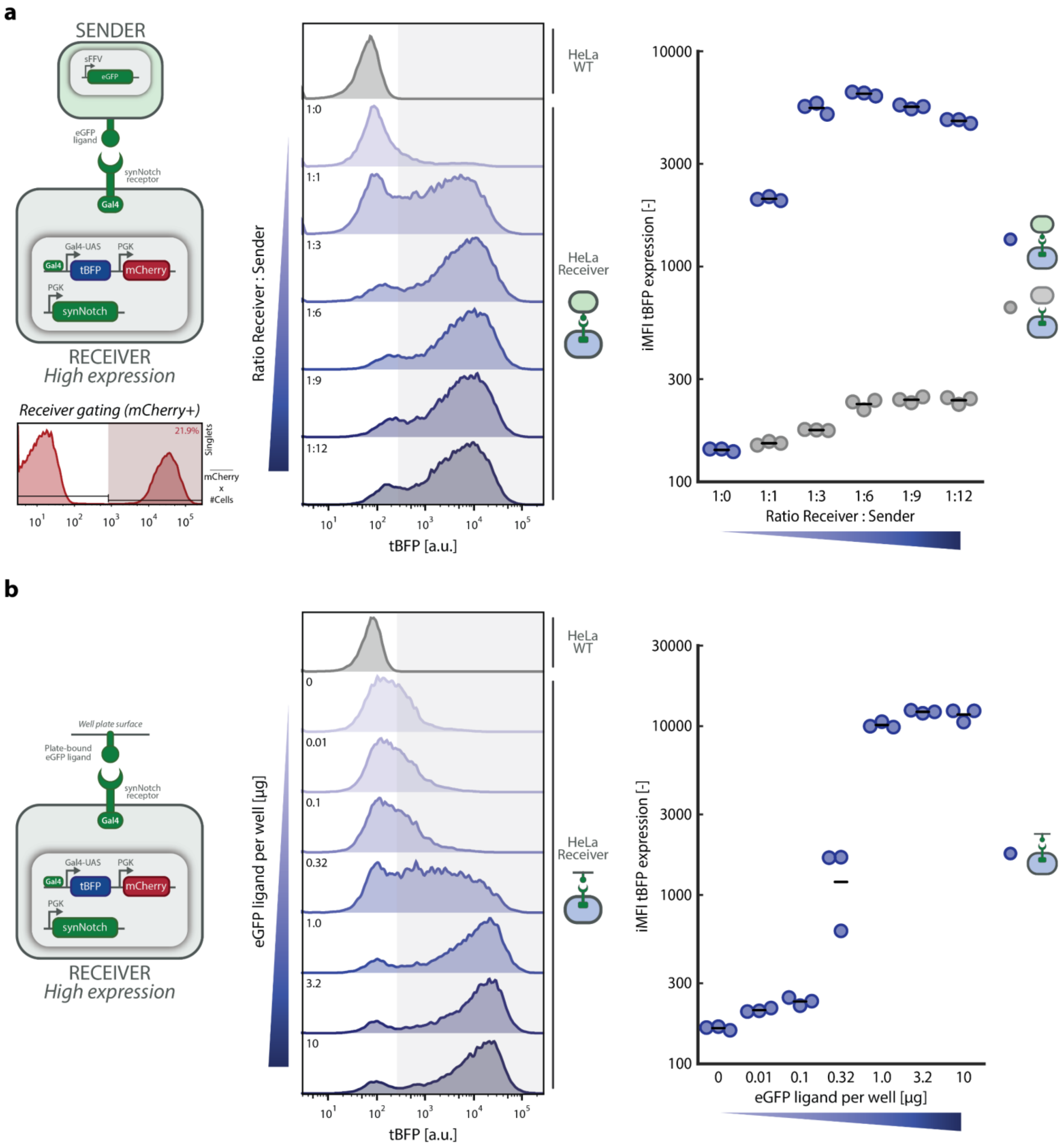


**Supplementary Figure S21: Activation of control *Receiver* cell line (0xTS), sorted for high reporter expression, using different amounts of K562 *Sender* cells or different concentrations of plate-bound eGFP ligand. a.** Activation experiment of control *Receiver* cell line (0xTS), sorted for high reporter expression, using eGFP-expressing K562 *Sender* cells. 50,000 *Receiver* cells were seeded in a 24-well plate, before co-culturing with either eGFP-expressing K562 *Sender* cells or control K562 WT cells (Methods). *Receiver* cells are co-cultured in different ratios with K562 cells, ranging from 1:0 to 1:12 (*Receiver* : *Sender*). Samples are then analysed using flow-cytometry, where they are first gated to identify HeLa and K562 cells in a FSC-A vs. SSC-A plot, whereafter these events are gated to discriminate single cells from doublets in a FSC-A vs. FSC-H plot. To discriminate HeLa *Receiver* cells from K562

*Sender* cells, events are gated on positive mCherry expression (Supplementary Fig. S6b). Subsequent analyses are performed on at least 10,000 events recorded in the '*Receivers*' gate. Representative histograms (left) show tBFP expression of *Receivers* activated by increasing amounts of K562 *Sender* cells. Shaded area indicates tBFP-positive cells (threshold defined by HeLa WT control). Scatter plot (right) shows integrated Median Fluorescence Intensity (iMFI) of tBFP expression. Horizontal lines indicate mean expression; individual data points represent independent replicates seeded and analysed in parallel (n=3 independent wells per condition). **b.** Activation experiment of control *Receiver* cell line (0xTS), sorted for high reporter expression, using plate-bound eGFP ligand. Wells of a 24-well plate were coated with different amounts of eGFP ligand one day prior to the seeding of 50,000 *Receiver* cells per well (Methods). *Receiver* cells are cultured on different amounts of plate-bound eGFP ligand, ranging from 0 to 10 µg per well. Samples are then analysed using flow-cytometry, where they are first gated to identify HeLa cells in a FSC-A vs. SSC-A plot, whereafter these events are gated to discriminate single cells from doublets in a FSC-A vs. FSC-H plot (Supplementary Fig. S6a). Subsequent analyses are performed on at least 10,000 events recorded in the 'Singlets' gate. Representative histograms (left) show tBFP expression of *Receivers* activated by increasing amounts of plate-bound eGFP ligand. Shaded area indicates tBFP-positive cells (threshold defined by HeLa WT control). Scatter plot (right) shows iMFI of tBFP expression. Horizontal lines indicate mean expression; individual data points represent independent replicates seeded and analysed in parallel (n=3 independent wells per condition).

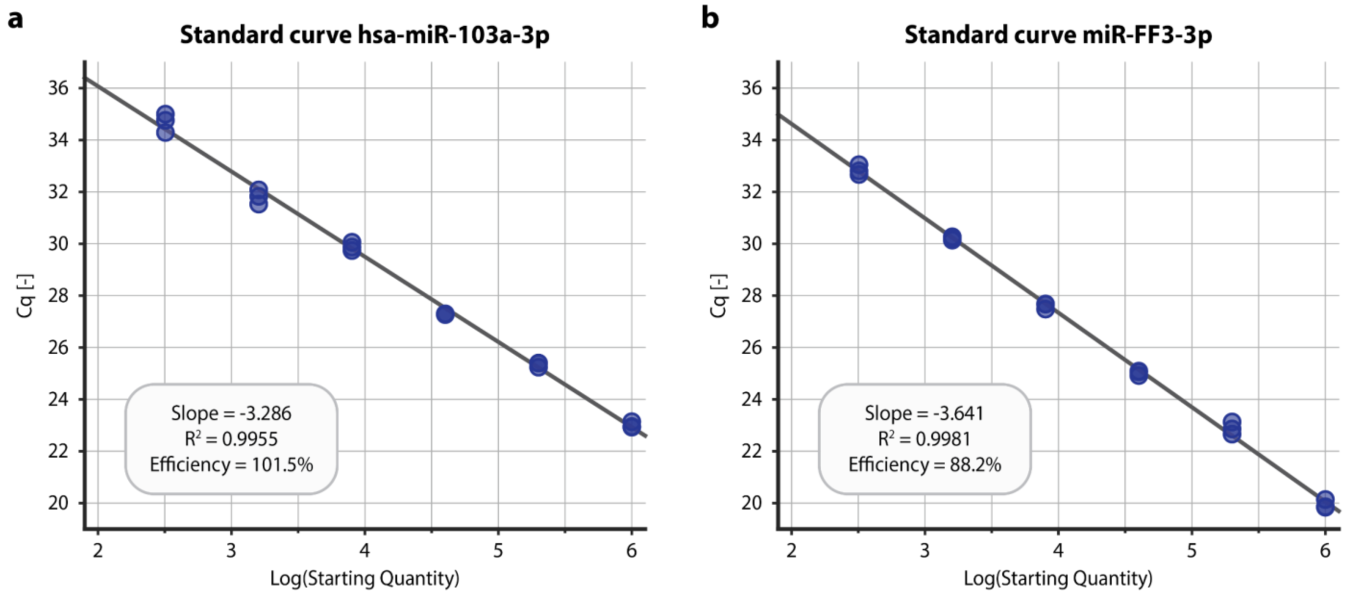


**Supplementary Figure S22: Standard curve for hsa-miR-103a-3p and miR-FF3-3p qPCR primer sets. a-b** Standard curves generated to determine the amplification efficiency of the qPCR primer sets used in this study (Supplementary Table S6). Primer specificity was confirmed by melt curve analysis showing single peaks for all reactions. **a** Standard curve for hsa-miR-103a-3p LNA primer pair. A 5-fold serial dilution of cDNA was amplified in technical triplicates, and measured Cq values were plotted against the $\log_{10}$ of the starting quantity (SQ) (Methods). Linear regression yielded a slope of −3.286, corresponding to an amplification efficiency of 101.5% ($R^2$ = 0.9955). **b** Standard curve for miR-FF3-3p LNA primer pair, generated using the same dilution series and analysis procedure. Linear regression yielded a slope of −3.641, corresponding to an amplification efficiency of 88.2% ($R^2$ = 0.9981).

**a**

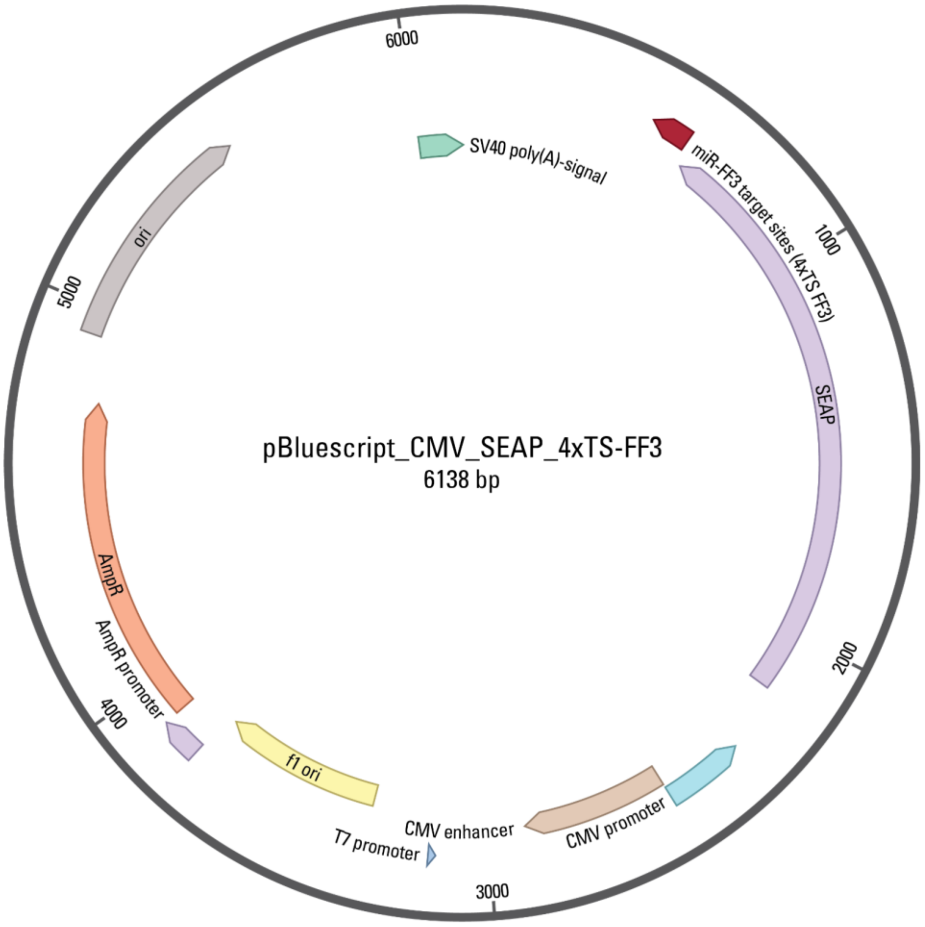


**b**

**4xTS FF3**

cccgtggtccccgcgttatccaacgatatgggctgaatacaaacgacaacgatatgggctgaa
tacaaaccctaacgatatgggctgaatacaaaaccaaacgatatgggctgaatacaaa

5′ Linker

FF3 Target site

Spacer

**Supplementary Figure S23: Plasmid map of the pBluescript expression vector for transient mammalian expression of secreted embryonic alkaline phosphatase (SEAP) with four miR-FF3 target sites. a** Plasmid map of pBluescript vector containing the SEAP reporter gene and miR-FF3 target sites (Supplementary Table S2 and S4). Expression of SEAP is constitutive and under control of a CMV promoter. Four tandem repeats of the miR-FF3 target sites are placed in the 3′ UTR of the SEAP gene. **b** Sequence of the four FF3 target site repeats, including linkers and spacers. The sequences and positions of the linker, miR-FF3 target sites and spacers are color-coded as shown in the legend on the right.

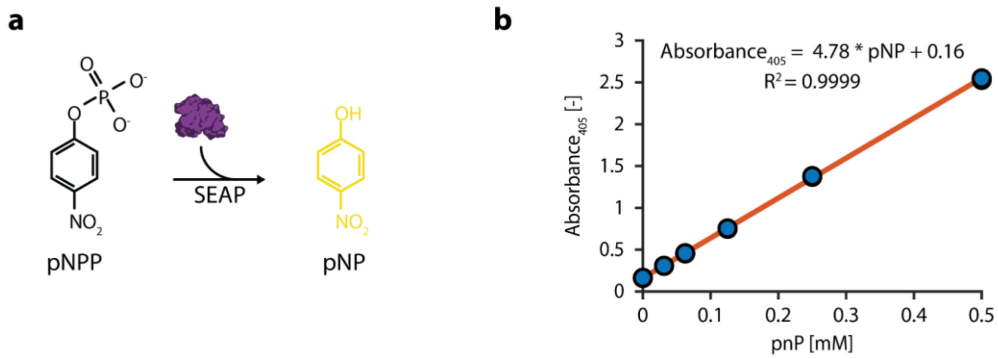


**Supplementary Figure S24: Calibration curve to determine pNPP conversion by SEAP. a** SEAP catalyses the hydrolysis of chromogenic substrate p-Nitrophenylphosphate (pNPP) to p-Nitrophenol (pNP), the absorbance of which can be measured at 405 nm. **b** To convert absorption units to amount of pNP, we titrated known concentrations of pNP (0 mM, 0.03125 mM, 0.125 mM, 0.25 mM and 0.5 mM), measured absorbance units (405 nm at 25°C), and fitted a linear regression curve through the datapoints. This yielded the following equation relating absorbance at 405 nm to the concentration of pNP (in mM) in the sample: $Absorbance_{405} = 4.78*pNP + 0.16$ ($R^2 = 0.9999$). Subsequently, the concentration of pNP is used to calculate the SEAP activity in U/L (Methods). Individual data points represent independent technical replicates (n=3).

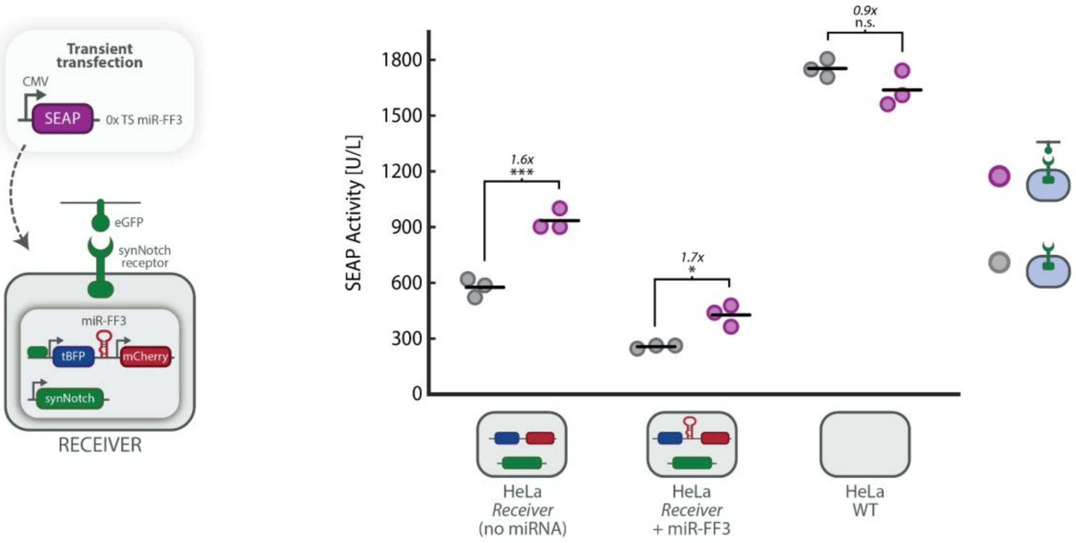


**Supplementary Figure S25: SEAP expression of SEAP construct without miRNA target sites transfected in HeLa *Receiver* and WT cells.** HeLa *Receiver* cells lacking or containing the inducible miR-FF3 cassette, or WT HeLa cells, were transiently transfected with a plasmid containing a SEAP reporter gene lacking miR-FF3 target sites (SEAP-0xTS; Supplementary Table S2), and cultured on uncoated (grey) or eGFP-coated plates (green). SEAP activity [U/L] is measured at 48 h post-transfection using a SEAP assay (Methods). A modest increase in SEAP activity was observed in *Receiver* cells cultured on eGFP-coated plates, suggesting a *Receiver*-specific synNotch-dependent effect independent of miRNA expression or target site regulation. Unlike the SEAP-4xTS reporter, which is repressed in activated miR-FF3 *Receivers* (Fig. 4d), the untargeted SEAP-0xTS did not show repression, supporting the conclusion that SEAP repression is mediated by induced functional miR-FF3 acting through complementary miRNA target sites in the reporter transcript. Horizontal lines indicate mean activity; individual data points represent independent replicates seeded, transfected, and analysed in parallel (n=3 independent wells per condition). Fold change and significance (one-way ANOVA with Tukey's multiple comparisons test) are noted above data points (Supplementary Table S9). n.s.: $p > 0.05$, *$p \leq 0.05$, **$p \leq 0.01$, ***$p \leq 0.001$.

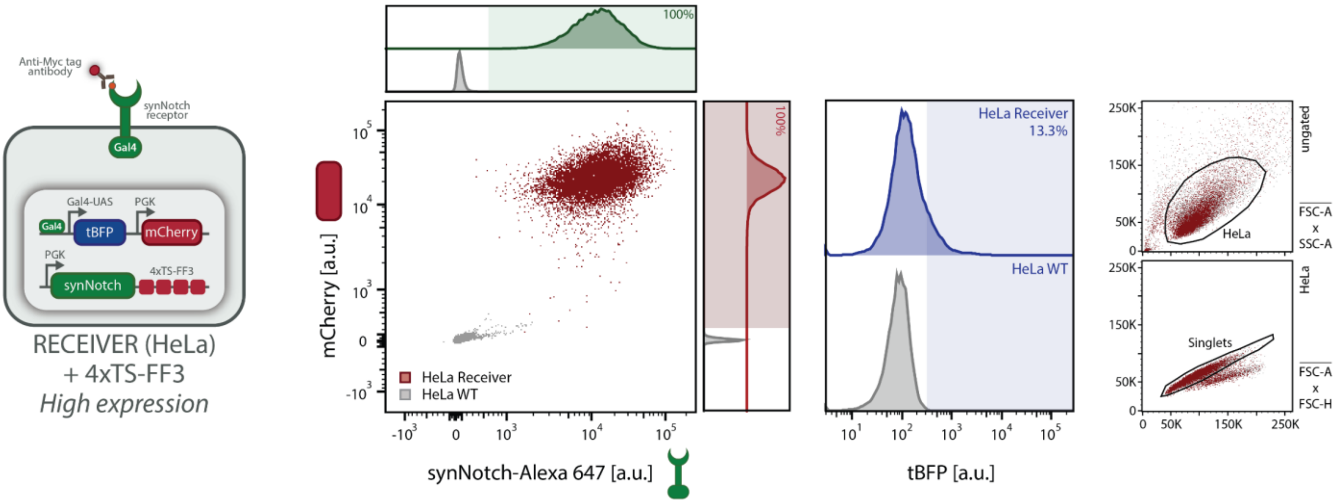


**Supplementary Figure S26: Flow cytometry characterization of HeLa *Receivers* containing a synNotch gene with four miR-FF3 target sites (4xTS) sorted for high reporter expression.** Flow cytometry validation of engineered HeLa *Receiver* cells containing a synNotch gene with four miR-FF3 target sites (4xTS) that have been sorted for higher expression of the mCherry reporter (compared to HeLa *Receiver* shown in Supplementary Fig. S7). Scatter plot and histograms of synNotch (stained with anti-Myc AF647 antibody; Methods) versus mCherry fluorescence confirm successful integration of both gene constructs (Supplementary Table S2) in HeLa *Receiver* cells (red) compared to HeLa WT control (grey). Histogram on the right shows tBFP background expression of *Receiver cells* (blue) compared to HeLa WT control (grey). Right panels show the gating strategy (Supplementary Fig. S6a) (HeLa: FSC-A vs. SSC-A; Singlets: FSC-A vs. FSC-H).

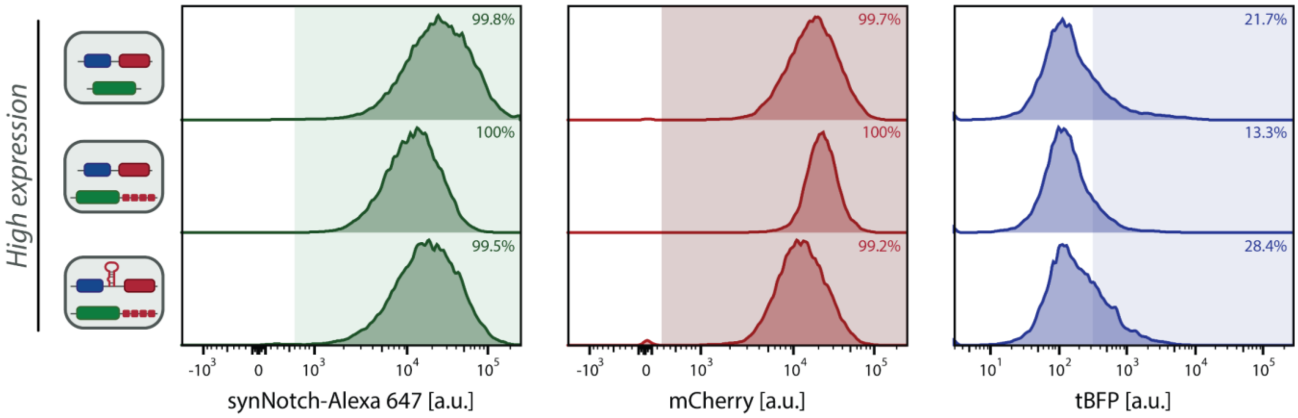


**Supplementary Figure S27: Flow cytometry characterization of HeLa *Receivers* sorted for high reporter expression.** Comparison of synNotch, mCherry and tBFP expression levels between *Receivers*, sorted for high expression of mCherry reporter, used in Fig. 5. Cells that have been sorted for higher expression of mCherry, also show higher background expression of the tBFP reporter. Top: HeLa *Receiver* (Supplementary Fig. S8); Middle: HeLa *Receiver* + 4xTS-FF3 (Supplementary Fig. S26); Bottom: HeLa *Receiver* + 4xTS-FF3 + miR30-FF3 (Supplementary Fig. S5);

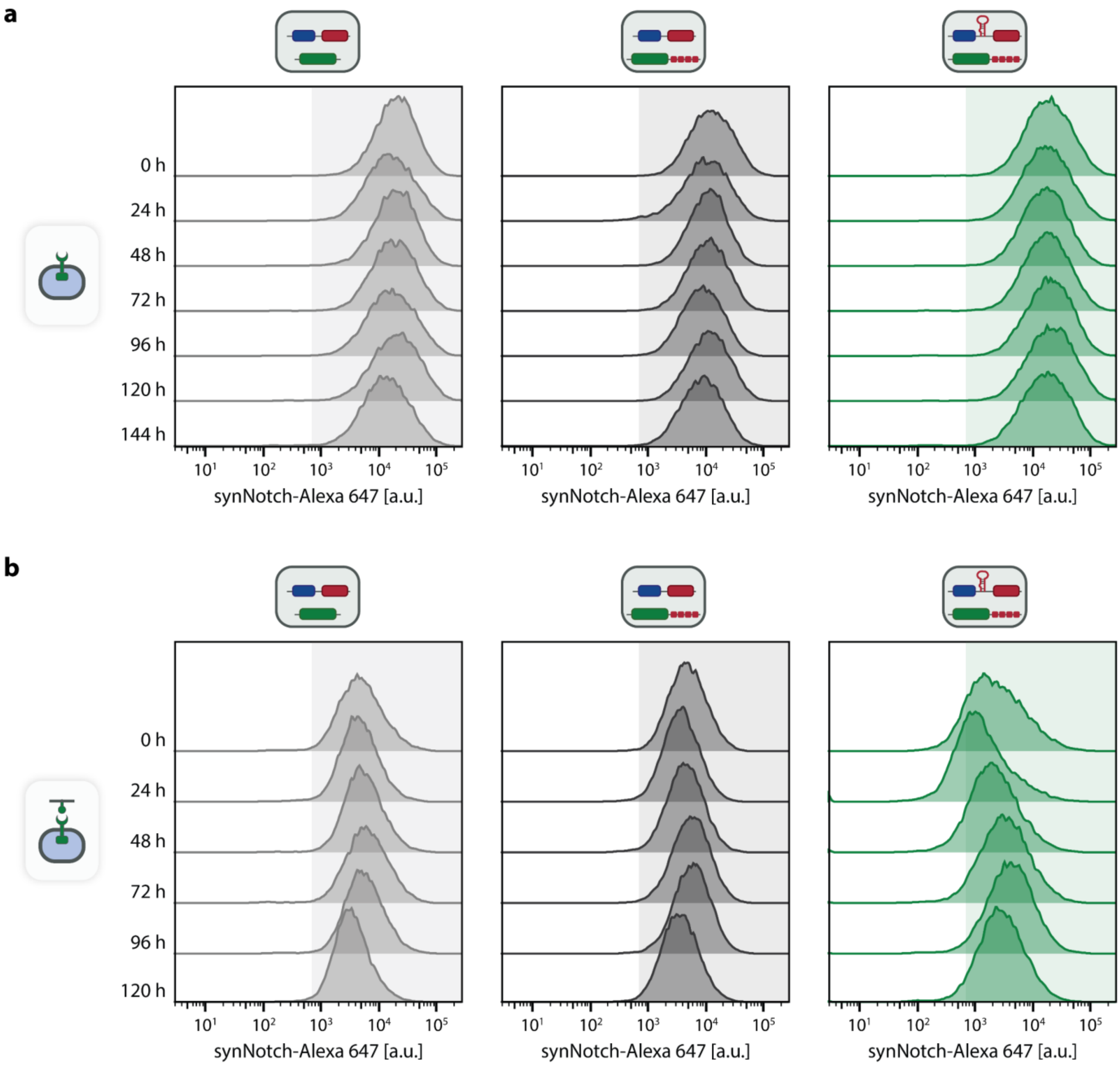


**Supplementary Figure S28: Expression of synNotch in three different *Receiver* cell lines cultured on uncoated and eGFP-coated plates. a-b** Representative flow cytometry histograms of synNotch expression as shown in Extended Data Fig. 3a-3b. Three types of *Receiver* cell lines (left: without feedback loop components, middle: without miR-FF3 gene and with miR-FF3 target sites, right: with miR-FF3 gene and target sites) were seeded at t = 0 h on uncoated **(a)** or eGFP-coated plates **(b)**. Expression of synNotch (stained with anti-Myc AF647 antibody; Methods) was quantified over time by flow cytometry analysis, with measurements every 24 hours. Shaded area indicates synNotch-positive cells (threshold defined by HeLa WT control).

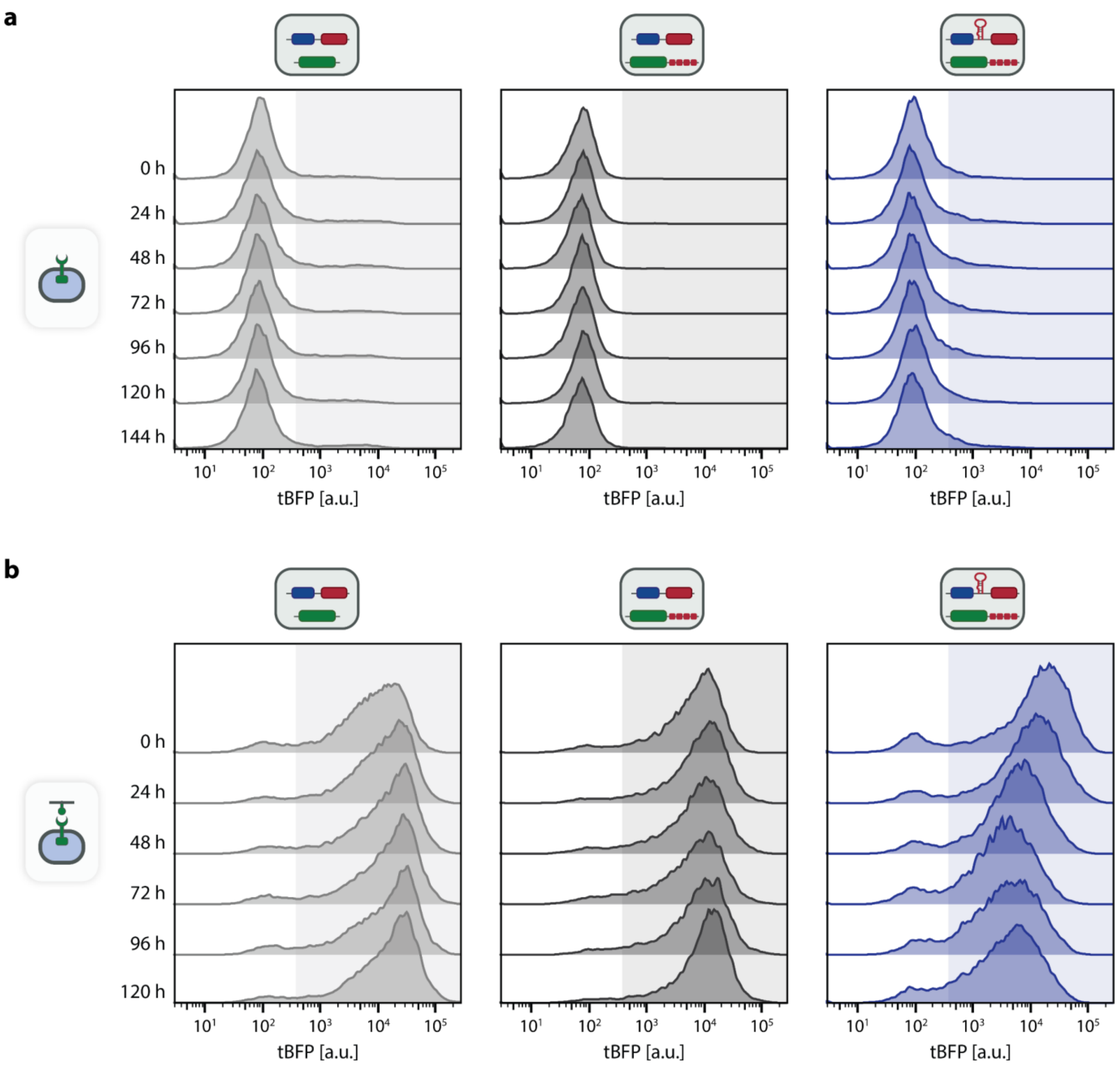


**Supplementary Figure S29: Expression of tBFP in three different *Receiver* cell lines cultured on uncoated and eGFP-coated plates. a-b** Representative flow cytometry histograms of tBFP expression as shown in Extended Data Fig. 3c-3d. Three types of *Receiver* cell lines (left: without feedback loop components, middle: without miR-FF3 gene and with miR-FF3 target sites, right: with miR-FF3 gene and target sites) were seeded at t = 0 h on uncoated **(a)** or eGFP-coated plates **(b)**. Expression of tBFP was quantified over time by flow cytometry analysis, with measurements every 24 hours. Shaded area indicates tBFP-positive cells (threshold defined by HeLa WT control).

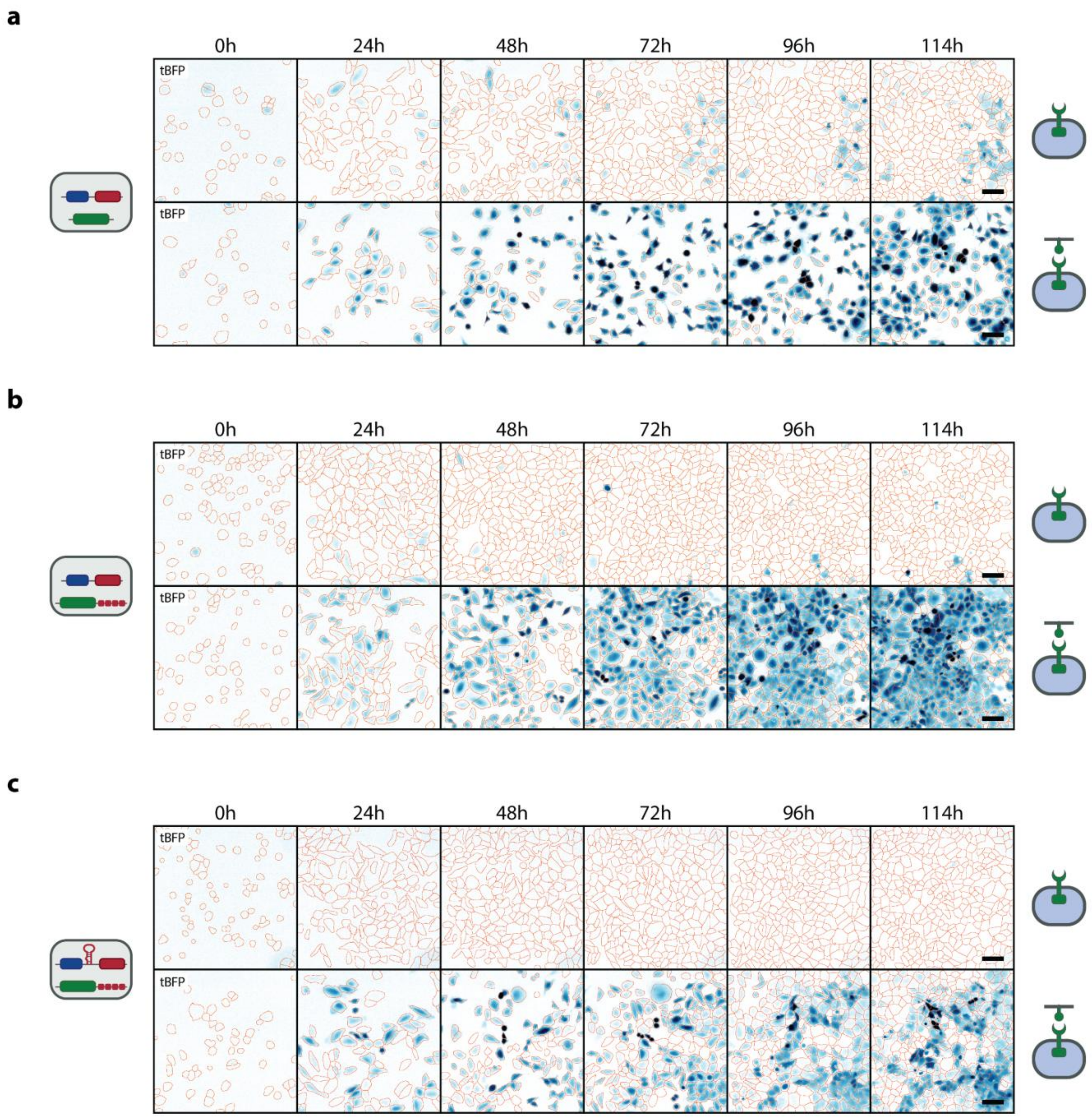


**Supplementary Figure S30: Fluorescence microscopy images showing tBFP expression of three different *Receiver* cell lines cultured on uncoated and eGFP-coated plates. a-c** Representative epifluorescence images showing tBFP expression in *Receiver* cells without feedback loop components **(a)**, without miR-FF3 gene and with miR-FF3 target sites **(b)**, or with miR-FF3 gene and target sites **(c)**, seeded at t = 0 h on uncoated (top) or eGFP-coated (bottom) plates (Methods). tBFP fluorescence is displayed in blue; orange outlines indicate cellular boundaries derived from mCherry-guided image segmentation (Methods). Scale bar: 100 µm.

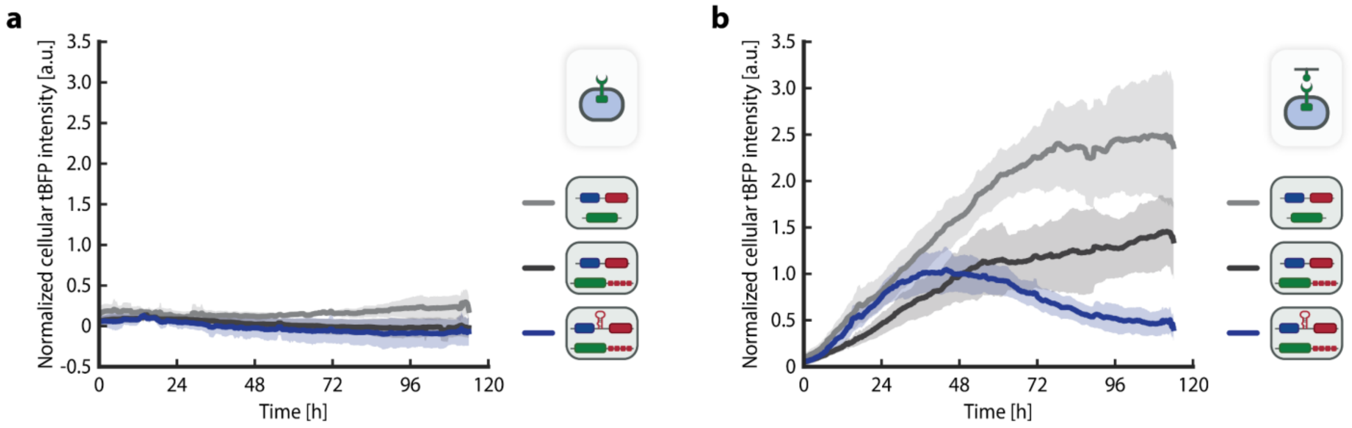


**Supplementary Figure S31: Time-lapse quantification of tBFP expression of three different *Receiver* cell lines cultured on uncoated and eGFP-coated plates. a-b** Time-lapse quantification of tBFP expression of three types of *Receiver* cell lines (light grey: without feedback loop components, dark grey: without miR-FF3 gene and with miR-FF3 target sites, blue: with miR-FF3 gene and target sites). *Receivers* were seeded at t = 0 h on uncoated **(a)** or eGFP-coated plates **(b)** and tBFP expression was quantified using widefield microscopy (Methods). Lines and shaded areas indicate the mean and 95% CI of normalized cellular tBFP intensity, measured every 15 minutes, of n=9 replicates from 3 biologically independent experiments (runs) performed on separate days with 3 wells per condition.

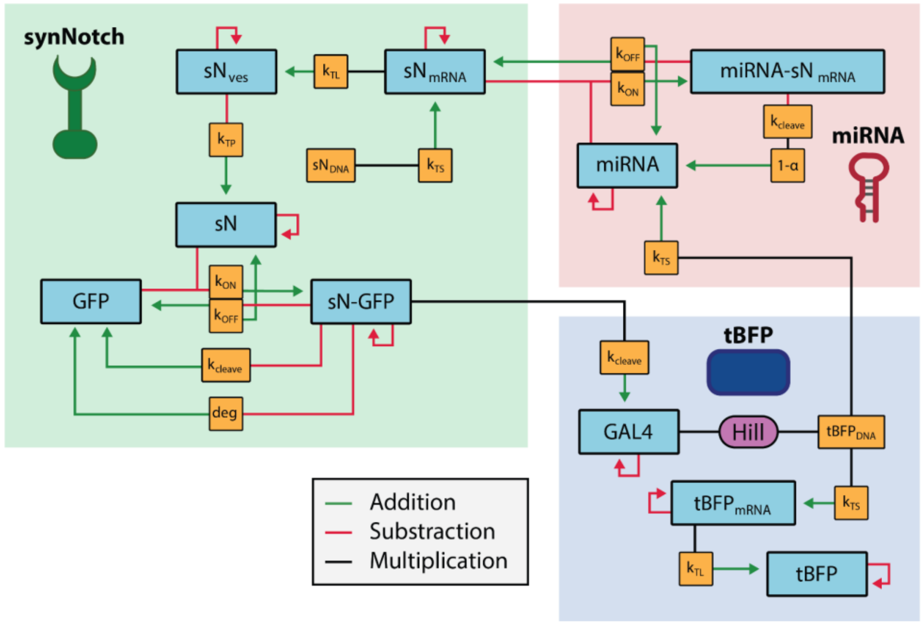


**Supplementary Figure S32: Schematic representation of the ordinary differential equation (ODE) model.** Schematic representation of the reaction network underlying the ODE model (Supplementary Notes). Species (blue boxes) and rate parameters (orange boxes) are organized into three functional modules: synNotch receptor dynamics (green), miRNA feedback (red), and tBFP reporter expression (blue). Relationships within and between the ODEs are depicted using green arrows to indicate additive terms, red arrows to indicate subtractive terms, and black arrows to indicate multiplicative terms. The model captures synNotch production, GFP ligand binding, GAL4-VP64 release and, Hill-kinetic transcriptional activation, and miRNA-mediated mRNA degradation.

**Caption for Supplementary Video S1: Time-lapse fluorescence microscopy of tBFP expression in *Receiver* cells, co-cultured with K562 cells, after precursor transfection.** Representative time-lapse epifluorescence microscopy images showing tBFP expression in *Receiver* cells containing no miR-FF3 target sites (0xTS; left) or four miR-FF3 target sites (4xTS; right), transfected with 0.1 nM miR-FF3 precursor miRNA (top) or scrambled precursor miRNA (bottom), and co-cultured with K562 *Sender* cells. Images were acquired every 3 h for 60 h following initiation of co-culture. tBFP fluorescence is displayed in blue; orange outlines indicate cellular boundaries derived from mCherry-guided image segmentation (Methods). Scale bar: 100 µm.

**Caption for Supplementary Video S2: Time-lapse fluorescence microscopy of tBFP expression in *Receiver* cells with or without the complete synNotch-miRNA feedback loop.** Representative time-lapse epifluorescence microscopy images showing tBFP expression in *Receiver* cells containing miR-FF3 target sites but lacking the miR-FF3 gene (4xTS; left), or *Receiver* cells containing both miR-FF3 target sites and the miR-FF3 gene (4xTS + miR-FF3; right). Cells were seeded at t = 0 h on eGFP-coated plates (top) or on uncoated plates (bottom). Images were acquired every 15 min for 120 h following seeding. tBFP fluorescence is displayed in blue; orange outlines indicate cellular boundaries derived from mCherry-guided image segmentation (Methods). Scale bar: 100 µm.

### Supplementary Notes

#### *Mathematical model of the synNotch-miRNA system*

#### Description of ordinary differential equation (ODE) model

##### Model overview

We developed a mechanistic ordinary differential equation (ODE) model to capture the dynamics of the synNotch-miRNA feedback circuit. The model describes a two-cell system where *Sender* cells express and presenting membrane-bound GFP ligand, and *Receiver* cells express synNotch receptor, tBFP reporter and miRNA. The model architecture comprises three functional modules organized as shown in Supplementary Fig. S32: synNotch receptor dynamics, tBFP reporter expression, and miRNA-mediated feedback regulation. The model employs mass-action kinetics for all reactions, with Hill kinetics only applied to transcriptional activation by the GAL4-VP64 transcription factor.

##### Model equations

$$(\mathbf{1})\ \frac{d[sN_{mRNA}]}{dt} = k_{ts-sN} \cdot sN_{DNA} - deg_{sN_{mRNA}}[sN_{mRNA}] - k_{on-miRNA-sN_{mRNA}}[miRNA][sN_{mRNA}] + k_{off-miRNA-sN_{mRNA}}[miRNA - sN_{mRNA}]$$

$$(\mathbf{2})\ \frac{d[sN_{ves}]}{dt} = k_{tl-sN}[sN_{mRNA}] - k_{tp-sN}[sN_{ves}] - deg_{sN_{ves}}[sN_{ves}]$$

$$(\mathbf{3})\ \frac{d[sN]}{dt} = k_{tp-sN}[sN_{ves}] - deg_{sN}[sN] - k_{on-sN-GFP}[GFP][sN] + k_{off-sN-GFP}[sN - GFP]$$

$$(\mathbf{4})\ \frac{d[GAL4]}{dt} = k_{cleave-sN-GFP}[sN - GFP] - deg_{GAL4}[GAL4]$$

$$(\mathbf{5})\ \frac{d[tBFP_{mRNA}]}{dt} = k_{ts-tBFP_{mRNA}} \cdot tBFP_{DNA} \cdot \frac{[GAL4]^{H_{GAL4}}}{K_{d-GAL4} + [GAL4]^{H_{GAL4}}} - deg_{tBFP_{mRNA}}[tBFP_{mRNA}]$$

$$(\mathbf{6})\ \frac{d[tBFP]}{dt} = k_{tl-tBFP}[tBFP_{mRNA}] - deg_{tBFP}[tBFP]$$

$$(\mathbf{7})\ \frac{d[GFP]}{dt} = deg_{GFP-sN}[sN - GFP] - k_{on-sN-GFP}[GFP][sN] + k_{off-sN-GFP}[sN - GFP] + k_{cleave-sN-GFP}[sN - GFP]$$

$$(\mathbf{8})\ \frac{d[sN - GFP]}{dt} = k_{on-sN-GFP}[GFP][sN] - k_{off-sN-GFP}[sN - GFP] - deg_{GFP-sN}[sN - GFP] - k_{cleave-sN-GFP}[sN - GFP]$$

$$(\mathbf{9})\ \frac{d[miRNA]}{dt} = k_{ts-miRNA} \cdot tBFP_{DNA} \cdot \frac{[GAL4]^{H_{GAL4}}}{K_{d-GAL4} + [GAL4]^{H_{GAL4}}} - k_{on-miRNA-sN_{mRNA}}[miRNA][sN_{mRNA}] + k_{off-miRNA-sN_{mRNA}}[miRNA - sN_{mRNA}] + (1 - \alpha_{miRNA-loss-rate}) \cdot k_{cleave-miRNA-sN_{mRNA}}[miRNA - sN_{mRNA}] - deg_{miRNA}[miRNA]$$

$$(\mathbf{10})\ \frac{d[miRNA - sN_{mRNA}]}{dt} = k_{on-miRNA-sN_{mRNA}}[miRNA][sN_{mRNA}] - k_{off-miRNA-sN_{mRNA}}[miRNA - sN_{mRNA}] - k_{cleave-miRNA-sN_{mRNA}}[miRNA - sN_{mRNA}]$$

**Description of model equations**

The model describes the dynamics of synNotch receptor expression, activation, and downstream transcriptional output, as well as miRNA-mediated repression of synNotch mRNA. The complete synNotch-miRNA system consists of 10 coupled ODEs describing the temporal dynamics of the involved molecular species (**Equations 1-10**).

The circuit begins with constitutive transcription of synNotch mRNA from genomic DNA (**Equation 1**). This mRNA can exist in two states: free and available for translation, or bound to miRNA in a regulatory complex. The free synNotch mRNA concentration increases through transcription and through dissociation of miRNA-mRNA complexes (which returns mRNA to the translatable pool), while it decreases through natural degradation and through binding to complementary mRNA, forming the miRNA-mRNA complex. Only this free mRNA serves as template for protein synthesis.

The synNotch mRNA is translated into synNotch receptor protein that initially resides in intracellular vesicles (**Equation 2**). This vesicular protein pool increases through translation and decreases through both transport to the cell membrane and natural degradation while in the vesicle compartment. The explicit modeling of this vesicular intermediate captures the temporal delay between mRNA translation and functional receptor availability at the cell surface.

Upon transport to the membrane, synNotch receptors become available for ligand binding (**Equation 3**). The membrane receptor population increases through arrival of newly synthesized protein from vesicles and through unbinding from the GFP ligand, while it decreases through natural membrane receptor degradation and through binding to GFP ligand presented on *Sender* cells.

The GFP ligand exists in multiple states within the system. Free GFP ligand (**Equation 7**) increases through three mechanisms: release from degraded receptor-ligand complexes, unbinding from intact complexes, and release during proteolytic cleavage of the synNotch-GFP complex. Free GFP ligand decreases through binding to synNotch receptor. The total GFP concentration (free plus bound) remains constant throughout simulations, reflecting the assumption that ligand is not internalized or degraded during receptor engagement.

When receptor and ligand bind, they form the synNotch-GFP complex (**Equation 8**), which represents the activated state of the receptor. This complex population increases through binding of free receptor to free ligand and decreases through three competing pathways: dissociation back to free components, natural degradation of the intact complex, and proteolytic cleavage. The proteolytic cleavage reaction constitutes the signal transduction event that links extracellular ligand binding to intracellular transcriptional activation.

Proteolytic cleavage of the synNotch-GFP complex releases the GAL4-VP64 transcription factor into the cytoplasm (**Equation 4**). The transcription factor concentration increases through receptor cleavage and decreases through natural protein degradation. This transcription factor drives expression of the downstream outputs by binding to GAL4-responsive promoters.

GAL4-VP64 activates transcription of two genes under control of the GAL4-UAS promoter: the tBFP reporter (**Equation 5**) and the regulatory miRNA (**Equation 9**). Both transcription reactions are modeled using identical Hill kinetics, as both genes share the same promoter and thus respond equivalently to transcription factor concentration. The Hill coefficient captures cooperativity in transcription factor binding, while the dissociation constant reflects binding affinity. For tBFP mRNA, transcription is balanced by natural mRNA degradation.

The tBFP mRNA is translated into fluorescent protein (**Equation 6**), which serves as the experimentally observable readout of circuit activity. Reporter protein concentration increases through translation and decreases through natural protein degradation.

The miRNA binding to complementary synNotch mRNA closes the negative feedback loop (**Equation 9**). Free miRNA, representing mature miRNA loaded into the RNA-induced silencing complex (RISC; not explicitly modeled), increases through three sources: transcription driven by GAL4-VP64, unbinding from synNotch mRNA targets, and recycling after catalyzing target mRNA cleavage. The recycling mechanism is governed by the parameter α, which determines the fraction of miRNA molecules that are degraded versus released during each mRNA cleavage event. Free miRNA decreases through binding to target mRNA and through natural miRNA degradation.

When miRNA binds to complementary synNotch mRNA, it forms a miRNA-mRNA complex (**Equation 10**). The concentration of this complex increases through binding of free miRNA to free synNotch mRNA and decreases through two pathways: dissociation that releases intact mRNA, or irreversible cleavage that degrades the mRNA. The cleavage reaction removes synNotch mRNA from the system, thereby reducing synNotch receptor expression.

##### Base synNotch model (without miRNA feedback)

The base synNotch model without miRNA feedback regulation consists of **Equations 1-8**, with all miRNA-related parameters set to zero. Specifically, this means setting all rate constants involving miRNA to zero ($k_{on-miRNA-sNmRNA} = 0$, $k_{off-miRNA-sNmRNA} = 0$, $k_{cleave-miRNA-sNmRNA} = 0$, $k_{ts-miRNA} = 0$, and $\deg_{miRNA} = 0$), and removing Equations 9 and 10 from the system. In this configuration, synNotch mRNA dynamics (**Equation 1**) reduce to simple constitutive transcription balanced by natural degradation, with no miRNA-mediated regulation.

##### Initial conditions

Initial conditions for simulations were determined by first running the system to steady state in the absence of GFP ligand. Specifically, the system was simulated for 120 hours with no GFP present until all species reached steady state concentrations. Using the default parameters, this yielded basal values of 0.25 nM for synNotch mRNA, 2.5 nM for synNotch protein (vesicle), and 16.5 nM for membrane-localized synNotch receptor. To simulate ligand-induced activation, GFP ligand was introduced at t = 0 h with a concentration of 16.5 nM and maintained constant throughout the simulation, representing constitutive GFP presentation by *Sender* cells. This GFP concentration was chosen to match the steady state synNotch receptor concentration, assuming similar production and degradation rates for membrane-bound GFP and synNotch receptor.

**Model simulation**

The ODE model was implemented in Python and numerically integrated using the 'solve_ivp' function, using the 'Radau' method, from the SciPy library (Methods). Simulations were run over 120-hour time courses with 500 evaluation points.

### ODE model parameter values and derivations

#### Parameter values

| Parameter | Value | Unit | Explanation |
|---|---|---|---|
| $k_{ts-sN}$ | 3.6 | $hr^{-1}$ | Transcription rate of synNotch mRNA from synNotch DNA[6]. |
| $sN_{DNA}$ | 0.0214 | $nM$ | Concentration of synNotch DNA in the nucleus of *Receiver* cell. Value is derived below. |
| $k_{tl-sN}$ | 36 | $hr^{-1}$ | Translation rate of synNotch mRNA[6]. |
| $k_{tp-sN}$ | 3.48 | $hr^{-1}$ | Trafficking rate of synNotch to the cell membrane[7]. |
| $k_{ts-tBFP_{mRNA}}$ | 3.6 | $hr^{-1}$ | Maximum transcription rate of tBFP mRNA when promoter is fully saturated by GAL4-VP64[6]. |
| $K_{d-GAL4}$ | 0.005 | $nM$ | Dissociation constant of GAL4-VP64 binding to GAL4-UAS promoter[8]. |
| $H_{GAL4}$ | 2 | – | Hill coefficient for GAL4-VP64 binding to GAL4-UAS promoter[8]. |
| $tBFP_{DNA}$ | 2.24 | $nM$ | Concentration of tBFP DNA in the nucleus of *Receiver* cell. Value is derived below. |
| $k_{tl-tBFP}$ | 36 | $hr^{-1}$ | Translation rate of tBFP mRNA[6]. |
| $k_{on-sN-GFP}$ | 11.16 | $nM^{-1}hr^{-1}$ | Association rate of GFP ligand binding to the LaG17 extracellular domain of synNotch[9]. |
| $k_{off-sN-GFP}$ | 504 | $hr^{-1}$ | Dissociation rate of synNotch-GFP complex[9]. |
| $k_{cleave-sN-GFP}$ | 0.03 | $hr^{-1}$ | Cleavage rate of synNotch in the synNotch-GFP complex, releasing GAL4-VP64 and GFP. Value is derived below. |
| $deg_{sN_{mRNA}}$ | 0.347 | $hr^{-1}$ | Degradation rate of synNotch mRNA. SynNotch is based on endogenous Notch1[1], which has a half-life of approximately 2 hours[10]. |
| $deg_{sN_{ves}}$ | 0.018 | $hr^{-1}$ | Degradation rate of synNotch in membrane transport vesicle[6]. |
| $deg_{sN}$ | 0.48 | $hr^{-1}$ | Degradation rate of synNotch on *Receiver* cell membrane[7]. |
| $deg_{GAL4}$ | 0.173 | $hr^{-1}$ | Degradation rate of GAL4-VP64, based on the half-life of a GAL4 chimeric protein[11]. |
| $deg_{tBFP_{mRNA}}$ | 0.347 | $hr^{-1}$ | Degradation rate of tBFP mRNA, same value as degradation rate of synNotch mRNA[10]. |
| $deg_{tBFP}$ | 0.1195 | $hr^{-1}$ | Degradation rate of tBFP. Value is derived below. |
| $deg_{GFP-sN}$ | 0.48 | $hr^{-1}$ | Degradation rate of the synNotch-GFP complex on the membrane. Rate is assumed to be identical to degradation rate of free synNotch on the membrane[7]. |
| $k_{ts-miRNA}$ | 3.6 | $hr^{-1}$ | Transcription rate of miRNA, the rate is assumed to be identical to the transcription rate of tBFP mRNA[6]. |
| $k_{on-miRNA-sN_{mRNA}}$ | 0.120 | $nM^{-1}hr^{-1}$ | Association rate of miRNA (RISC) binding to synNotch mRNA. Value is derived below. |
| $k_{off-miRNA-sN_{mRNA}}$ | 10.8 | $hr^{-1}$ | Dissociation rate of the miRNA-mRNA complex. The rate is based on AGO2-RISC guided by let7a miRNA[12]. |
| $k_{cleave-miRNA-sN_{mRNA}}$ | 237.6 | $hr^{-1}$ | Cleavage rate of synNotch mRNA by RISC[12]. |
| $\alpha_{miRNA-loss-rate}$ | 0.10 | – | The miRNA loss rate is the probability that a miRNA is recycled after having interacted with one of its complementary mRNA targets. Value is derived below. |
| $deg_{miRNA}$ | 0.046 | $hr^{-1}$ | Degradation rate of miRNA. miRNA used in this study is based on miR-30, which has a half-life of ≈15 hours[14]. |

**Derivation of synNotch and tBFP DNA concentration**

SynNotch DNA ($sN_{DNA}$) is derived from solving the ODE model **Equations 1, 2 and 3** without presence of ligand ([GFP] = 0) at a steady state (t → inf):

$$[sN_{mRNA}] = \frac{k_{ts-sN} sN_{DNA}}{deg_{sN_{mRNA}}},$$

$$[sN_{ves}] = \frac{k_{tl-sN}[sN_{mRNA}]}{k_{tp-sN} + deg_{sN_{ves}}},$$

$$[sN] = \frac{k_{tp-sN}[sN_{ves}]}{deg_{sN}}.$$

Solving for $sN_{DNA}$, we reach the expression:

$$sN_{DNA} = \frac{deg_{sN}}{k_{tp-sN}} \frac{k_{tp-sN} + deg_{sN_{ves}}}{k_{tl-sN}} \frac{deg_{sN_{mRNA}}}{k_{ts-sN}} [sN].$$

SynNotch DNA concentration is calculated based on the expected receptor density on the *Receiver* cells. We assume that each gene copy yields a mature receptor. In addition, we assume that it traffics identically to EGF since Notch has several EGF repeats[15]. To compute the expected receptor density for synNotch, the concentration of EGF receptors on a cell was taken as representative, which is $2.5*10^4$ receptors/cell[16]. The volume of an HeLa cell is approximately 2500 $\mu m^3$ when adhered to a surface[17]. When combining the receptor count per cell and the cell volume we get an approximate receptor density for the synNotch receptors of $[sN] = 16.5\, nM$. Substituting this value into the previous equation yields $sN_{DNA} = 0.0214\, nM$.

tBFP DNA ($tBFP_{DNA}$) is derived in an identical way from ODE model **Equations 5 and 6**:

$$tBFP_{DNA} = \frac{deg_{tBFP}}{k_{tl-tBFP}} \frac{deg_{tBFP_{mRNA}}}{k_{ts-tBFP_{mRNA}}} [tBFP].$$

tBFP DNA concentration is calculated based on the expected concentration of tBFP protein $[tBFP]$ at steady state. We assume that when tBFP is fully expressed with an excess of GAL4-VP64 transcription factor, the concentration of the tBFP protein will approach a similar value as that of the GFP protein concentration (7 $\mu M$) when constitutively expressed in HeLa cells[18]. Substituting this value into the previous equation yields us $tBFP_{DNA} = 2.2397\, nM$.

**Derivation of tBFP degradation rate**

We assume that tBFP has a similar degradation rate as GFP. The degradation rate of the tBFP protein has been estimated based on *Figure S2b* from Morsut et al.[1]. This figure shows the downstream activation upon synNotch receptor activation and the observed timescales of this activation. In the second part of the figure, *Sender* cells are removed, so we assume that the concentration of the GFP reporter decreases with the protein degradation rate corresponding to GFP. The half-life of GFP from this figure is between 5.8 and 7.5 hours. Furthermore, Corish and Tyler-Smith[19] made GFP with half-lives of 9.8, 5.8 and 5.5 hours, which is in a similar order of magnitude as the estimated GFP half-life in Morsut et al.[1]. Therefore, for this model a tBFP half-life of 5.8 hours was used, as it agrees with both sources, resulting in a degradation rate of $deg_{tBFP} = 0.1195\, hr^{-1}$.

**Derivation of miRNA loss rate and miRNA-mRNA binding rates**

The miRNA loss rate α is calculated using the following equation from Yuan et al.[13]: $\theta_i = \frac{k_s}{\alpha_i g_{R_i}}$, where $\theta$ is an experimentally fitted constant (103.98), $k_s$ is the rate of miRNA production, and $g_{R_i}$ is the rate of natural miRNA degradation. Rewriting and solving the equation we yield:

$$\alpha_{miRNA-loss-rate} = \frac{k_{ts-miRNA}}{\theta deg_{miRNA}} = 0.1\ .$$

In a similar fashion, the miRNA-mRNA binding rate is determined based on the equations of Yuan et al.[13]: $\lambda_i = \frac{g_s}{\alpha_i k_{i+}}\left(\frac{k_{i-}}{g_i} + 1\right)$, where the constant λ was experimentally determined to be 4.01. $g_s$ is the degradation rate of miRNA, $\alpha_i$ is the loss rate of miRNA, $k_{i+}$ is the miRNA-mRNA binding rate, $k_{i-}$ is the miRNA-mRNA unbinding rate and $g_i$ is the cleavage rate of miRNA-bound mRNA. Rewriting and solving the equation we yield:

$$k_{on-miRNA-sN_{mRNA}} = \frac{deg_{miRNA}}{\alpha_{miRNA-loss-rate}\lambda}\left(\frac{k_{off-miRNA-sN_{mRNA}}}{k_{cleave-miRNA-sN_{mRNA}}} + 1\right) = 0.120 nM^{-1}hr^{-1}$$

**Derivation of synNotch-GFP complex cleavage rate**

Gonzalez-Perez et al. investigated a custom Notch ligand with a Notch-ligand dissociation constant between 24 and 54 nM[20], which is close to the dissociation constant of the LaG17 nanobody utilized in our system (50 nM)[9]. Furthermore, Gonzalez-Perez et al. similarly utilize a GAL4 intracellular Notch domain and a reporter under GAL4 transcription factor control. Gonzalez-Perez et al. found that the half-maximum response of their reporter was reached for a concentration of 1.7 nM of the Delta$^{MAX}$ ligand[20]. Therefore, the cleavage rate of the synNotch-GFP complex was set such that at a concentration of 1.7 nM of GFP ligand, the tBFP reporter would reach its half-maximal steady state response. The cleavage rate of the synNotch-GFP complex was subsequently set to $k_{cleave-sN-GFP} = 0.03 hr^{-1}$.

### List of abbreviations

| | |
|---|---|
| **AmpR** | Ampicillin Resistance |
| **cPPT/CTS** | central PolyPurine Tract/Central Termination Sequence |
| **eGFP** | enhanced Green Fluorescent Protein |
| **FACS** | Fluorescence-Activated Cell Sorting |
| **FSC** | Forward Scatter |
| **hsa** | homo sapiens |
| **LTR** | Long Terminal Repeat |
| **miRNA** | microRNA |
| **ORI** | Origin of Replication |
| **PGK** | Phosphoglycerate Kinase |
| **qPCR** | quantitative Polymerase Chain Reaction |
| **RISC** | RNA-Induced Silencing Complex |
| **RRE** | Rev Response Element |
| **SEAP** | Secreted Embryonic Alkaline Phosphatase |
| **SFFV** | Spleen Focus-Forming Virus |
| **SSC** | Side Scatter |
| **SV40** | Simian Vacuolating Virus 40 |
| **tBFP** | tag Blue Fluorescent Protein |
| **TS** | Target Site |
| **UAS** | Upstream Activating Sequence |
| **WPRE** | Woodchuck Hepatitis Virus Post-transcriptional Regulatory Element |
| **WT** | Wild-Type |
